# AIDE: annotation-assisted isoform discovery with high precision

**DOI:** 10.1101/437350

**Authors:** Wei Vivian Li, Shan Li, Xin Tong, Ling Deng, Hubing Shi, Jingyi Jessica Li

## Abstract

Genome-wide accurate identification and quantification of full-length mRNA isoforms is crucial for investigating transcriptional and post-transcriptional regulatory mechanisms of biological phenomena. Despite continuing efforts in developing effective computational tools to identify or assemble full-length mRNA isoforms from second-generation RNA-seq data, it remains a challenge to accurately identify mRNA isoforms from short sequence reads due to the substantial information loss in RNA-seq experiments. Here we introduce a novel statistical method, AIDE (Annotation-assisted Isoform DiscovEry), the first approach that directly controls false isoform discoveries by implementing the testing-based model selection principle. Solving the isoform discovery problem in a stepwise and conservative manner, AIDE prioritizes the annotated isoforms and precisely identifies novel isoforms whose addition significantly improves the explanation of observed RNA-seq reads. We evaluate the performance of AIDE based on multiple simulated and real RNA-seq datasets followed by a PCR-Sanger sequencing validation. Our results show that AIDE effectively leverages the annotation information to compensate the information loss due to short read lengths. AIDE achieves the highest precision in isoform discovery and the lowest error rates in isoform abundance estimation, compared with three state-of-the-art methods Cufflinks, SLIDE, and StringTie. As a robust bioinformatics tool for transcriptome analysis, AIDE will enable researchers to discover novel transcripts with high confidence.

## Introduction

A transcriptome refers to the entire set of RNA molecules in a biological sample. Alternative splicing, a post-transcriptional process during which particular exons of a gene may be included into or excluded from a mature messenger RNA (mRNA) isoform transcribed from that gene, is a key contributor to the diversity of eukaryotic transcriptomes (Ghigna et al. 2008). Alternative splicing is a prevalent phenomenon in multicellular organisms, and it affects approximately 90%-95% of genes in mammals (Hooper 2014). Understanding the diversity of eukaryotic transcriptomes is essential to interpreting gene functions and activities under different biological conditions (Adams 2008). In transcriptome analysis, a key task is to accurately identify the set of truly expressed isoforms and estimate their abundance levels under a specific biological condition, because the information on isoform composition is critical to understanding the isoform-level dynamics of RNA contents in different cells, tissues, and developmental stages. Abnormal splicing events have been known to cause many genetic disorders (Wang and Cooper 2007), such as retinitis pigmentosa (Mordes et al. 2006) and spinal muscular atrophy (Singh and Singh 2011). Accurate isoform identification and quantification will shed light on gene regulatory mechanisms of genetic diseases, thus assisting biomedical researchers in designing targeted therapies for diseases.

The identification of truly expressed isoforms is an indispensable step preceding accurate isoform quantification. However, compared with the quantification task, isoform discovery is an inherently more challenging problem both theoretically and computationally. The reasons behind this challenge are threefold. First, second-generation RNA-seq reads are too short compared with full-length mRNA isoforms. RNA-seq reads are typically no longer than 300 base pairs (bp) in Illumina sequencing (Chhangawala et al. 2015), while more than 95% of human isoforms are longer than 300 bp, with a mean length of 1, 712 bp (the GENCODE annotation, Release 24) (Harrow et al. 2012). Hence, RNA-seq reads are short fragments of full-length isoforms. Since most isoforms of the same gene share some overlapping regions, many RNA-seq reads do not unequivocally map to a unique isoform. As a result, isoform origins of those reads are ambiguous and need to be inferred from a huge pool of candidate isoforms. Another consequence of short reads is that “junction reads” spanning more than one exon-exon junctions are underrepresented in second-generation RNA-seq data, due to the difficulty of mapping junction reads (every read needs to be split into at least two segments and has all the segments mapped to different exons in the reference genome). The underrepresentation of those junction reads further increases the difficulty of discovering full-length RNA isoforms accurately. Second, the number of candidate isoforms increases exponentially with the number of exons. Hence, computational efficiency becomes an inevitable factor that every method must account for, and an effective isoform screening step is often needed to achieve accurate isoform discovery (Ye and Li 2016). Third, it is a known biological phenomenon that often only a small number of isoforms are truly expressed under one biological condition. Given the huge number of candidate isoforms, how isoform discovery methods balance the parsimony and accuracy of their discovered isoforms becomes a critical and meanwhile difficult issue (Canzar et al. 2016; Mezlini et al. 2013). For more comprehensive discussion and comparison of existing isoform discovery methods, readers can refer to (Steijger et al. 2013) and (Li and Li 2018).

Over the past decade, computational researchers have developed multiple state-of-the-art isoform discovery methods to tackle one or more of the challenges mentioned above. The two earliest annotation-free methods are Cufflinks (Trapnell et al. 2010) and Scripture (Guttman et al. 2010), which can assemble mRNA isoforms solely from RNA-seq data without using annotations of known isoforms. Both methods use graph-based approaches, but they differ in how they construct graphs and then parse a graph into isoforms. Scripture first constructs a connectivity graph with nodes as genomic positions and edges determined by junction reads. It then scans the graph with fixed-sized windows, scores each path for significance, connects the significant paths into candidate isoforms, and finally refines the isoforms using paired-end reads. Cufflinks constructs an overlap graph of mapped reads, and it puts a directed edge based on the genome orientation between two compatible reads that could arise from the same isoform. It then finds a minimal set of paths that cover all the fragments in the overlap graph. A more recent method StringTie also uses the graph idea (Pertea et al. 2015). It first creates a splice graph with read clusters as nodes to identify isoforms, and it then constructs a flow network to estimate the expression levels of isoforms using a maximum flow algorithm.

Another suite of methods utilize different statistical and computational tools and regularization methods to tackle the problems of isoform discovery and abundance estimation. IsoLasso (Li et al. 2011b), SLIDE (Li et al. 2011a), and CIDANE (Canzar et al. 2016) all build linear models, where read counts are summarized as the response variable, and isoform abundances are treated as parameters to be estimated. IsoLasso starts from enumerating valid isoforms and then uses the LASSO algorithm (Tibshirani 1996) to achieve parsimony in its discovered isoforms. SLIDE incorporates the information of gene and exon boundaries in annotations to enumerate candidate isoforms, and it uses a modified LASSO procedure to select isoforms. CIDANE also uses the LASSO regression in its first phase, and then it employs a delayed column generation technique in its second phase to check if new isoforms should be added to improve the solution. Another method iReckon takes a different approach and tackles the isoform discovery problem via maximum likelihood estimation (Mezlini et al. 2013). It first constructs all the candidate isoforms supported by RNA-seq data, and then it utilizes a regularized expectation-maximization (EM) algorithm (Dempster et al. 1977) to reduce the number of expressed isoforms and avoid over-fitting.

Aside from the intrinsic difficulty of isoform identification due to the short read lengths and the huge number of candidate isoforms, the excess biases in RNA-seq experiments further afflict the isoform discovery problem. Ideally, RNA-seq reads are expected to be uniformly distributed within each isoform. However, the observed distribution of RNA-seq reads significantly violates the uniformity assumption due to multiple sources of biases. The most commonly acknowledged bias source is the different levels of guanine-cytosine (GC) contents in different regions of an isoform. The GC content bias was first investigated by Dohm et al. (Dohm et al. 2008), and a significantly positive correlation was observed between the read coverage and the GC contents. Another work later showed that the effects of GC contents tend to be sample-specific (Risso et al. 2011). Another major bias source is the positional bias, which causes the uneven read coverage at different relative positions within an isoform. As a result of the positional bias, reads are more likely to be generated from certain regions of an isoform, depending on experimental protocols, e.g., whether cDNA fragmentation or RNA fragmentation is used (Li et al. 2010; Frazee et al. 2015). Failing to correct these biases will likely lead to high false discovery rates in isoform discovery and unreliable statistical results in downstream analyses, such as differential isoform expression analysis (Patro et al. 2017). Current computational methods account for the non-uniformity of reads using three main approaches: to adjust read counts summarized in defined genomic regions to offset the non-uniformity bias (Li et al. 2010; Zheng et al. 2011; Roberts et al. 2011b), to assign a weight to each single read to adjust for bias (Hansen et al. 2010), and to incorporate the bias as a model parameter in likelihood-based methods (Bohnert and Rätsch 2010; Roberts et al. 2011b; Jiang and Salzman 2015; Love et al. 2016).

Despite continuous efforts the bioinformatics community has spent on developing effective computational methods to identify full-length isoforms from second-generation RNA-seq data, the existing methods still suffer from low accuracy for genes with complex splicing structures (Steijger et al. 2013; Conesa et al. 2016). A comprehensive assessment has shown that methods achieving good accuracy in identifying isoforms in *Drosophila melanogaster* (34, 776 annotated isoforms) and *Caenorhabditis elegans* (61, 109 annotated isoforms) fail to maintain good performance in *Homo sapiens* (200, 310 annotated isoforms) (Steijger et al. 2013; Zerbino et al. 2017). Although it is generally believed that deeper sequencing will lead to better isoform discovery results, the improvement is not significant in *H. sapiens*, compared with *D. melanogaster* and *C. elegans*, due to the complex splicing structures of human genes (Mortazavi et al. 2008; Steijger et al. 2013). Moreover, despite increasing accuracy of identified isoforms evaluated at the nucleotide level and the exon level, it remains difficult to improve the isoform-level performance. In other words, even when all sub-isoform elements (i.e., short components of transcribed regions such as exons) are correctly identified, accurate assembly of these elements into full length isoforms remains a big challenge.

Motivated by the observed low accuracy in identifying full-length isoforms solely from RNA-seq data, researchers have considered leveraging information from reference annotations (e.g., Ensembl (Zerbino et al. 2017), GENCODE (Harrow et al. 2012), and UCSC Genome Browser (Rosenbloom et al. 2015)) to aid isoform discovery. Existing efforts include two approaches. In the first approach, methods extract the coordinates of gene and exon boundaries, i.e., known splicing sites, from annotations and then assemble novel isoforms based on the exons or subexons (the regions between adjacent known splicing sites) of every gene (Canzar et al. 2016; Pertea et al. 2015; Li et al. 2011a). In the second approach, methods directly incorporate all the isoforms in annotations by simulating faux-reads from the annotated isoforms (Roberts et al. 2011a). However, the above two approaches have strong limitations in their use of annotations. The first approach does not fully use annotation information because it neglects the splicing patterns of annotated isoforms, and these patterns could assist learning the relationship between short reads and full-length isoforms. The second approach is unable to filter out non-expressed annotated isoforms because researchers lack prior knowledge on which annotated isoforms are expressed in an RNA-seq sample; hence, its addition of unnecessary faux-reads will bias the isoform discovery results and lose control of the false discovery rate.

Here we propose a more statistically principled approach, AIDE (**A**nnotation-assisted **I**soform **D**iscov**E**ry), to leverage annotation information in a more advanced manner to increase the precision and robustness of isoform discovery. Our approach is rooted in statistical model selection, which takes a conservative perspective to search for the smallest model that fits the data well, after adjusting for the model complexity. In our context, a model corresponds to a set of candidate isoforms (including annotated and non-annotated ones), and a more complex model contains more isoforms. Our rationale is that a robust and conservative computational method should only consider novel isoforms as credible if adding them would significantly better explain the observed RNA-seq reads than using only the annotated isoforms. AIDE differs from many existing approaches in that it does not aim to find all seemingly novel isoforms. It enables controlling false discoveries in isoform identification by employing a statistical testing procedure, which ensures that the discovered isoforms make statistically significant contributions to explaining the observed RNA-seq reads. Specifically, AIDE learns gene and exon boundaries from annotations and also selectively borrows information from the annotated isoform structures using a stepwise likelihood-based selection approach. Instead of fully relying on the annotation, AIDE is capable of identifying non-expressed annotated isoforms and removing them from the identified isoform set. Moreover, AIDE simultaneously estimates the abundance of the identified isoforms in the process of isoform reconstruction.

## Results

The AIDE method utilizes the likelihood ratio test to identify isoforms via a stepwise selection procedure, which gives priority to the annotated isoforms and selectively borrows information from their structures. The stepwise selection consists of both forward and backward steps. Each forward step finds an isoform whose addition contributes the most to the explanation of RNA-seq reads given the currently identified isoforms. Each backward step rectifies the identified isoform set by removing the isoform with the most trivial contribution given other identified isoforms. AIDE achieves simultaenous isoform discovery and abundance estimation based on a carefully constructed probabilistic model of RNA-seq read generation (Methods; Figure 1). We first used a comprehensive transcriptome-wide study to evaluate the performance of AIDE and three other widely used methods (Cufflinks, SLIDE, and StringTie) provided with varying read coverages and annotations of different quality. Second, we assessed the precision and recall of these four methods on real human and mouse RNA-seq datasets. In addition, we compared the four methods based on isoforms identified by the long-read sequencing technologies. Third, we validated the performance of AIDE using polymerase chain reaction (PCR) experiments and an additional comparison with the Nanostring data. We finally used a simulation study to demonstrate the necessity and superiority of stepwise selection in AIDE. In all these studies, AIDE demonstrated its advantages in achieving the highest precision in isoform discovery and the best accuracy in
isoform quantification among the four methods.

**Figure 1:**
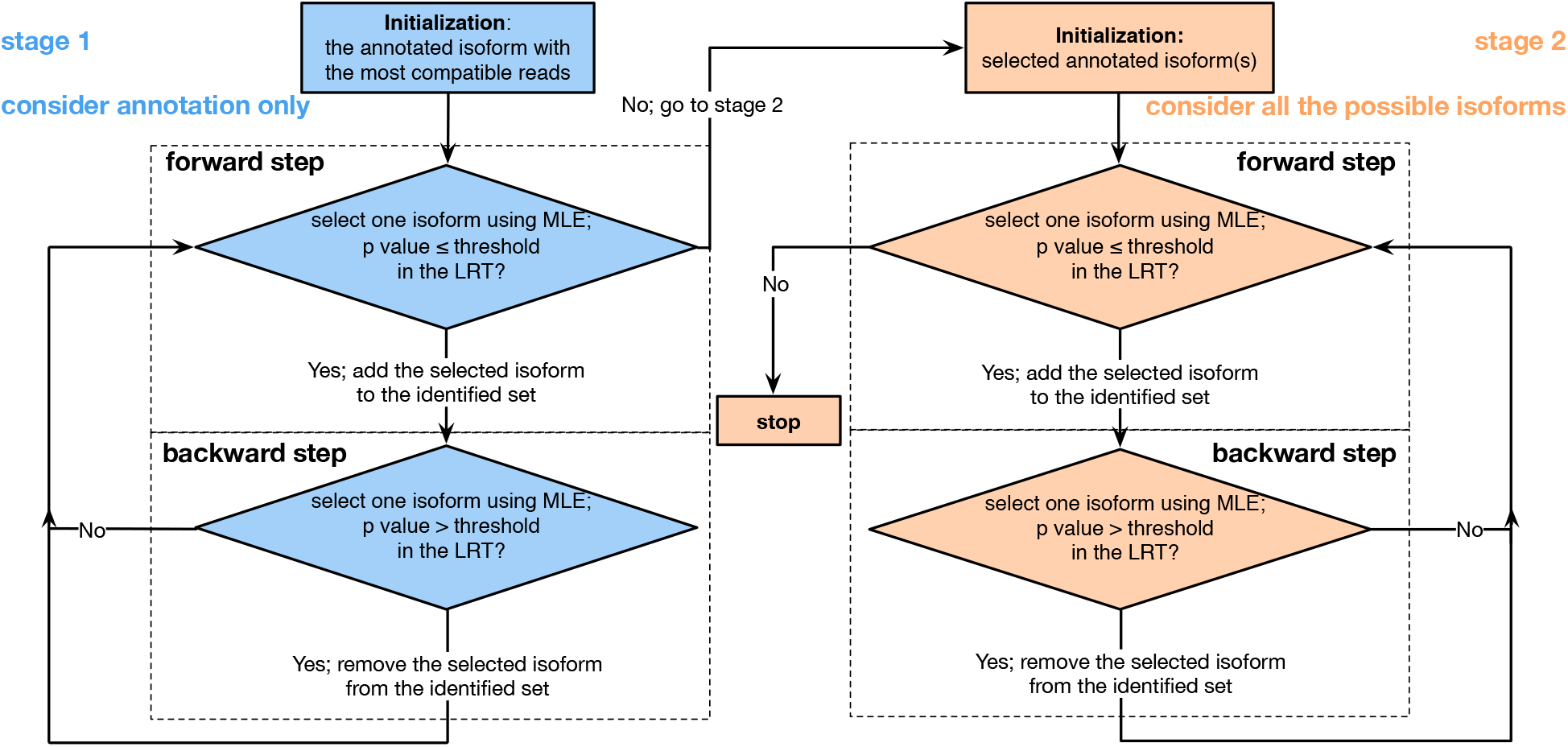
Workflow of the stepwise selection in the AIDE method. Stage 1 starts with a single annotated isoform compatible with the most reads, and all the other annotated isoforms are considered as candidate isoforms. Stage 2 starts with the annotated isoforms selected in stage 1, and all the possible isoforms, including the unselected annotated isoforms, are considered as candidate isoforms. In the forward step in both stages, AIDE identifies the isoform that mostly increases the likelihood, and it uses the likelihood ratio test (LRT) to decide whether this increase is statistically significant. If significant, AIDE adds this isoform to its identified isoform set; otherwise, AIDE keeps its identified set and terminates the current stage. In the backward step in both stages, AIDE finds the isoform in its identified set such that the removal of this isoform decreases the likelihood the least, and it uses the LRT to decide whether this decrease is statistically significant. If not significant, AIDE removes this isoform from its identified set; otherwise, AIDE keeps the identified set. After the backward step, AIDE returns to the forward step. AIDE stops when the forward step in Stage 2 no longer adds a candidate isoform to the identified set.

### AIDE outperforms state-of-the-art methods on simulated data

We compared AIDE with three other state-of-the-art isoform discovery methods, Cufflinks (Trapnell et al. 2010), StringTie (Pertea et al. 2015), and SLIDE (Li et al. 2011a), in a simulation setting that well mimicked real RNA-seq data analysis. The four methods tackle the isoform discovery task from different perspectives. Cufflinks assembles isoforms by constructing an overlap graph and searching for isoforms as sparse paths in the graph. SLIDE utilizes a regularized linear model, and demonstrated precise results in large-scale comparisons (Steijger et al. 2013). StringTie uses a network-based algorithm, and achieves the best computational efficiency and memory usage among the existing methods (Pertea et al. 2015). Unlike all these three methods, our proposed method AIDE, built upon likelihood ratio tests and stepwise selection, converts the isoform discovery problem into a statistical variable selection problem.

To conduct a fair assessment, we simulated RNA-seq datasets using the R (R Core Team 2018) package `polyester` (Frazee et al. 2015), which uses both built-in models and real RNA-seq data to generate synthetic RNA-seq data that exhibit similar properties to those of real data. We simulated eight human RNA-seq datasets with different read coverages (10×, 20×, …, 80×) and pre-determined isoform fractions (see Methods). An “*n*×” coverage means that an exonic genomic locus is covered by *n* reads on average. We compared the accuracy of the four methods supplied with annotations of different quality. In real scenarios, annotations contain both expressed (true) and non-expressed (false) isoforms in a specific RNA-seq sample. Annotations might miss some expressed isoforms in that RNA-seq sample because alternative splicing is known to be condition-specific and widely diverse across different conditions. Therefore, it is critical to evaluate the extent to which different methods rely on the accuracy of annotations, specifically the annotation purity and the annotation completeness, which we define as the proportion of expressed isoforms among the annotated ones and the proportion of annotated isoforms among the expressed ones, respectively. We constructed nine sets of synthetic annotations, as opposed to the real annotation from GENCODE (Harrow et al. 2012) or Ensembl (Aken et al. 2016), with varying purity and completeness (Table 1). For example, annotation 4 has a 60% purity and a 40% completeness, meaning that 60% of the annotated isoforms are truly expressed and the annotated isoforms constitute 40% of the truly expressed isoforms.

**Table 1:** Synthetic annotations. Purity and completeness of the nine sets of synthetic annotations are calculated based on the truly expressed isoforms in the simulated data.

| synthetic annotation set | 1 | 2 | 3 | 4 | 5 | 6 | 7 | 8 | 9 |
| --- | --- | --- | --- | --- | --- | --- | --- | --- | --- |
| purity | 40% | 40% | 40% | 60% | 60% | 60% | 80% | 80% | 80% |
| completeness | 40% | 60% | 80% | 40% | 60% | 80% | 40% | 60% | 80% |

We first compared the four methods in terms of their gene-level isoform discovery accuracy. Regarding both the precision rate (i.e., the proportion of expressed isoforms in the discovered isoforms) and the recall rate (i.e., the proportion of discovered isoforms in the expressed isoforms), AIDE outperforms the other three methods with all the nine synthetic annotation sets (Figure 2, Supplemental Figures S1 and S2). Especially with the less accurate synthetic annotation sets 1-4, AIDE demonstrates its clear advantages thanks to its stepwise selection strategy, which prevents AIDE from being misled by the wrongly annotated isoforms. When the read coverage is 10× and the annotations have 40% purity (sets 1-3), the median precision rates of AIDE are as high as the 3rd-quantile precision rates of Cufflinks. In addition, AIDE achieves high precision rates (> 75%) much more frequently than the other three methods (Figure 2A). In terms of the recall rates, AIDE and Cufflinks exhibit better capability in correctly identifying truly expressed isoforms than StringTie and SLIDE. AIDE achieves high recall rates (> 75%) in more genes with the annotation sets 1-3 (purity = 40%), and its recall rates are close to those of Cufflinks when the annotation purity is increased to 60% and 80% (annotation sets 4-9) (Figure 2B). We also compared the accuracy of the four methods on the expressed but non-annotated isoforms (Supplemental Figure S3a-b), and AIDE demonstrates greater accuracy in identifying these novel isoforms. In addition, AIDE is robust to sequencing depths. On the contrary, the recall rates of StringTie and SLIDE deteriote as sequencing depths decrease (Supplemental Figure S2). Regarding to what extent does the accuracy of annotations affect isoform discovery, we find that the annotation purity is more important than the annotation completeness for isoform discovery (Figures 2, S1, and S2). This observation suggests that if practitioners have to choose between two annotation sets, one with high purity but low completeness and the other with low purity but high completeness, they should use the former annotation set as input into AIDE. When no annotation is given, AIDE still presents higher accuracy to predict isoform structures (Supplemental Figure S3c).

**Figure 2:**
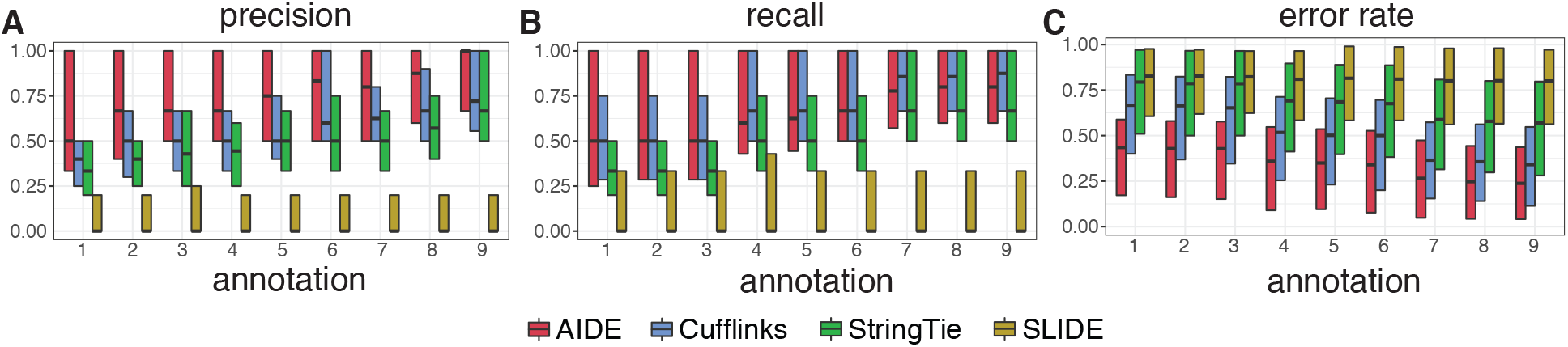
Gene-level isoform discovery and abundance estimation results of AIDE, Cufflinks, StringTie, and SLIDE on simulated RNA-seq data with 10× coverage. Each box gives the 1st quantile, median, and 3rd quantile of the gene-level accuracy given each of the nine sets of synthetic annotations. **A**: precision rates in isoform discovery; **B**: recall rates in isoform discovery; **C**: error rates (defined as one half of the sum of the absolute differences between the true and estimated isoform proportions) in abundance estimation.

As a concrete example, we considered the human gene *DPM1* and its annotated isoforms in annotation sets 1 and 9 (Table 1). For *DPM1*, the annotation set 1 has a 67% purity and a 67% completeness, and the annotation set 9 has a 60% purity and a 100% completeness. In Supplemental Figures S4 and S5, we plotted the distribution of RNA-seq reads in the reference genome, along with the truly expressed isoforms, the annotated isoforms, and the isoforms identified by AIDE, Cufflinks, and StringTie, respectively. Thanks to its capacity to selectively incorporate information from the annotated isoforms, AIDE successfully identified the shortest truly expressed isoform, which is missing in the annotation set 1 (Supplemental Figure S4). With the annotation set 9 that contains two non-expressed isoforms, AIDE correctly identified the three truly expressed isoforms (Supplemental Figure S5). In contrast, the other three methods missed some of the truly expressed isoforms with both annotation sets, and they identified too many non-expressed isoforms with the less pure annotation set 9.

We also summarized the genome-wide average accuracy of AIDE and the other three methods at three different levels: base, exon, and transcript levels (Supplemental Figure S6, calculation formulas in Supplemental Methods). All the four methods have high accuracy at the base and exon levels regardless of the annotations quality. However, even when exons are correctly identified, it remains challenging to accurately assemble exons into full-length isoforms. At the transcript level, AIDE achieves the best precision rates, recall rates comparable to Cufflinks, and the best *F*_1_ scores with all the synthetic annotation sets. In addition to the reconstruction accuracy based on the initial output of each method, we also compared the precision-recall curves of different methods by applying varying thresholds on the estimated isoform expression levels (Figure 3). Regardless of the annotation quality, AIDE achieves higher precision than the other three methods when all the methods lead to the same recall. It is worth noting that the results of AIDE were filtered by statistical significance before thresholded by expression values, while the results of the other three methods were only thresholded by isoform expression. Therefore, it is not proper to directly compare the maximum recall rates or area under the curve (AUC) scores of different methods. Nonetheless, AIDE still has the largest AUC scores. These results demonstrate the advantage of AIDE in achieving high precision and low false discovery rates in isoform discovery.

**Figure 3:**
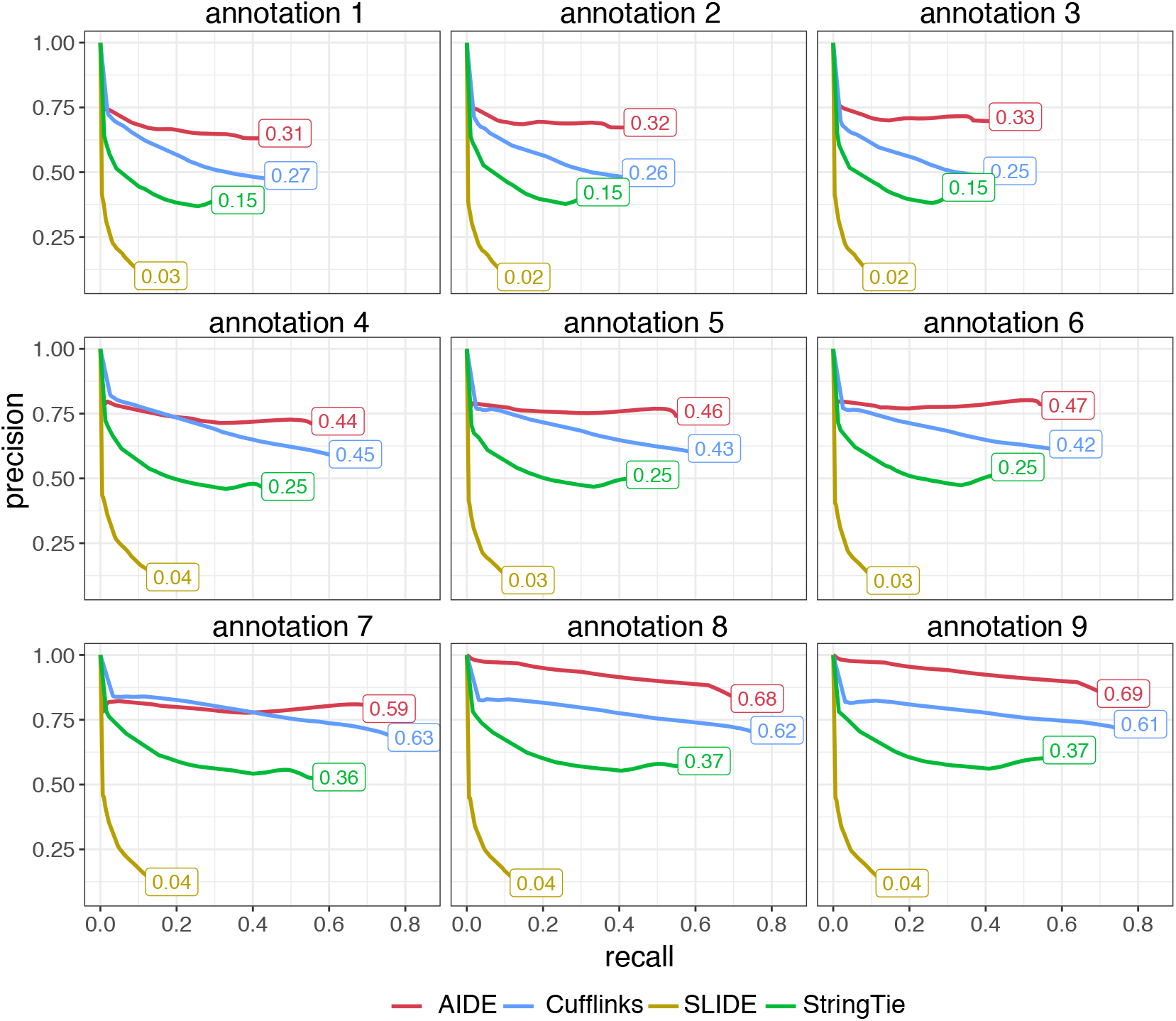
Comparison between AIDE and the other three isoform discovery methods in simulation. Given each synthetic annotation set, we applied AIDE, Cufflinks, StringTie, and SLIDE for isoform discovery, and summarized the expression levels of the predicted isoforms using the FPKM (Fragments Per Kilobase Million) unit. Then the precisionrecall curves were obtained by thresholding the FPKM values of the predicted isoforms. The AUC of each method is also marked in the plot. The shown results are based on RNA-seq data with a 10× coverage.

As the proportions of the expressed isoforms were specified in this simulation study, we also compared AIDE with the other three methods in terms of their accuracy in isoform abundance estimation. We use ***α*** = (*α*_1_, …, *α*_*J*_)′ to denote the proportions of *J* possible isoforms enumerated from a given gene’s known exons, and we use 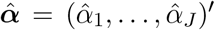 to denote the estimated proportions by a method 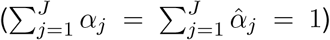. We define the estimation error rate as: 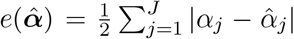. The error rate is a real value in [0, 1], with a value of 0 representing a 100% accuracy. With all the nine synthetic annotation sets, AIDE achieves the overall smallest error rates (Figure 2C). The advantages of AIDE over the other three methods are especially obvious with the first three annotations, whose purity is only 40%.

### AIDE improves isoform discovery on real data

We performed a transcriptome-wide comparison of AIDE and the other three methods based on real RNA-seq datasets. Since transcriptome-wide benchmark data are unavailable for real RNA-seq experiments, we used the isoforms in the GENCODE annotation as a surrogate basis for evaluation (Harrow et al. 2012). For every gene, we randomly selected half of the annotated isoforms and input them as partial annotations into every isoform discovery method. For RNA-seq data, we collected three human embryonic stem cell (ESC) datasets and three mouse bone marrow-derived macrophage (BMDM) datasets (Supplemental Table S1). For each gene, we applied AIDE, Cufflinks, StringTie, and SLIDE to these six datasets for isoform discovery with partial annotations, and we evaluated each method by comparing their identified isoforms with the complete set of annotated isoforms. Although the annotated isoforms are not equivalent to the truly expressed isoforms in the six samples from which RNA-seq data were generated, the identified isoforms, if accurate, are supposed to largely overlap with the annotated isoforms given the quality of human and mouse annotations. Especially if we assume the human and mouse annotations are unions of known isoforms expressed under various well-studied biological conditions including human ESCs and mouse BMDMs, it is reasonable to use those annotations to estimate the precision rates of the discovered isoforms, i.e., what proportions of the discovered isoforms are expressed. Estimation of the recall rates is more difficult, because the annotations are likely to include some isoforms that are non-expressed in human ESCs or mouse BMDMs.

We summarized the accuracy of the four isoform discovery methods at both the exon level and the transcript level (Figure 4). At the exon level, AIDE has the highest precision and recall rates on all the six datasets, achieving *F*_1_ scores greater than 90% (Figure 4A-B). The second best method at the exon level is StringTie. Since connecting exons into the full-length isoforms is much more challenging than simply identifying the individual exons, all the methods have lower accuracy at the isoform level than at the exon level. Although having recall rates slightly lower than but similar to Cufflinks, AIDE achieves the highest precision rates (~ 70% on human datasets and ~ 60% on mouse datasets) at the isoform level (Figure 4C-D). Moreover, when all the methods achieve the same recall rates after thresholding the estimated isoform expression levels (in FPKM unit), AIDE has the largest precision rate on all the six samples (Supplemental Figure S7). Although precision and recall rates are both important measures for isoform discovery, high precision results (equivalently, low false discovery results) of computational methods are often preferable for experimental validation. In the three mouse BMDM samples, AIDE identified novel isoforms for genes *MAPKAPK2, CXCL16*, and *HIVEP1*, which are known to play important roles in macrophage activation (Limbourg et al. 2015; Zhang et al. 2009; Schultze and Schmidt 2015). In our previous simulation results, we have shown that the accuracy of annotations is a critical factor determining the performance of AIDE. Even though the partial annotations used in this study only have a 50% completeness, AIDE achieves the best precision rates among the four methods, and we expect AIDE to achieve high accuracy when supplied with annotations of better quality in real applications.

**Figure 4:**
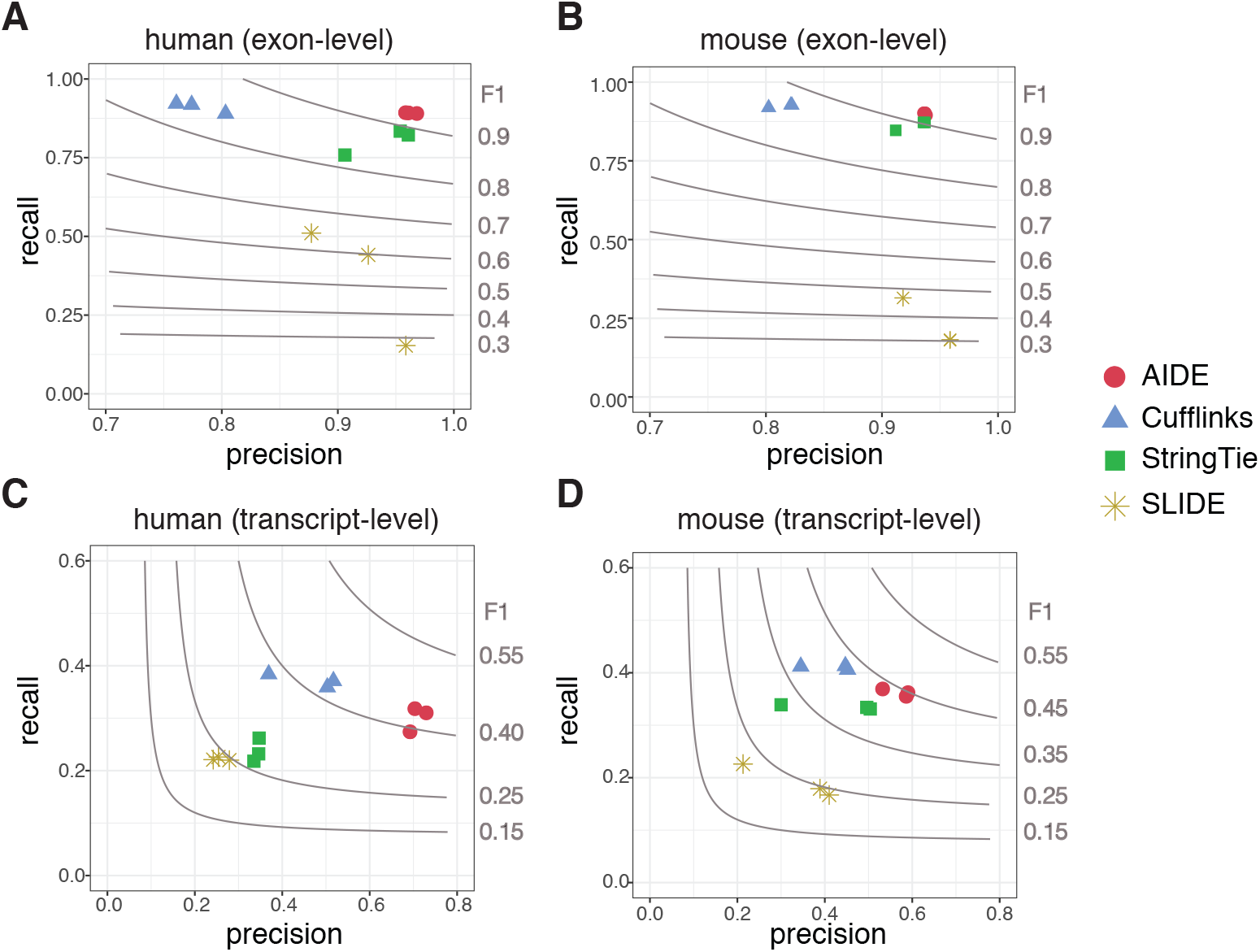
Comparison of AIDE and the other three methods on real data. **A**: exon-level accuracy in the human ESC samples; **B**: exon-level accuracy in the mouse BMDM samples; **C**: transcript-level accuracy in the human ESC samples; **D**: transcript-level accuracy in the mouse BMDM samples. The gray contours denote the *F*_1_ scores, as marked on the right of each panel.

AIDE is able to achieve more precise isoform discovery than existing methods because it utilizes a statistical model selection principle to determine whether to identify a candidate isoform as expressed. Therefore, only those isoforms that are statistically supported by the observed reads are retained. We used four example genes *ZBTB11*, *TOR1A*, *MALSU1*, and *SRSF6* to illustrate the superiority of AIDE over the other three methods (Supplemental Figures S8 and S9). The genome browser plots show that AIDE identifies the annotated isoforms with the best precision, while the other methods either miss some annotated isoforms or lead to too many false discoveries. We also used these four genes to show that AIDE is robust to the choice of the *p*-value threshold used in the likelihood ratio tests (Methods). The default choice of the threshold is 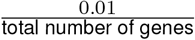, which is 4.93 × 10^−7^ for human samples and 4.54 × 10^−7^ for mouse samples. We used AIDE to identify isoforms with different thresholds and tracked how the results change while the threshold decreases from 10^−2^ to 10^−10^ (Supplemental Figures S10 and S11). As expected, AIDE tends to discover slightly more isoforms when the threshold is larger, and it becomes more conservative with a smaller threshold. However, the default threshold leads to accurate results for those four genes, and the discovered isoform set remains stable around the default threshold.

### AIDE achieves the best consistency with long-read sequencing technologies

We conducted another transcriptome-wide study to evaluate the isoform discovery methods by comparing their reconstructed isoforms (from the second-generation, short RNA-seq reads) to those identified by the third-generation long-read sequencing technologies, including Pacific Biosciences (PacBio) (Rhoads and Au 2015) and Oxford Nanopore Technologies (ONT) (Mikheyev and Tin 2014). Even though PacBio and ONT platforms have higher sequencing error rates and lower throughputs compared to second-generation sequencing technologies, they are able to generate much longer reads (1-100 kbp) to simultaneously capture multiple splicing junctions (Weirather et al. 2017). Here we used the full-length transcripts identified from the PacBio or ONT sequencing data as a surrogate gold standard to evaluate the isoform discovery methods.

We applied AIDE, Cufflinks, StringTie, and SLIDE to a second-generation RNA-seq sample of human ESCs, and compared their identified isoforms with those discovered from PacBio or ONT data generated from the same ESC sample (Weirather et al. 2017). The comparison based on ONT and PacBio are highly consistent: AIDE achieves the best precision and the highest overall accuracy (*F*_1_ score) at both the base level and the transcript level (Figure 5). We also compared the precision-recall curves of different methods by applying varying thresholds on the predicted isoform expression levels (Supplemental Figure S12). When all four methods achieve a recall rate greater than 30%, AIDE has the highest precision. The fact that all the three methods have high accuracy at the exon level but much lower accuracy at the transcript level again indicates the difficulty of assembling exons into full-length isoforms based on short RNA-seq reads. Among all the four methods, AIDE is advantageous in achieving the best precision in full-length isoform discovery.

**Figure 5:**
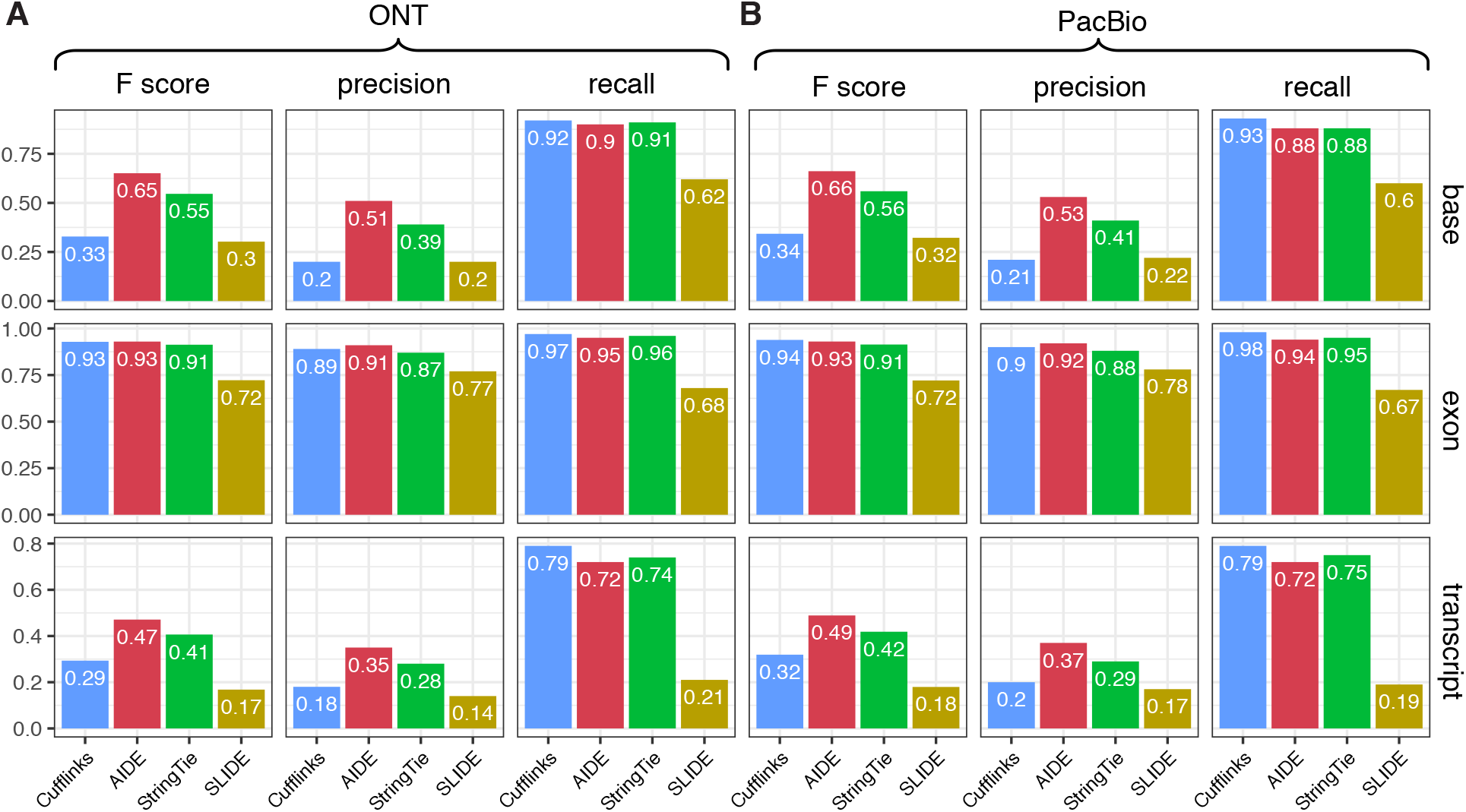
Evaluation of isoform discovery methods based on long reads. The *F*_1_ score, precision, and recall of the four discovery methods were calculated at the base, exon, and transcript levels. **A**: Evaluation based on isoforms identified by ONT; **B**: Evaluation based on isoforms identified by PacBio.

### PCR-Sanger sequencing validates the precision of AIDE

Since AIDE and Cufflinks have demonstrated higher accuracy than other methods in the assessment of genome-wide isoform discovery, we further evaluated the performance of these two methods on a small cohort of RNA-seq datasets using PCR followed by Sanger sequencing. We applied both AIDE and Cufflinks to five breast cancer RNA-seq datasets for isoform discovery.

After summarizing the genome-wide isoform discovery results, we randomly selected ten genes that have annotated transcripts uniquely predicted by only AIDE or Cufflinks with FPKM > 2 for experimental validation (Supplemental Table S2, experimental details in Supplemental Methods). We divided the genes into two categories: six genes with annotated isoforms identified only by Cufflinks but not by AIDE (category 1), and four genes with annotated isoforms identified only by AIDE but not by Cufflinks (category 2). For four out of the six genes in category 1, *MTHFD2*, *NPC2*, *RBM7*, and *CD164*, our experimental validation found that the isoforms uniquely predicted by Cufflinks were false positives (Figure 6A-D). Specifically, both AIDE and Cufflinks correctly identified the full-length isoforms *MTHFD2-201*, *NPC2-207*, *RBM7-203*, *CD164-003* for the four genes, respectively. However, the isoforms *MTHFD2-203*, *NPC2-205*, *RBM7-208*, *CD164-210* predicted only by Cufflinks were all false discoveries. For two out of the four genes in category 2, we validated the isoforms uniquely predicted by AIDE as true positives (Figure 6E-F). In detail, AIDE correctly identified isoforms *FGFR1-238* and *FGFR1-201* for gene *FGFR1*, as well as isoform *ZFAND5-208* for gene *ZFAND5*. On the other hand, Cufflinks only identified *FGFR1-201* and missed the other two isoforms. The experimental validation for both category 1 and category 2 genes presented results that are consistent with our genome-wide computational results.

**Figure 6:**
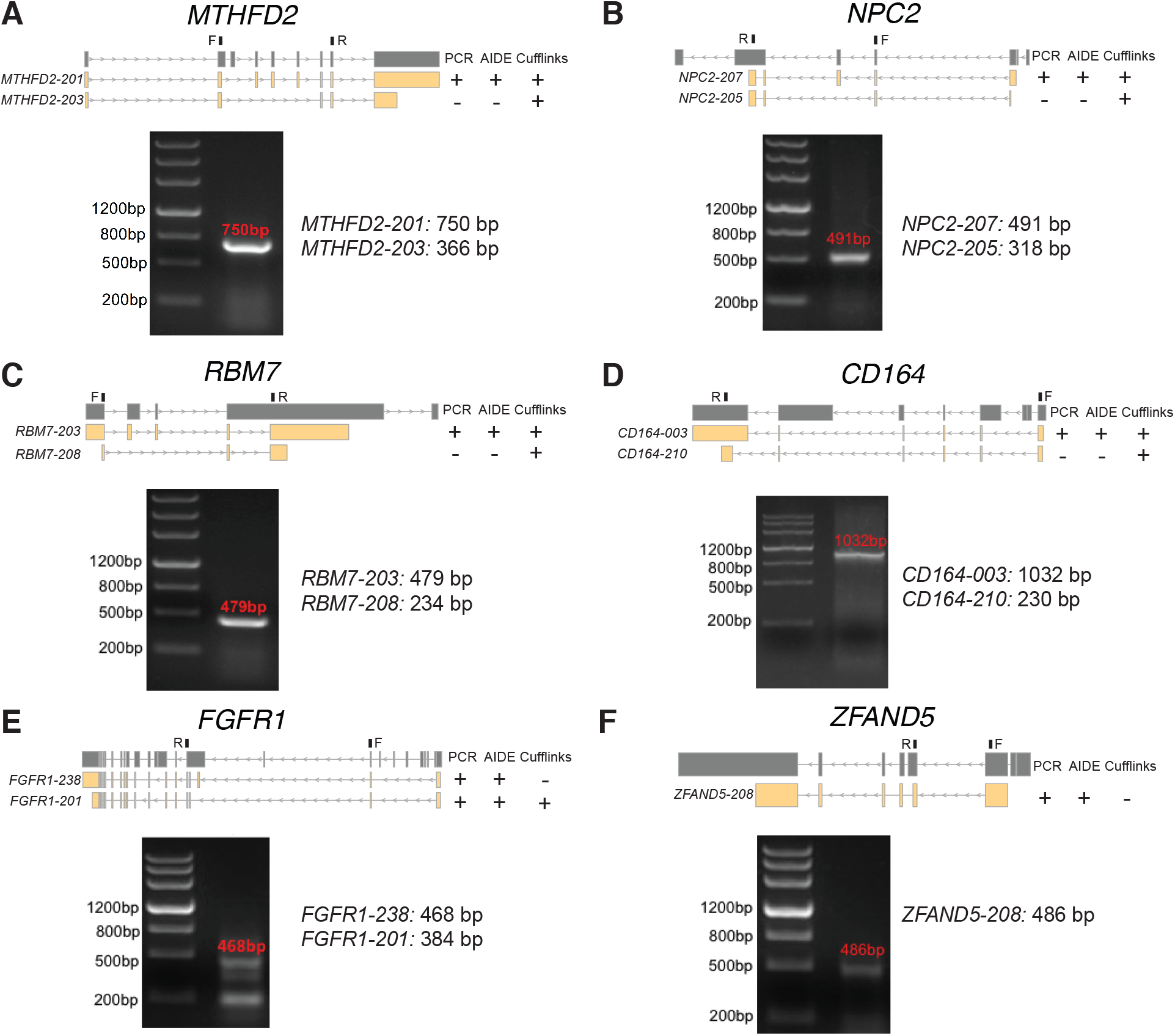
Experimental validation of isoforms predicted by AIDE and Cufflinks. Isoforms of genes *MTHFD2* (**A**), *NPC2* (**B**), *RBM7* (**C**), *CD164* (**D**), *FGFR1* (**E**), and *ZFAND5* (**F**) were validated by PCR and Sanger sequencing. The isoforms to validate (yellow) are listed under each gene (dark gray), with + / − indicating whether an isoform was / was not identified by PCR or a computational method. The forward (F) and reverse (R) primers are marked on top of each gene. For each gene, the agarose gel electrophoresis result demonstrates the molecular lengths of PCR products.

### AIDE identifies isoforms with biological relevance

We investigated the biological functions of *FGFR1-238*, an isoform predicted by AIDE but not by Cufflinks. Since *FGFR1-238* was identified in breast cancer RNA-seq samples, we evaluated its functions in breast cancer development by a loss-of-function assay. In detail, we validated the expression of *FGFR1-238* in breast cancer cell lines MCF-7, BT549, SUM149, MB231, BT474, and SK-BR-3 using PCR (Figure 7A), and we designed primers to uniquely amplify a sequence of 533 bp in its exon 18 (Supplemental Table S3). Results show that high levels of *FGFR1-238* were detected in cell lines MCF-7, BT549, MB231, and BT474 (Figure 7A). Next, we designed five small interfering RNAs (siRNAs) that specifically target the unique coding sequence of *FGFR1-238* (Supplemental Table S4). Then we studied the dependence of tumor cell growth on the expression of *FGFR1-238* by conducting a long-term (10 days) cell proliferation assay in the presence or absence (control) of siRNA knockdown. Our experimental results show that the knockdown of the *FGFR1-238* isoform inhibits the survival of MCF-7 and BT549 cells (Figure 7B). Therefore, *FGFR1* and especially its isoform *FGFR1-238* could be promising targets for breast cancer therapy, implying the ability of AIDE in identifying full-length isoforms with biological functions in pathological conditions.

**Figure 7:**
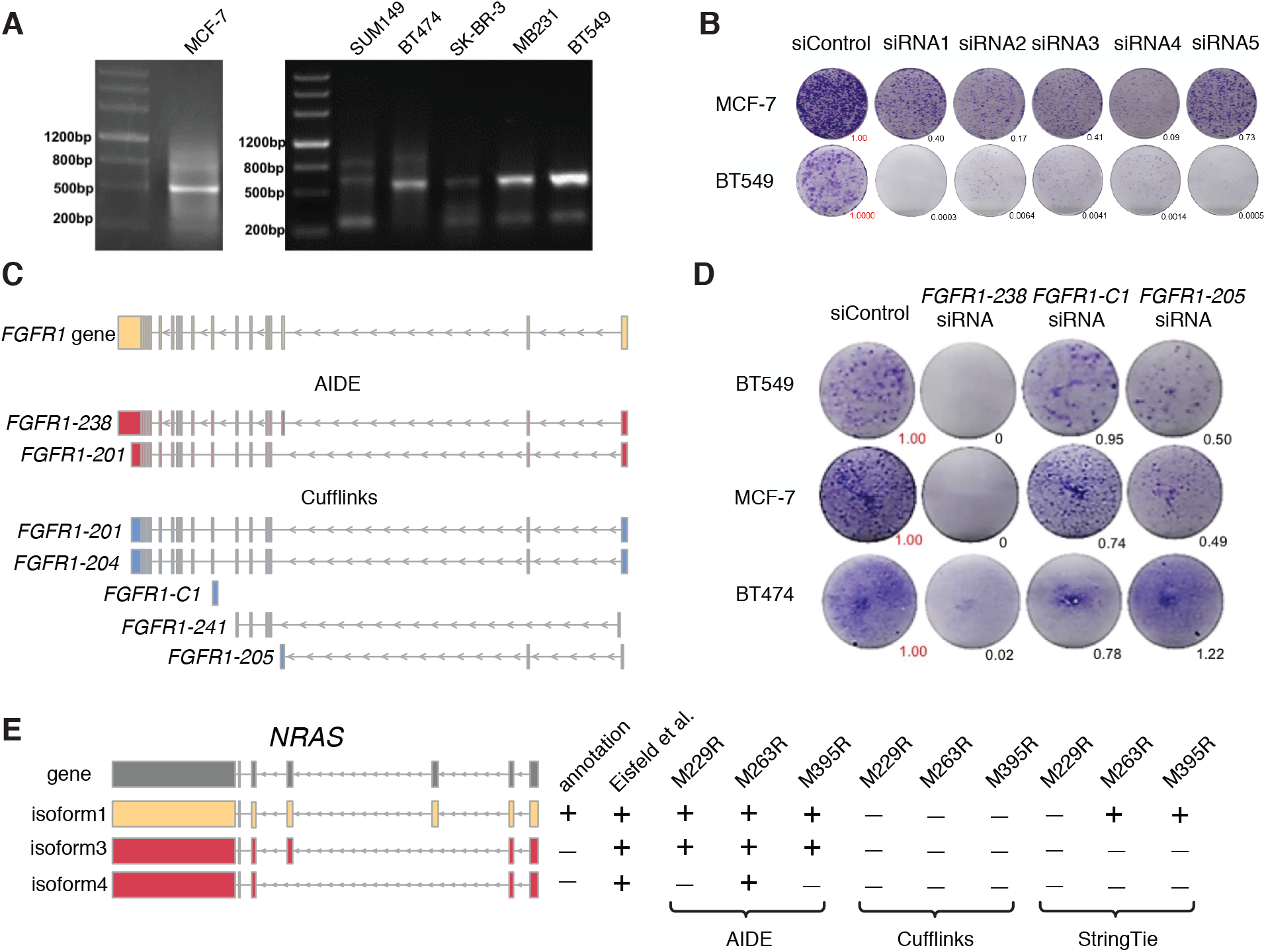
AIDE identifies isoforms with biological relevance. **A**: PCR experiments validated the expression of *FGFR1-238* in breast cancer cell lines MCF-7, SUM149, BT474, SK-BR-3, MB231, and BT549. **B**: Long-term colonegenic assay with lipo3000 controls (“siControl”) and *FGFR1-238* knockdowns. Tumor growths relative to the siControl were quantified by the ImageJ software (Schneider et al. 2012). **C**: *FGFR1* isoforms identified by AIDE and Cufflinks. **D**: Long-term colonegenic assay with siControl (negative control), si-*FGFR1-238* (positive control), si-*FGFR1-205*, and si-*FGFR1-C1*. Tumor growths relative to the siControl were quantified by the ImageJ software. **E**: *NRAS* isoforms in the GENCODE annotation, reported by Eisfeld et al. (2014), and discovered by AIDE, Cufflinks, or StringTie in three melanoma BRAF inhibitor resistant cell lines: M229R, M263R, and M395R.

To validate the specific biological function of isoform *FGFR1-238*, we also designed siRNAs targeting two non-functional isoforms *FGFR1-205* and *FGFR1-C1* (non-annotated) which were predicted by Cufflinks (Figure 7C, Supplemental Table S4). The expressions of *FGFR1-205* and *FGFR1-C1* were validated in three breast cancer cell lines, BT549, MCF-7, and BT474, by PCR experiments (Supplemental Figure S13). The lengths of the amplified *FGFR1-205* and *FGFR1-C1* are 510 bp and 528 bp, respectively. The three isoforms were knocked down in the host mammalian cells BT549, MCF-7, and BT474 by RNA interference, respectively. The results of the colonegenic assay show that only the deletion of *FGFR1-238* but not *FGFR1-205* or *FGFR1-C1* impacted long term cell survival (Figure 7D). This isoform-specific function, which was only identified by AIDE in this case, further highlights the importance of full-length isoform identification.

To further compare AIDE and other reconstruction methods in identifying isoforms with biological functions, we applied AIDE, Cufflinks, and StringTie to the RNA-seq data of three melanoma cell lines. As one of the most predominant driver oncogenes, the tumorigenic function of *NRAS* was well described (Goel et al. 2006). Recently, some novel isoforms of the *NRAS* gene were experimentally identified using quantitative PCR and shown to have potential roles in cell proliferation and malignancy transformation (Eisfeld et al. 2014). Except for the canonical isoform 1, the other two isoforms identified by Eisfeld et al. have not been included in the GENCODE (Harrow et al. 2012) or Ensembl (Zerbino et al. 2017) annotation (Figure 7E). In our previous work, we profiled the transcriptomes of a serial of melanoma cell lines (Moriceau et al. 2015; Song et al. 2017). We applied both AIDE and the other two reconstruction methods to the RNA-seq data of these cell lines. Our results showed that, among the expressed isoforms, (1) two novel *NRAS* isoforms, in addition to the annotated isoform (isoform 1), were identified by AIDE; (2) only isoform 1 was identified in two out of the three cell lines by StringTie; 3) none of them was identified by Cufflinks (Figure 7E). These results again demonstrate the potential of AIDE as a powerful bioinformatics tool for isoform discovery from short-read sequencing data.

### AIDE improves isoform abundance estimation on real data

Since the structures and abundance of expressed isoforms are unobservable in real RNA-seq data, we seek to evaluate the performance of different methods by comparing their estimated isoform expression levels in the FPKM unit with the NanoString counts, which could serve as benchmark data for isoform abundance when PCR validation is not available (Steijger et al. 2013; Geiss et al. 2008; Germain et al. 2016; Li et al. 2018). The NanoString nCounter technology is considered as one of the most reproducible and robust medium-throughput assays for quantifying gene and isoform expression levels (Kulkarni 2011; Prokopec et al. 2013; Veldman-Jones et al. 2015). We expect an accurate isoform discovery method to discover a set of isoforms close to the expressed isoforms in an RNA-seq sample. If the identified isoforms are accurate, the subsequently estimated isoform abundance is more likely to be accurate and agree better with the NanoString counts.

We therefore applied AIDE, Cufflinks, and StringTie to six samples of the human HepG2 (liver hepatocellular carcinoma) immortalized cell line with both RNA-seq and NanoString data (Steijger et al. 2013) (Supplemental Table S1). The NanoString nCounter technology is not designed for genome-wide quantification of RNA molecules, and the HepG2 NanoString datasets have measurements of 140 probes corresponding to 470 isoforms of 107 genes. Since one probe may correspond to multiple isoforms, we first found the isoforms compatible with every probe, and we then compared the sum or the maximum of the estimated abundance of these isoforms with the count of that probe. For each HepG2 sample, we calculated the Spearman’s correlation coefficient between the estimated isoform abundance (“sum” or “max”) and the NanoString probe counts to evaluate the accuracy of each method (Figure 8). AIDE has the highest correlations in five out of the six samples, suggesting that AIDE achieves more accurate isoform discovery as well as better isoform abundance estimation in this application. It is also worth noting that all three methods have achieved high correlation with the Nanostring counts for samples 3 and 4, since these two samples have the longest reads of 100 bp among all the samples. It is well acknowledged that longer reads assist isoform identification by capturing more exon-exon junctions (Li et al. 2018).

**Figure 8:**
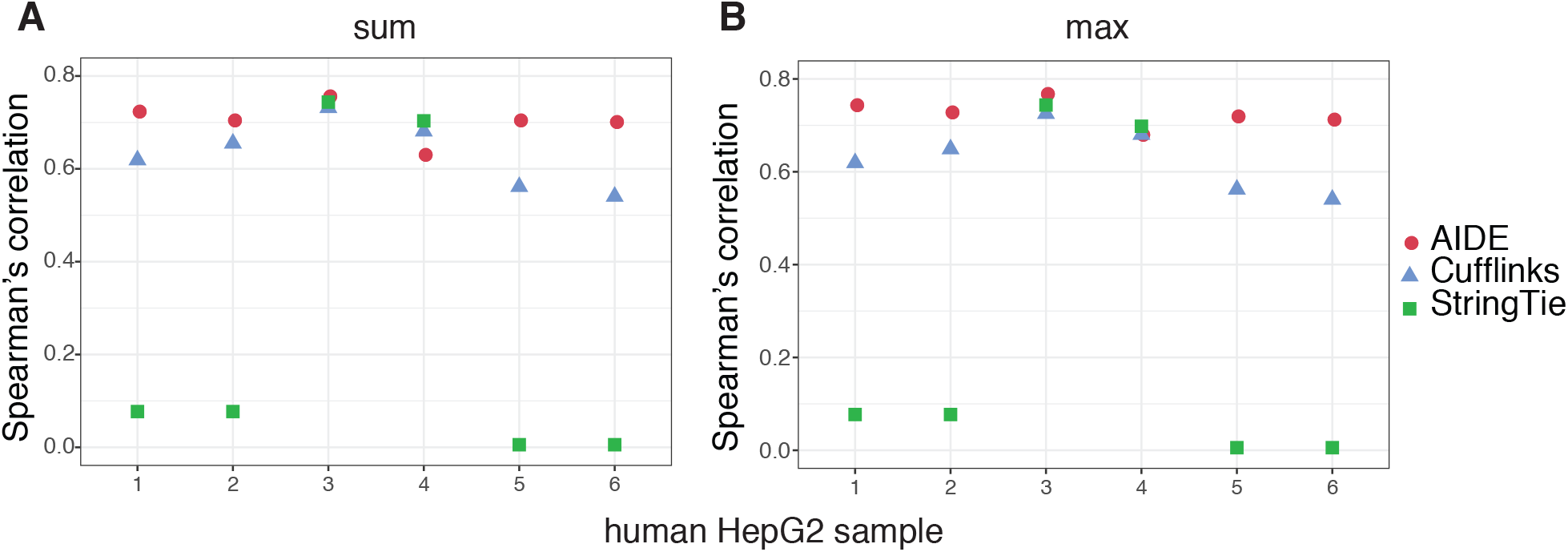
Spearman’s correlation coefficients between the estimated isoform expression and the benchmark NanoString counts. **A**: For every probe, the sum of the expression levels of its corresponding isoforms is used in the calculation. **B**: For every probe, the maximum of the expression levels of its corresponding isoforms is used in the calculation.

### AIDE improves isoform discovery accuracy via stepwise selection

We also conducted a proof-of-concept simulation study to verify the accuracy of our proposed AIDE method. We used this study to show why simply performing forward selection is insufficient and how stepwise selection leads to more robust isoform discovery results. Here we considered 2, 262 protein-coding genes from the human GENCODE annotation (version 24) (Harrow et al. 2012). We treated the annotated isoforms as the true isoforms and simulated paired-end RNA-seq reads from those isoforms with pre-determined abundance levels (detailed simulation strategy in Supplemental Methods). For every gene, we applied AIDE, which uses stepwise selection, and its counterpart AIDEf, which only uses forward selection, to discover isoforms from the simulated reads. To evaluate the robustness of AIDE to the accuracy of annotation, we considered three types of annotation sets: (1) “N” (no) annotations: no annotated isoforms were used; (2) “I” (inaccurate) annotations: the “annotated isoforms” consisted of half of the randomly selected true isoforms and the same number of false isoforms; (3) “A” (accurate) annotations: the “annotated isoforms” consisted of half of the randomly selected true isoforms.

The simulation results show that AIDE and AIDEf perform the best when the “A” annotations are supplied, and they have the worst results with the “N” annotations, as expected (Supplemental Figure S14). Given the “A” annotations, AIDE and AIDEf have similarly good performance. However, when supplied with the “I” or “N” annotations, AIDE has much better performance than AIDEf. In addition, the performance of AIDE with the “I” annotations is close to that with the “A” annotations, demonstrating the robustness of AIDE to inaccurate annotations. On the other hand, AIDEf has decreased precision rates when the “I” annotations are supplied because forward selection is incapable of removing non-expressed annotated isoforms from its identified isoform set in stage 1 (Figure 1). Given the “N” annotations, AIDE also has better performance than AIDEf. These results suggest that choosing stepwise selection over forward selection is reasonable for AIDE because perfectly accurate annotations are usually unavailable in real scenarios. Supplemental Figure S14 also suggests that both approaches exhibit improved performance with all the three types of annotations as the read coverages increase. Higher read coverages help the most when the “N” annotations are supplied, but its beneficial effects become more negligible with the “A” annotations. Moreover, we observe that the *F*_1_ scores of AIDE with the “N” annotations and the 80× coverage are approximately 30% lower than the *F*_1_ scores with the “A” annotations and the 30× coverage. This suggests that accurate annotations can assist isoform discovery and reduce the costs for deep sequencing depths to a large extent.

We also summarized the precision and recall of AIDE at the individual gene level (Supplemental Figure S15). When the “A” annotations are supplied, both AIDE and AIDEf achieve 100% precision and recall rates for over 80% of the genes. When the “N” or “I” annotations are supplied, we observe a 2.0- or 2.6- fold increase in the number of genes with 100% precision and recall rates from AIDEf to AIDE. These results again demonstrate the effectiveness of AIDE in removing non-expressed annotated isoforms and identifying novel isoforms with higher accuracy due to its use of statistical model selection principles.

## Discussion

We propose a new method AIDE to improve the precision of isoform discovery and the accuracy of isoform quantification from the second-generation RNA-seq data, by selectively borrowing full-length isoform information from annotations. AIDE iteratively identifies isoforms in a stepwise manner while placing priority on the annotated isoforms, and it performs statistical testing to automatically determine what isoforms to retain. We demonstrate the efficiency and superiority of AIDE compared to three state-of-the-art methods, Cufflinks, SLIDE, and StringTie, on multiple synthetic and real RNA-seq datasets followed by an experimental validation through PCR-Sanger sequencing, and the results suggest that AIDE leads to much more precise discovery of full-length RNA isoforms and more accurate isoform abundance estimation. In an evaluation based on the third-generation long-read RNA-seq data, AIDE also leads to the most consistent isoform discovery results than the other methods do.

In addition to reducing false discoveries, AIDE is also demonstrated to identify full-length mRNA isoforms with biological relevance in disease conditions. First, we assessed the biological relevance of the isoform *FGFR1-238*, which was only identified by AIDE, using a loss-of-function assay. We selected breast cancer samples that originally had this isoform expressed, and we experimentally proved that cell proliferation was inhibited with this isoform being knocked down. Second, we applied AIDE, Cufflinks, and StringTie to RNA-seq data of melanoma cell lines for isoform discovery. Only AIDE was able to detect two expressed but non-annotated isoforms of *NRAS*, which were reported to play a role in the drug resistance mechanism of BRAF-targeted therapy.

Due to technical limitations, the isoforms not amplified by PCR may still exist at an extremely low level. We attempted to reduce this possibility by optimizing the design and parameters used in our PCR experiments. First, we only validated the isoforms uniquely predicted by AIDE or Cufflinks if those isoforms have comparable abundance estimates (in FPKM unit). Hence, the PCR amplifications started with similar template amounts. Second, we designed the PCR primers to preferably amplify the isoforms predicted by Cufflinks, so that if Cufflinks correctly identifies an isoform, the PCR experiment would capture it with high confidence. Specifically, when PCR primers are compatible with multiple isoforms of different lengths but similar abundance levels, PCR reaction preferentially amplifies the shorter isoform(s). Therefore, we experimentally validated the genes for which the isoform predicted by AIDE is longer than the isoform predicted by Cufflinks, and we designed the primers to be compatible with both isoforms. Third, we performed extensive amplification by setting the PCR cycle number to 50. Therefore, if an isoform is not captured by the PCR, it either does not exist or has extremely low abundance level. In either case, the isoform is not supposed to be biologically functional. Given the above considerations in experimental design and the validation results (Figure 6), we could safely draw the conclusion that AIDE has unique advantages in identifying full-length mRNA isoforms with a high precision and reducing false discoveries.

Even though long reads generated by PacBio and ONT have advantages over second-generation short RNA-seq reads for assembling full-length isoforms, it remains necessary to improve computational methods for short-read based isoform discovery. First, wide application of the long-read sequencing technologies is still hindered by their lower throughput, higher error rate, and higher cost per base (Rhoads and Au 2015). Meanwhile, the second-generation short-read sequencing technology is still the mainstream assay for transcriptome profiling. Second, a huge number of second-generation RNA-seq datasets have been accumulated over the past decade. Considering that many biological or clinical samples used to generate those datasets are precious and no longer available for long-read sequencing, the existing short-read data constitute an invaluable resource for studying RNA mechanisms in these samples. Therefore, an accurate isoform discovery method will be indispensable for studying full-length isoforms from these data. Meanwhile, we also expect that with increased availability of long read data, we will be better equipped to compare and evaluate the reconstruction methods for short read data.

To the best of our knowledge, AIDE is the first isoform discovery method that identifies isoforms by selectively leveraging information from annotations using a testing-based model selection approach. The stepwise likelihood ratio testing procedure in AIDE has multiple advantages. First, AIDE only selects the isoforms that significantly contribute to the explanation of the observed reads, leading to more precise results and reduced false discoveries than those of existing methods. Second, the forward steps allow AIDE to start from and naturally give priority to the annotated isoforms, which have higher chances to be expressed in a given sample. Meanwhile, the backward steps allow AIDE to adjust its previously selected isoforms given its newly added isoform so that all the selected isoforms together better explain the observed reads. Third, the testing procedure in AIDE allows the users to adjust the conservatism and precision of the discovered isoforms according to their desired level of statistical significance. Because of these advantages, AIDE identifies fewer novel isoforms at a higher precision level than previous methods do, making it easier for biologists to experimentally validate the novel isoforms. In applications where the recall rate of isoform discovery is of great importance (i.e., the primary goal is to discover novel isoforms with a not-too-stringent criterion), users can increase the *p*-value threshold of AIDE to discover more novel isoforms.

Through the application of AIDE to multiple RNA-seq datasets, we demonstrate that selectively incorporating annotated splicing patterns, in addition to simply obtaining gene and exon boundaries from annotations, greatly helps isoform discovery. The stepwise selection in AIDE also differentiates it from the methods that directly assume the existence of all the annotated isoforms in an RNA-seq sample. The development and application of AIDE has lead us to interesting observations that could benefit both method developers and data users. First, we find that a good annotation can help reduce the need for deep sequencing depths. AIDE has been shown to achieve good accuracy on datasets with low sequencing depths when supplied with accurately annotated isoforms, and its accuracy is comparable to that based on deeply sequenced datasets. Second, we find it more important for an annotation to have high purity than to have high completeness, in order to improve the isoform discovery accuracy of AIDE and the other methods we compared with in our study. Ideally, instead of using all the annotated isoforms in isoform discovery tasks, a better choice is to use a filtered set of annotated isoforms with high confidence. This requires annotated isoforms to have confidence scores, which unfortunately are unavailable in most annotations. Therefore, how to add confidence scores to annotated isoforms becomes an important future research question, and answering this question will help the downstream computational prediction of novel isoforms.

In analysis tasks of discovering differential splicing patterns between RNA-seq samples from different biological conditions, a well-established practice is to first estimate the isoform abundance in each sample by using a method like Cufflinks, and then perform statistical testing to discover differentially expressed isoforms (Seyednasrollah et al. 2013; Trapnell et al. 2012). However, as we have demonstrated in both synthetic and real data studies, existing methods suffer from high risks of predicting false positive isoforms, i.e., estimating non-zero expression levels for unexpressed isoforms in a sample. Such false positive isoforms will severely reduce the accuracy of differential splicing analysis, leading to inaccurate comparison results between samples under two conditions, e.g., healthy and pathological samples. In contrast, AIDE’s conservative manner in leveraging the existing annotations allows it to identify truly expressed isoforms at a greater precision and subsequently estimate isoform abundance with a higher accuracy. We expect that the application of AIDE will increase the accuracy of differential splicing analysis, lower the experimental validation costs, and lead to new biological discoveries at a higher confidence level.

The probabilistic model in AIDE is very flexible and can incorporate reads of varying lengths and generated by different platforms. The non-parametric approach to learning the read generating mechanism makes AIDE a data-driven method and does not depend on specific assumptions of the RNA-seq experiment protocols. Therefore, a natural extension of AIDE is to combine the short but more accurate reads from the second-generation technologies with the longer but more error-prone reads generated by new sequencing technologies such as PacBio (Rhoads and Au 2015) and Nanopore (Byrne et al. 2017). Joint modeling of the two types of reads using the AIDE method has the potential to greatly improve the overall accuracy of isoform detection (Fu et al. 2018), since AIDE is shown to have better precision than existing methods, and longer RNA-seq reads capture more splicing junctions and can further improve the recall rate of AIDE. A second extension of AIDE is to allow the detection of novel, non-annotated exon boundaries from RNA-seq reads, as considered by the Cufflinks and StringTie Methods. Aside from the stepwise selection procedure used by AIDE, another possible way to incorporate priority on the annotated isoforms in the probabilistic model is to add regularization terms only on the unannotated isoforms. However, this approach is less interpretable compared with AIDE, since the regularization terms lack direct statistical interpretations as the *p*-value threshold does. Moreover, this approach may lose control of the false discovery rate when the annotation has a low purity. Another future extension of AIDE is to jointly consider multiple RNA-seq samples for more robust and accurate transcript reconstruction. It has been shown that it is often possible to improve the accuracy of isoform quantification by integrating the information in multiple RNA-seq samples (Li et al. 2018; Lin et al. 2012; Behr et al. 2013). Especially, using our previous method MSIQ we have demonstrated that it is necessary to account for the possible heterogeneity in the quality of different samples to improve the robustness of isoform quantification (Li et al. 2018). Therefore, by extending AIDE to combine the consistent information from multiple technical or biological samples, it is likely to achieve better reconstruction accuracy, and enable researchers to integrate publicly available and new RNA-seq samples for transcriptome studies.

## Methods

### Isoform discovery and abundance estimation using AIDE

The AIDE method is designed to identify and quantify the mRNA isoforms of each gene independently. Suppose that a gene has *m* non-overlapping exons (see Supplemental Methods for a detailed definition) and *J* candidate isoforms. If no filtering steps based on prior knowledge or external information is applied to reduce the set of candidate isoforms, *J* equals 2^*m*^ − 1, the number of all possible combinations of exons into isoforms. The observed data are the *n* RNA-seq reads mapped to the gene: ***R*** = {*r*_1_, …, *r*_*n*_}. The parameters we would like to estimate are the isoform proportions ***α*** = (*α*_1_, …, *α*_*J*_)′, where *α*_*j*_ is the proportion of isoform *j* among all the isoforms (i.e., the probability that a random RNA-seq read is from isoform *j*) and 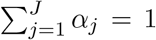. We also introduce hidden variables ***Z*** = {*Z*_1_, …, *Z*_*n*_} to denote the isoform origins of the *n* reads, with *Z*_*i*_ = *j* indicating that read *r*_*i*_ is from isoform *j*, and *P* (*Z*_*i*_ = *j*) = *α*_*j*_, for *i* = 1*, …, n*.

The joint probability of read *r*_*i*_ and its isoform origin can be written as:

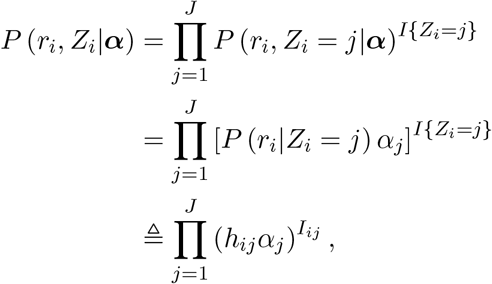

where 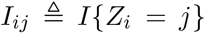 indicates whether read *r*_*i*_ is from isoform *j*, and 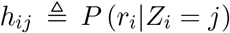 is the generating probability of read *r*_*i*_ given isoform *j*, calculated based on the read generating mechanism described in the following subsection.

### Read generating mechanism

We have defined *h*_*ij*_ as the generating probability of read *r*_*i*_ given isoform *j*. Specifically, if read *r*_*i*_ is not compatible with isoform *j* (read *r*_*i*_ contains regions not overlapping with isoform *j*, or vice versa), then *h*_*ij*_ = 0; otherwise, *h*_*ij*_ = *P* (starting position of read *r*_*i*_ | isoform *j*)*P*(fragment length of read *r*_*i*_ | isoform *j*) 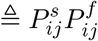.

In the literature, different models have been used to calculate the starting position distribution *P^s^* and the fragment length distribution *P^f^*. Most of these models are built upon a basic model:

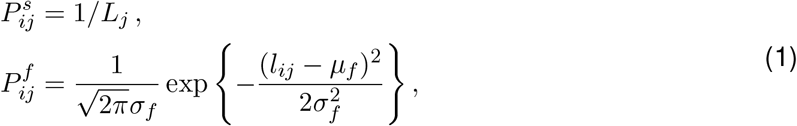

where *L*_*j*_ is the effective length of isoform *j* (the isoform length minus the read length), and *l*_*ij*_ is the length of fragment *i* given that read *r*_*i*_ comes from isoform *j*. However, this basic model does not account for factors like the GC-content bias or the positional bias. Research has shown that these biases affect read coverages differently, depending on different experimental protocols. For example, reverse-transcription with poly-dT oligomers results in an over-representation of reads in the 3’ ends of isoforms, while reverse-transcription with random hexamers results in an under-representation of reads in the 3’ ends of isoforms (Finotello et al. 2014). Similarly, different fragmentation protocols have varying effects on the distribution of reads within an isoform (Frazee et al. 2015). Existing methods such as Cufflinks (Roberts et al. 2011b), StringTie (Pertea et al. 2015), SLIDE (Li et al. 2011a), and Salmon (Patro et al. 2017) all adapted model (1) to account for varying bias sources.

Given these facts, we decide to use a non-parametric method to estimate the distribution *P^s^* of read starting positions, because non-parametric estimation is intrinsically capable of accounting for the differences in the distribution due to different protocols. We use a multivariate kernel regression to infer *P^s^* from the reads mapped to the annotated single-isoform genes. Suppose there are a total of *c*_*s*_ exons in the single-isoform genes. For *k*′ = 1, …, *c*_*s*_, we use *q*_*k′*_ to denote the proportion of reads, whose starting positions are in exon *k*′, among all the reads mapped to the gene containing exon *k*′. Given any gene with *J* isoforms, suppose there are *c*_*j*_ exons in its isoform *j*, *j* = 1, …, *J*. We estimate the conditional probability that a (random) read *r*_*i*_ starts from exon *k* (*k* = 1, 2, …, *c*_*j*_), given that the read is generated from isoform *j*, as:

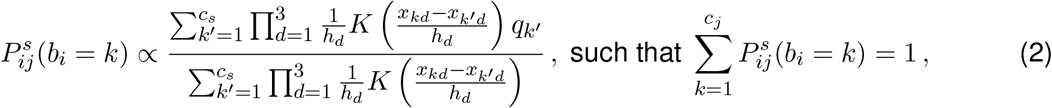

where *b*_*i*_ denotes the (random) index of the exon containing the starting position of the read *r*_*i*_. When *b*_*i*_ = *k*, we use *x*_*k*1_, *x*_*k*2_, and *x*_*k*3_ to denote the GC content, the relative position, and the length of exon *k*, respectively. The GC content of exon *k*, *x*_*k*1_, is defined as the proportion of nucleotides G and C in the sequence of exon *k*. The relative position of exon *k*, *x*_*k*2_, is calculated by first linearly mapping the genomic positions of isoform *j* to [0, 1] (i.e., the start and end positions are mapped to 0 and 1 respectively) and then locating the mapped position of the center position of the exon. For example, if isoform *j* spans from position 100 to position 1100 in a chromosome, and the center position of the exon is 200 in the same chromosome, then the relative position of this exon is 0.1. The meaning of *x*_*k*′*d*_ (*k*′ = 1, …, *c*_*s*_; *d* = 1, …, 3) are be defined in the same way.

The kernel function *K*(·) is set as the Gaussian kernel 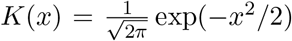. *h*_*d*_ denotes the bandwidth of dimension *d* and is selected by cross validation. The whole estimation procedure of 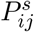 is implemented through the R package `np`.

As for the fragment length distribution *P*^*f*^, Cufflinks uses the truncated Gaussian distribution and SLIDE uses the truncated Exponential distribution. In contrast, we assume that the fragment length follows a truncated log normal distribution. This is because mRNA fragments that are too long or too short are filtered out in the library preparation step before the sequencing step. In addition, the empirical fragment length distribution is usually skewed to right instead of being symmetric (Supplemental Figure S16). Therefore, a truncated log normal distribution generally fits well the empirical distribution of fragment lengths:

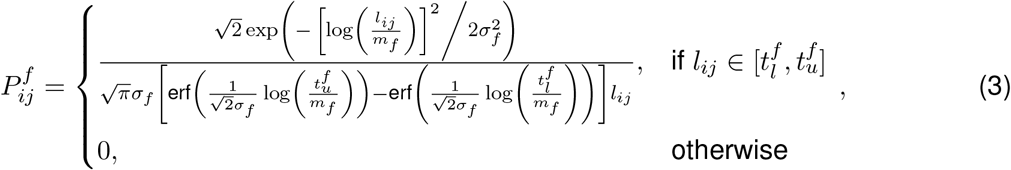

where *m*_*f*_ and *σ*_*f*_ are the median and the shape parameter of the distribution, respectively; 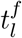 is the lower truncation threshold, and 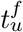 is the upper truncation threshold. The function erf(·) (the “error function” encountered in integrating the normal distribution) is defined as 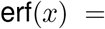 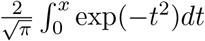.

### The probabilistic model and parameter estimation in AIDE

Given the aforementioned settings, the joint probability of all the observed and hidden data is

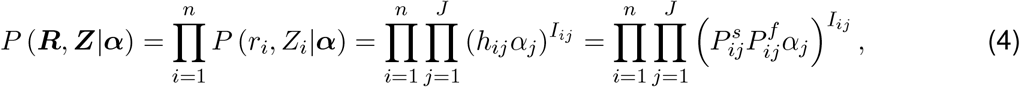

where 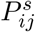 and 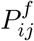 are defined in Equations (2) and (3), and the complete log-likelihood is

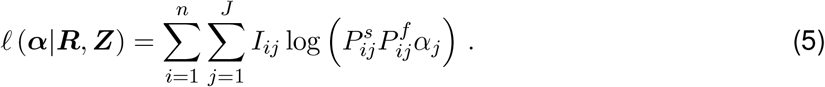

However, as ***Z*** and the resulting *I*_*ij*_’s are unobservable, the problem of isoform discovery becomes to estimate ***α*** via maximizing the log-likelihood based on the observed data:

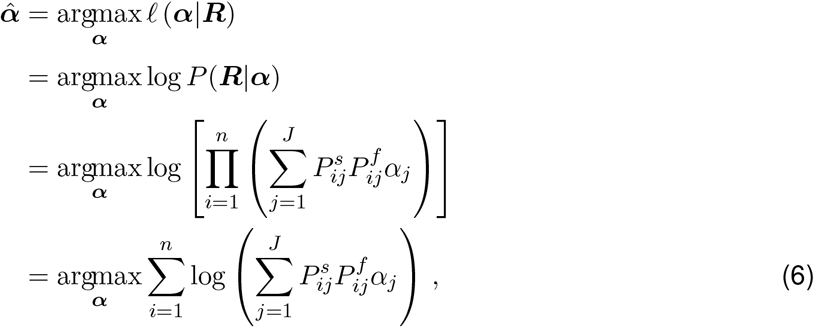

subject to *α*_*j*_ ≥ 0 and 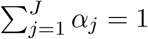. To directly solve Eq. (6) is not easy, so we use the expectation-maximization (EM) algorithm along with the complete log-likelihood Eq. (5), and it follows that we can iteratively update the estimated isoform proportions as

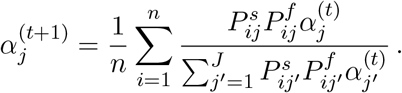

As the algorithm converges, we obtain the estimated isoform proportion 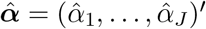.

### Stepwise selection in AIDE

If we directly consider all the *J* = 2^*m*^ − 1 candidate isoforms in formula (6) and calculate 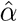, the problem is unidentifiable when *J* > *n*. Even when *J* ≤ *n*, this may lead to many falsely discovered isoforms whose 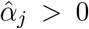, especially for complex genes, because the most complex model with all the possible candidate isoforms would best explain the observed reads. Therefore, instead of directly using the EM algorithm to maximize the log-likelihood with all the possible candidate isoforms, we perform a stepwise selection of isoforms based on the likelihood ratio test (LRT). This approach has two advantages. On the one hand, we can start from a set of candidate isoforms with high confidence based on prior knowledge, and then we can sequentially add new isoforms to account for reads that cannot be fully explained by existing candidate isoforms. For example, a common case is to start with annotated isoforms. On the other hand, the stepwise selection by LRT intrinsically introduces sparsity into the isoform discovery process. Even though the candidate isoform pool can be huge when a gene has a large number of exons, the set of expressed isoforms is usually much smaller in a specific biological sample. LRT can assist us in deciding a termination point where adding more isoforms does not further improve the likelihood.

The stepwise selection consists of steps with two opposite directions: the forward step and the backward step (Figure 1). The forward step aims at finding a new isoform to best explain the RNA-seq reads and significantly improve the likelihood given the already selected isoforms. The backward step aims at rectifying the isoform set by removing the isoform with the most trivial contribution among the selected isoforms. Since stepwise selection is in a greedy-search manner, some forward steps, especially those taken in the early iterations, may not be the globally optimal options. Therefore, backward steps are necessary to correct the search process and result in a better solution path for the purpose of isoform discovery.

We separate the search process into two stages. We use stepwise selection to update the identified isoforms at both stages, but the initial isoform sets and the candidate sets are different in the two stages. Stage 1 starts with a single annotated isoform that explains the most number of reads, and it considers all the annotated isoforms as the candidate isoforms. Stage 1 stops when the the forward step can no longer find an isoform to add to the identified isoform set, i.e., the LRT does not reject the null hypothesis given the *p*-value threshold. Stage 2 starts with the isoforms identified in stage 1, and considers all the possible isoforms, including the annotated isoforms not chosen in stage 1, as the candidate isoforms (Figure 1). The initial isoform set is denoted as 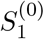 (stage 1) or 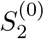 (stage 2), the candidate isoform set is denoted as *C*_1_ (stage 1) or *C*_2_ (stage 2), and the annotation set is denoted as *A*. At stage 1, the initial set and candidate set are respectively defined as

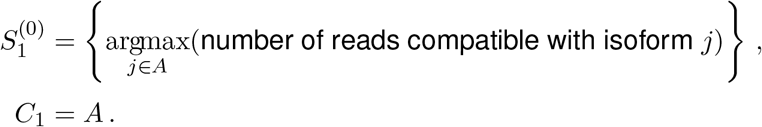

Suppose the stepwise selection completes after *t*_1_ steps in stage 1, and the estimated isoform proportions after step *t* (*t* = 1, 2, …, *t*_1_) are denoted as 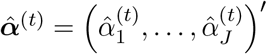. Note that ∀*j* ∉ *C*_1_, 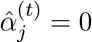 in stage 1. At stage 2, the initial set and the candidate set are respectively defined as

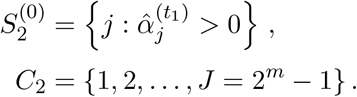

Here we introduce how to perform forward and backward selection based on a defined initial isoform set *S*^(0)^ and a candidate set *C*. We ignore the stage number subscripts for notation simplicity. At both stages, we first estimate the expression levels of the initial isoform set *S*^(0)^:

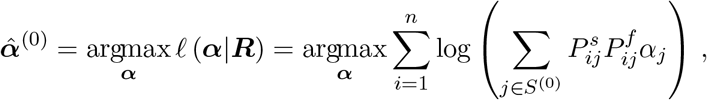

subject to *α*_j_ ≥ 0 if *j* ∈ *S*^(0)^, *α*_*j*_ = 0 if *j* ∉ *S*^(0)^, and 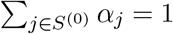, based on the EM algorithm.

### Forward step

The identified isoform set at step *t* is denoted as 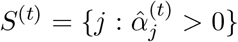. The log-likelihood at step *t* is

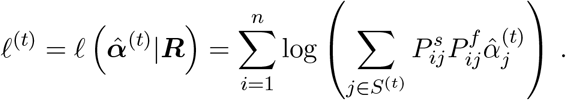

At step (*t* + 1), we consider adding one isoform *k* ∈ *C*\*S*^(*t*)^ into *S*^(*t*)^ as a forward step. Given *S*^(*t*)^ and *k*, we estimate the corresponding isoform proportions as

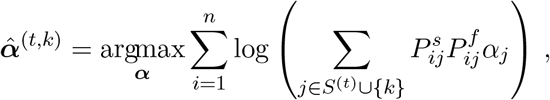

subject to *α*_*j*_ ≥ 0 if *j* ∈ *S*^(*t*)^ ∪ {*k*}, and *α*_*j*_ = 0 otherwise. Then we choose the isoform

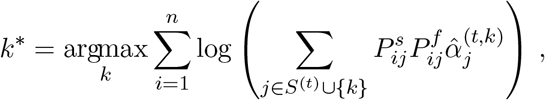

which maximizes the likelihood among all the newly added isoforms. Then the log-likelihood with the addition of this isoform *k*\* becomes

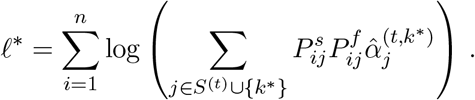

To decide whether to follow the forward step and add isoform *k*\* into the identified isoform set, we use a likelihood ratio test (LRT) to test the null hypothesis (*H*_0_: *S*^(*t*)^ is the true isoform set from which the RNA-seq reads were generated) against the alternative hypothesis (*H*_*a*_: *S*^(*t*)^ ∪ {*k*\*} is the true isoform set). Under *H*_0_ we asymptotically have −2(*ℓ*^(*t*)^ − *ℓ*\*) ~ *χ*^2^(1). If the null hypothesis is rejected at a pre-specified significance level (i.e., *p*-value threshold), then *S*^(*t*+1)^ = *S*^(*t*)^ ∪ {*k*\*}, 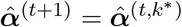, and the log-likelihood is updated as

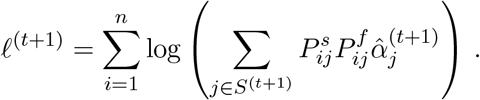

Otherwise, *S*^(*t*+1)^ = *S*^(*t*)^, 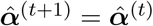 and *ℓ*^(*t*+1)^ = *ℓ*^(*t*)^.

### Backward step

Every time we add an isoform to the identified isoform set in a forward step (say the updated isoform set is *S*^(*t*+1)^), we subsequently consider possibly removing one isoform *k* ∈ *S*^(*t*+1)^ from *S*^(*t*+1)^ in a backward step. Given *S*^(*t*+1)^ and *k*, we estimate the corresponding isoform proportions as

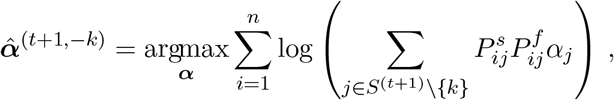

subject to *α*_*j*_ ≥ 0 if *j* ∈ *S*^(*t*+1)^\{*k*}, and *α*_*j*_ = 0 otherwise. Then we choose the isoform

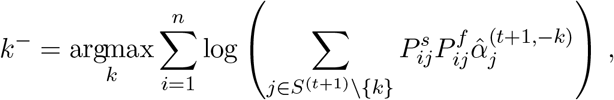

which maximizes the likelihood among all the isoforms in *S*^(*t*+1)^. Then the log-likelihood with the removal of this isoform *k*^−^ becomes

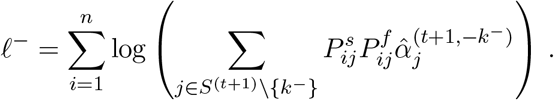

To decide whether to follow the forward step and remove isoform *k*^−^ from the identified isoform set, we use a likelihood ratio test to test the null hypothesis (*H*_0_: *S*^(*t*+1)^\{*k*^−^} is the true isoform set from which the RNA-seq reads were generated) against the alternative hypothesis (*H*_*a*_: *S*^(*t*+1)^ is the true isoform set). Under *H*_0_ we asymptotically have −2(*l*^−^ − *l*^(*t*+1)^) ~ *χ*^2^(1). If the null hypothesis is not rejected at a pre-specified significance level (i.e., *p*-value threshold), then *S*^(*t*+2)^ = *S*^(*t*+1)^\{*k*^−^}, 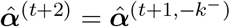, and the log-likelihood is updated as

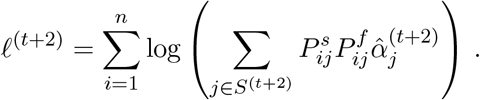

Otherwise, *S*^(*t*+2)^ = *S*^(*t*+1)^, 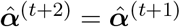, and *ℓ*^(*t*+2)^ = *ℓ*^(*t*+1)^.

In both stage 1 and stage 2, we iteratively consider the forward step and backward step and stop the algorithm at the first time when a forward step no longer adds an isoform to the identified set (Figure 1). To determine whether to reject a null hypothesis in the LRT, we set a threshold on the *p*-value. The default threshold is 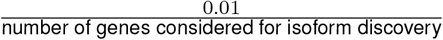. Unlike the thresholds set on the FPKM values or isoform proportions in other methods, this threshold on *p*-values allow users to tune the AIDE method based on their desired level of statistical significance. A larger threshold generally leads to more discovered isoforms and a better recall rate, while a smaller threshold leads to fewer discovered isoforms that are more precise.

### Software implementation notes

Note 1. If read *r*_*i*_ is not compatible with any isoforms in the currently selected set, then its conditional probability is theoretically 0. When implementing the AIDE method, we set this probability to *e*^−10,000^ ≈ 0. This setting requires almost all the reads to be explained by at least one selected isoform. However, if some reads present lower quality and the users want to allow some reads to be left unexplained, this probability can be set to a relatively larger value.

Note 2. Suppose a gene has *m* non-overlapping exons, then the total number of possible isoform structures is 2^*m*^ − 1. For genes with complex structures, we only consider splicing junctions supported by at least one RNA-seq read to reduce the computational complexibility. By default, we apply this filtering step to genes with more than 15 exons, but users can change the value based on their research questions.

### Software version

We carried out our analysis using the Cufflinks software v2.2.1, the StringTie software v1.3.3b, the SLIDE software, and the AIDE package v1.0.0. As an example of computational efficiency, we analyzed the human ESC Illumina RNA-seq sample (containing around 167.5 million paired-end reads) using the Ubuntu 14.04.5 system and 2 CPUs of Intel(R) Xeon(R) CPU E5-2687W v4 @ 3.00GHz. Using 12 cores, the running time of Cufflinks, StringTie, AIDE, and SLIDE was 1, 445 minutes, 196 minutes, 413 minutes, and 223 minutes, respectively. The memory usage of Cufflinks, StringTie, AIDE, and SLIDE was 17G, 8G, 25G, and 10G, respectively.

For studies included an comparison with SLIDE, the RNA-seq reads were aligned to the reference genome using STAR v2.5 (Dobin et al. 2013) with the option –alignEndsType EndToEnd and the other parameters set to default. This allows the aligned reads to have equal lengths. For studies not using SLIDE, the reads were aligned using STAR v2.5 with default parameters. The reference genomes are GRCh38 (Schneider et al. 2017) for human and GRCm38 for mouse (Church et al. 2011).

### Simulation for comparing isoform reconstruction methods

We considered 18, 960 protein-coding genes from the human GENCODE annotation (version 24). For each gene, we set the proportions of isoforms not in the GENCODE database to zero. As for the annotated isoforms in GENCODE, their isoform proportions were simulated from a symmetric Dirichlet distribution with parameters 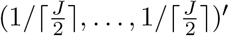, where *J* denotes the number of annotated isoforms for a given gene. When simulating the RNA-seq reads, we treated these simulated proportions as the pre-determined ground truth. Next, for each target read coverage among the eight choices (10×, 20×, …, 80×), we used the R package polyester to simulate one RNA-seq sample given the pre-determined isoform proportions. All the simulated RNA-seq samples contained paired-end reads with 100 bp length.

### Long read data processing

In the comparison with the long-read sequencing technologies, the isoforms were identifed by Weirather et al. (Weirather et al. 2017) based on the long reads generated using the ONT or PacBio technologies. In summary, the long reads were first processed using the SMRT software (https://www.pacb.com/products-and-services/analytical-software/smrt-analysis/; for PacBio) or the poretools software (https://poretools.readthedocs.io/en/latest/; for ONT). Then the reads were aligned and full-length transcripts were identified using the AlignQC software (https://www.healthcare.uiowa.edu/labs/au/AlignQC/). Please refer to Weirather et al. for details.

## Software Availability

The AIDE method has been implemented in the R package AIDE, which is available at https://github.com/Vivianstats/AIDE and also in the Supplemental code.

## Acknowledgments

The authors would like to thank Dr. Kin Fai Au and Dr. Yunhao Wang for providing their second- and third-generation human ESC RNA-seq data and processed mRNA isoforms from the third-generation data. The authors would also like to thank Dr. Alexander Hoffmann at UCLA for his insightful feedbacks. This work was supported by the following grants: UCLA Dissertation Year Fellowship (to W.V.L), PhRMA Foundation Research Starter Grant in Informatics, Johnson & Johnson WiSTEM2D Award, and Sloan Research Fellowship (to J.J.L), and National Key Research and Development Program of China (No. 2016YFC0906000 [2016YFC0906003]), National Natural Science Foundation of China (No. 81773752), Key Program of the Science and Technology Bureau of Sichuan (No. 2017SZ00005), the Recruitment Program of Global Young Experts (“the Thousand Young Talents Plan”) (to H.S.).

