## Supplemental Code for "AIDE: annotation-assisted isoform discovery with high precision": Rd2.tex

\documentclass[a4paper]{book}
\usepackage[times,inconsolata,hyper]{Rd}
\usepackage{makeidx}
\usepackage[latin1]{inputenc} % @SET ENCODING@
% \usepackage{graphicx} % @USE GRAPHICX@
\makeindex{}
\begin{document}
\chapter*{}
\begin{center}
{\textbf{\huge \R{} documentation}} \par\bigskip{{\Large of \file{AIDE}}}
\par\bigskip{\large \today}
\end{center}
