## Supplemental Code for "AIDE: annotation-assisted isoform discovery with high precision": AIDE.pdf

### Package ‘AIDE’

May 11, 2018

**Title** Annotation-based transcript reconstruction and quantification

**Version** 0.0.1

**Author** Wei Vivian Li, Jingyi Jessica Li

**Maintainer** Wei Vivian Li <>

**Description** What the package does (one paragraph).

**Depends** R (>= 3.4.2)

**License** GPL

**Encoding** UTF-8

**LazyData** true

**RoxygenNote** 6.0.1

**LinkingTo** Rcpp, RcppArmadillo

**Imports** Rcpp, RcppArmadillo, Biostrings, GenomicAlignments, GenomicFeatures, GenomeInfoDb, IRanges, S4Vectors, GenomicRanges, gtools, igraph, np, rbamtools, Rsamtools, stringr, truncdist, stats, utils, tidyr, dplyr

#### R topics documented:

aide . . . . . . . . . . . . . . . . . . . . . . . . . . . . . . . . . . . . . . . . . 1

**Index** **3**

---

`aide` *use AIDE for transcript reconstruction and quantification*

---

##### Description

use AIDE for transcript reconstruction and quantification

##### Usage

```
aide(gtf_path, bam_path, fasta_path, out_dir, readLen, strandmode = 0,
  genes = NULL, pval = NULL, ncores = 5)
```

#### Arguments

| | |
|---|---|
| gtf_path | A character specifying the full path of the GTF file. |
| bam_path | A character specifying full path of the BAM file. The BAM file should be sorted and indexed, with the BAI file in the same folder. The BAM file should be aligned using the GTF file as supplied by gtf_path. |
| fasta_path | A character specifying full path of the fasta file for genome sequences, used in GC-content bias correction. |
| out_dir | A character specifying the full path of the output directory. |
| readLen | An integer giving the length of the RNA-seq reads. |
| strandmode | An integer specifying the library type: 0 means unstranded, 1 means second-strand, and strandmode 2 means firststrand. Default is 0. |
| genes | An character vector specifying the ids of genes to be estimated. Must match the gene ids in the GTF file. Default is NULL, meaning that all genes in the GTF file will be estimated. |
| pval | An number specifying the threshold on p-values used in the likelihood ratio tests. Default is 0.01/(number of genes estimated). |
| ncores | A integer specifying the number of cores used for parallel computation. Default is 5. |

#### Value

aide saves a GTF file with reconstructed transcripts and their FPKM values to x to the directory out_dir.

#### Author(s)

Wei Vivian Li, <>

Jingyi Jessica Li, <>

### Index

aide, 1
