## Supplemental Code for "AIDE: annotation-assisted isoform discovery with high precision": aide.R

```
main = function(cgene, bam_path, bamTotal, readLen, cutoff, readm, readsd,
                sig_fward, sig_bward, starts_data, genome, bws, strandmode,
                mode = "regular", ne = 25){
  n = cgene$exonNum
  exon_start = cgene$exonStarts
  exon_end = cgene$exonEnds
  exon_len = cgene$exonLens

  reads = get_reads_from_bam(cgene, num_thre = 2, bam_path, strandmode = strandmode)
  if (is.null(reads)) return(NULL)

  nreads_ub = round(sum(exon_len)* 1.5)
  if (nrow(reads) > max(6000,nreads_ub)){
    reads = reads[sample(1:nrow(reads), nreads_ub), , drop = FALSE]
  }

  rpkm_gene = 10^9 * nrow(reads) / ((sum(exon_len)-readLen) * bamTotal)

  J_annt = cgene$txNum
  IsoM_annt = t(sapply(1:J_annt, function(j){
    tp = rep(0, n)
    tp[cgene$txs[[j]]] = 1
    return(tp)
  }))

  # use reads to filter candidates
  if(mode == "regular"){
    if(n <= 10){
      IsoM = as.matrix(expand.grid( lapply(numeric(n), function(x) c(0, 1)) ))[-1, ]
    }else if (n <= 15 ){
      IsoM = try(filter_Isomat_by_reads_nlarge(reads, nExon=n, exon_len, readLen, cutoff, nthre = n-8),
                 silent = TRUE)
      if (class(IsoM) == "matrix"){
        IsoM = rbind(IsoM, IsoM_annt)
        IsoM = unique(IsoM, MARGIN = 1)
      }
    }else if (n <= ne){
      IsoM = try(filter_Isomat_by_reads_nlarge(reads, nExon=n, exon_len, readLen, cutoff, nthre = n-6),
                 silent = TRUE)
      if (class(IsoM) == "matrix"){
        IsoM = rbind(IsoM, IsoM_annt)
        IsoM = unique(IsoM, MARGIN = 1)
      }else{ IsoM = IsoM_annt}
    }else{ IsoM = IsoM_annt }

    Ind_annt = sapply(1:J_annt, function(i){
      comp = sweep(IsoM, MARGIN = 2, IsoM_annt[i,], FUN = "-")
      tp = which(rowSums(abs(comp)) == 0)[1]
      if (length(tp ) == 0){return(NA)}
      return(tp)
    })

    if(nrow(IsoM) == 1){
      result = stepwise_LRT_wrap_single(Ind_annt)
    }else{
      if(is.null(bws)){
        Hmat = calculate_H_skipbias(reads, Iso_mat = IsoM, exon_len, readLen, readm, readsd, cutoff)
      }else{
        Hmat = calculate_H(reads, Iso_mat = IsoM, exon_len, readLen, readm, readsd,
                           cutoff, bws, genome, cgene, starts_data)
      }
      Hmat[is.nan(Hmat)] = 0
      ### LRT
      nreads = nrow(reads)
      gc()
      result = stepwise_LRT_wrap(J_annt, Ind_annt, Hmat, nExon = n,
                                 nreads, sig_fward, sig_bward)
    }

    txs = IsoM[result$Ind_check, , drop = FALSE]
    txs = lapply(1:nrow(txs), function(i){
      which(txs[i, ] == 1)
    })
    txslen = sapply(txs, function(x) sum(cgene$exonLens[x]))
    rpkm = 10^9 * nrow(reads) * result$alpha_check / ((txslen-readLen) * bamTotal)


    res = list(txs = txs, Ind_check = result$Ind_check, alpha_check = result$alpha_check,
               Ind_annt = Ind_annt, nExon = n,
               rpkm_gene = rpkm_gene, rpkm = rpkm)
  }

  if(mode == "strict"){
    if(n > ne){IsoM = IsoM_annt
    }else{
      if(n <= 10){
        IsoM = try(filter_Isomat_by_reads_strict(reads, nExon=n, exon_len, readLen, cutoff, nthre = n),
                   silent = TRUE)
      }else if(n <= 15){
        IsoM = try(filter_Isomat_by_reads_strict(reads, nExon=n, exon_len, readLen, cutoff, nthre = n-8),
                   silent = TRUE)
      }else if( n <= ne){
        IsoM = try(filter_Isomat_by_reads_strict(reads, nExon=n, exon_len, readLen, cutoff, nthre = n-5),
                   silent = TRUE)
      }
      # if(class(IsoM) == "try-error") print(IsoM)
      # print(paste("n", nrow(IsoM)))
      if (class(IsoM) == "matrix"){
        IsoM = rbind(IsoM, IsoM_annt)
      }else{ IsoM = IsoM_annt }
    }
    fe = which(colSums(IsoM) > 0)
    mt = matrix(0, nrow = length(fe), ncol = n)
    for(rr in 1:nrow(mt)){mt[rr, fe[rr]] = 1}
    IsoM = rbind(IsoM, mt)
    IsoM = unique(IsoM, MARGIN = 1)

    Ind_annt = sapply(1:J_annt, function(i){
      comp = sweep(IsoM, MARGIN = 2, IsoM_annt[i,], FUN = "-")
      tp = which(rowSums(abs(comp)) == 0)[1]
      if (length(tp ) == 0){return(NA)}
      return(tp)
    })

    if(nrow(IsoM) == 1){
      result = stepwise_LRT_wrap_single(Ind_annt)
    }else{
      if(is.null(bws)){
        Hmat = calculate_H_skipbias(reads, Iso_mat = IsoM, exon_len, readLen, readm, readsd, cutoff)
      }else{
        Hmat = calculate_H(reads, Iso_mat = IsoM, exon_len, readLen, readm, readsd,
                           cutoff, bws, genome, cgene, starts_data)
      }
      Hmat[is.nan(Hmat)] = 0
      ### LRT
      nreads = nrow(reads)
      gc()
      result = stepwise_LRT_wrap(J_annt, Ind_annt, Hmat, nExon = n,
                                 nreads, sig_fward, sig_bward)
    }

    txs = IsoM[result$Ind_check, , drop = FALSE]
    txs = lapply(1:nrow(txs), function(i){
      which(txs[i, ] == 1)
    })
    txslen = sapply(txs, function(x) sum(cgene$exonLens[x]))
    rpkm = 10^9 * nrow(reads) * result$alpha_check / ((txslen-readLen) * bamTotal)


    res = list(txs = txs, Ind_check = result$Ind_check, alpha_check = result$alpha_check,
               Ind_annt = Ind_annt, nExon = n,
               rpkm_gene = rpkm_gene, rpkm = rpkm)
  }


  return(res)
}

main_one = function(cgene, bam_path, bamTotal, readLen, strandmode){
  n = cgene$exonNum
  exon_start = cgene$exonStarts
  exon_end = cgene$exonEnds
  exon_len = cgene$exonLens

  reads = get_reads_from_bam(cgene, num_thre = 2, bam_path, strandmode = strandmode)
  if (is.null(reads)) return(NULL)

  rpkm_gene = 10^9 * 2 * nrow(reads) / ((sum(exon_len)-readLen) * bamTotal)

  J_annt = 1

  txs = cgene$txs[1]
  txslen = sapply(txs, function(x) sum(cgene$exonLens[x]))
  rpkm = rpkm_gene


  res = list(txs = txs, Ind_check = 1, alpha_check = 1,
             Ind_annt = 1, nExon = 1,
             rpkm_gene = rpkm_gene, rpkm = rpkm)
  return(res)
}


#' use AIDE for transcript reconstruction and quantification
#'
#' @param gtf_path A character specifying the full path of the GTF file.
#' @param bam_path A character specifying full path of the BAM file.
#' The BAM file should be sorted and indexed, with the BAI file in the same folder.
#' The BAM file should be aligned using the GTF file as supplied by \code{gtf_path}.
#' @param fasta_path A character specifying full path of the fasta file for genome sequences,
#' used in GC-content bias correction.
#' @param out_dir A character specifying the full path of the output directory.
#' @param readLen An integer giving the length of the RNA-seq reads.
#' @param strandmode An integer specifying the library type: 0 means unstranded,
#' 1 means secondstrand, and strandmode 2 means firststrand. Default is 0.
#' @param genes An character vector specifying the ids of genes to be estimated.
#' Must match the gene ids in the GTF file. Default is \code{NULL},
#' meaning that all genes in the GTF file will be estimated.
#' @param pval An number specifying the threshold on p-values used in the likelihood ratio tests.
#' Default is 0.01/(number of genes estimated).
#' @param ncores An integer specifying the number of cores used for parallel computation.
#' Default is 5.
#' @param mode An character specifying running mode. Default is "regular".
#' @param ne An integer indicating complex genes.
#' @return \code{aide} saves a GTF file with reconstructed transcripts and their FPKM values to
#' x to the directory \code{out_dir}.
#' @export
#' @import parallel
#' @import gtools
#' @import stringr
#' @import truncdist
#' @import GenomicAlignments
#' @import GenomicFeatures
#' @importFrom IRanges IRanges
#' @import S4Vectors
#' @importFrom GenomicRanges GRanges disjoin
#' @import Biostrings
#' @import rbamtools
#' @importFrom Rsamtools ScanBamParam
#' @importFrom GenomeInfoDb dropSeqlevels
#' @importFrom igraph all_simple_paths graph_from_adjacency_matrix
#' @import np
#' @importFrom stats fitted quantile setNames
#' @importFrom utils stack write.table
#' @useDynLib AIDE
#' @importFrom Rcpp sourceCpp
#' @author Wei Vivian Li, \email{}
#' @author Jingyi Jessica Li, \email{}
aide = function(gtf_path, bam_path, fasta_path, out_dir, readLen, strandmode = 0, genes = NULL, pval = NULL, ncores = 5,
                mode = "regular", ne = 25){
  options(np.messsages = FALSE)
  set.seed(1)

  tp_dir = paste0(out_dir, "temporary/")
  dir.create(tp_dir, recursive = TRUE)
  dir.create(paste0(tp_dir, "data/"), recursive = TRUE)

  print("extracting gene models from GTF ...")
  gene_models = NULL
  save_txdb_path = paste0(tp_dir, "gene_models.RData")
  gtf_to_gene_models(gtf_path, save_txdb_path, ncores)

  load(paste0(tp_dir, "gene_models.RData"))
  exonNum = sapply(gene_models, function(x) x$exonNum)
  geneNames = names(gene_models[exonNum >= 2])
  rowID = which(sapply(gene_models, function(x) x$exonNum) == 1)

  if(is.null(genes)){
    #run_genes = 1:length(geneNames)
    run_genes = 1:length(gene_models)
  }else{
    run_genes = match(genes, names(gene_models))
    if(sum(!is.na(run_genes))==0) stop("No genes match the GTF file!")
    if(sum(is.na(run_genes))>0){
      print(paste(sum(is.na(run_genes)), "genes are not present in the GTF file!"))}
    run_genes = run_genes[!is.na(run_genes)]
  }

  reader = bamReader(bam_path, idx=TRUE)
  count = bamCountAll(reader,verbose=TRUE)
  bamClose(reader)
  bamTotal = sum(count[, "nAligns"])

  print("estimating fragment length ...")
  paras = get_fragment_length_dist(gene_models, rowID, num_thre = 100,
                                   bam_path, strandmode = strandmode, quant = 0.99, ncores = ncores)
  cutoff = paras$cutoff
  if(is.na(cutoff[1])) {cutoff = c(readLen, 400)}
  readm = paras$mean
  readsd = paras$sd
  print(readm)
  print(readsd)
  print(cutoff)

  print("estimating bias ...")
  gene_models_1tx = gene_models[sapply(gene_models, function(x) x$txNum) == 1]
  exonNum = sapply(gene_models_1tx, function(x) x$exonNum)
  gene_models_1tx = gene_models_1tx[exonNum > 1]

  exonlen = lapply(gene_models_1tx, function(x) x$exonLens)
  print(paste("number of genes with 1 tx:", length(gene_models_1tx)))

  genome = readDNAStringSet(fasta_path)
  chrstrs = strsplit(names(genome), split = " ")
  chrstrs = sapply(chrstrs, function(x) x[1])
  names(genome) = chrstrs

  starts_data = get_onetx_starts_data(gene_models_1tx, num_thre = 20, strandmode = strandmode, flag = 2,
                                      bam_path, genome, ncores = ncores*2)
  if(nrow(starts_data) < 1500 *2){
    bws = NULL
    print("not enough reads for estimation ...")
    print("skipping bias estimation ...")
  }else{
    bws = get_bandwidth(starts_data, ncores = ncores)
    print(paste("bandwidth:", bws))
  }
  # bws = c(0.05952130, 0.03026894, 0.01828689)
  print(paste("bandwidth:", bws))

  print("reconstructing transcripts ...")
  if(is.null(pval)){pval = 0.01/length(geneNames)}

  LRT_res = mclapply(run_genes, function(id){
    set.seed(id)
    gc()

    cgene = gene_models[[id]]
    print(paste("gene:", id))
    if(cgene$exonNum == 1){
      result = try(main_one(cgene, bam_path, bamTotal, readLen, strandmode), silent = TRUE)
    }else{
      result = try(main(cgene, bam_path, bamTotal, readLen, cutoff,
                        readm, readsd,
                        sig_fward = pval, sig_bward = pval,
                        starts_data, genome, bws, strandmode,
                        mode = mode, ne = ne), silent = TRUE)
    }

    save(result, file = paste0(tp_dir, "data/", id, ".RData"))
    #telap = round((proc.time()-t1)["elapsed"], 2)

    return(0)
  }, mc.cores = ncores)

  print("writing results ...")
  gtf_name = paste0(out_dir, "transcripts.gtf")
  rdata_to_gtf(tp_dir, gene_models, gtf_name, ncores)
  return(0)
}
```
