## Supplemental Code for "AIDE: annotation-assisted isoform discovery with high precision": calculate_h_bias.R

```
# impute fragment lengths
# (i, j) gives the length of fragment i if it comes from tx j
# 0 if not compatible
impute_fragment_len = function(reads, Iso_mat, exon_len, nExon, readLen){
  bin_mat = reads[, c("b1", "e1", "b2", "e2"), drop = FALSE]
  coords = reads[, c("b_bpOnExon", "e_bpOnExon"), drop = FALSE]
  compat_mat = matrix( nrow=nrow(bin_mat), ncol=nrow(Iso_mat))
  M = nrow(Iso_mat)
  
  # tick = proc.time()
  # Type I: on the same exon 
  rowIds.beSameExon = which(bin_mat[ ,"b1"] == bin_mat[ ,"e2"])
  if (length(rowIds.beSameExon) > 0){
    exon.vec = bin_mat[rowIds.beSameExon, "b1"]
    ifExonInTranscript = t(sapply(exon.vec, function(x){
      Iso_mat[,x]==1
    }))
    len = coords[rowIds.beSameExon, 2] - coords[rowIds.beSameExon, 1] + 1
    compat_mat[rowIds.beSameExon,] = sweep(ifExonInTranscript, 1, len, FUN = "*")
  }
  # print(proc.time() - tick)
  
  # tick = proc.time()
  ### check later !!!
  lenOfExonsInBtw.list = list(); 
  if (nExon > 2){
    for(b.exonId in 1:(nExon-2)){
      for(e.exonId in (b.exonId+2):nExon){
        # indicating whether there are exons in between
        #print(paste(b.exonId, e.exonId))
        rowIds = sapply(1:M, function(m){ 
          sum( Iso_mat[m, c(b.exonId,e.exonId)]==0)==0})
        len =  as.vector(Iso_mat[,(b.exonId+1):(e.exonId-1), drop=FALSE] %*% 
                           matrix(exon_len[(b.exonId+1):(e.exonId-1)], ncol=1))
        len[!rowIds] = 0
        lenOfExonsInBtw.list[[paste(b.exonId,e.exonId,sep="_")]] = len
      }
    }
  }
  #print(proc.time() - tick)
  #tick = proc.time()
  
  # type II: 4-dim on 2 exons 
  rowIds.be2Exons = which(apply(bin_mat, 1, function(x) length(unique(x)) >= 2)) 
  if (length(rowIds.be2Exons) > 0){
    exonPair.typeII.mat = bin_mat[rowIds.be2Exons, , drop = FALSE] 

    ifExonPairInTranscript.typeII = 
      ifExonPairInTranscript(exonPair = exonPair.typeII.mat, Iso_mat = Iso_mat)
    
    # length of fragments on two ends
    lenv = coords[rowIds.be2Exons, 2] + exon_len[exonPair.typeII.mat[,"b1"]] - 
      coords[rowIds.be2Exons, 1] + 1
    # length matrix, read by isoform
    len = matrix(rep(lenv, nrow(Iso_mat)), ncol = nrow(Iso_mat), byrow = FALSE)
    ifskip = which(exonPair.typeII.mat[,"e2"] - exonPair.typeII.mat[,"b1"] > 1)
    if (length(ifskip) > 0){
      pair_skip = exonPair.typeII.mat[ifskip, c("b1", "e2"), drop = FALSE]
      skiped_len = lenOfExonsInBtw.list[paste(pair_skip[,"b1"], pair_skip[,"e2"], sep="_")]
      skiped_len = matrix(unlist(skiped_len, use.names = FALSE), 
                          nrow = length(ifskip), byrow = TRUE)
      len[ifskip, ] = len[ifskip, ] + skiped_len
       # Reduce(rbind, lenOfExonsInBtw.list[paste(pair_skip[,"b1"], pair_skip[,"e2"], sep="_")])
    }
    
    compat_mat[rowIds.be2Exons, ] = ifExonPairInTranscript.typeII * len
  }
  
  # check if fragment length is shorter than read length (invalid mapping)
  # left-end
  check_read_len_1st = which((bin_mat[,"b1"]+1 < bin_mat[,"e1"])) 
  if (length(check_read_len_1st) > 0){
    len1a = lenOfExonsInBtw.list[paste(bin_mat[check_read_len_1st,"b1"], 
                                       bin_mat[check_read_len_1st,"e1"], sep="_")]
    len1a = matrix(unlist(len1a, use.names = FALSE), 
                   nrow = length(check_read_len_1st), byrow = TRUE)
    # len1a = Reduce(rbind, lenOfExonsInBtw.list[paste(bin_mat[check_read_len_1st,"b1"], 
    #                                                  bin_mat[check_read_len_1st,"e1"], sep="_")])
    # if (length(check_read_len_1st) == 1){
    #   len1a = matrix(len1a, nrow = length(check_read_len_1st), byrow = TRUE)
    # }
    len1b = exon_len[bin_mat[check_read_len_1st,"b1"]] - coords[check_read_len_1st, 1] + 1
    len1 = sweep(len1a, 1, len1b, FUN = "+")
    compat_mat[check_read_len_1st, ][len1 >= readLen] = 0
  }
  # right-end
  check_read_len_2nd = which((bin_mat[,"b2"]+1 < bin_mat[,"e2"]))
  if (length(check_read_len_2nd) > 0){
    len2a = lenOfExonsInBtw.list[paste(bin_mat[check_read_len_2nd,"b2"], 
                                       bin_mat[check_read_len_2nd,"e2"], sep="_")]
    len2a = matrix(unlist(len2a, use.names = FALSE), 
                   nrow = length(check_read_len_2nd), byrow = TRUE)
    # len2a = Reduce(rbind, lenOfExonsInBtw.list[paste(bin_mat[check_read_len_2nd,"b2"], 
    #                                                  bin_mat[check_read_len_2nd,"e2"], sep="_")])
    # if (length(check_read_len_2nd) == 1){
    #   len2a = matrix(len2a, nrow = length(check_read_len_2nd), byrow = TRUE)
    # }
    len2b = coords[check_read_len_2nd, 2]
    len2 = sweep(len2a, 1, len2b, FUN = "+")
    compat_mat[check_read_len_2nd, ][len2 >= readLen] = 0
  }
  return(compat_mat)
}

# calculate the read generating mechanism matrix H_(nreads * J)
calculate_H = function(reads, Iso_mat, exon_len, readLen, readm, readsd, cutoff,
                       bws, genome, cgene, starts_data){
  nExon = ncol(Iso_mat)
  # calculate imputed fragment length
  impute_len = impute_fragment_len(reads, Iso_mat, exon_len, nExon, readLen)
  meanlog = (4*log(readm) - log(readsd^2 + readm^2))/2
  sdlog = sqrt( log(readsd^2/(readm^2) + 1))
  # calculate probability of fragment length
  Probf = dtrunc(impute_len, spec = "lnorm",
                 meanlog = meanlog, sdlog = sdlog,
                 a = cutoff[1], b = cutoff[2])
  
  Probf = matrix(Probf, ncol = ncol(impute_len), byrow = FALSE)
  Probf[impute_len == 0] = 0
  
  # calculate probability of starting position
  startprob = estimate_startprob(cgene, Iso_mat, exon_len, genome, bws, starts_data)
  
  if (cgene$str == "+"){
    starts = reads[, "b1", drop = FALSE]
  }else{starts = reads[, "e2", drop = FALSE] }
  
  Probs = t(startprob[,starts, drop = FALSE])
  
  Hmat = Probf * Probs
  return(Hmat)
}


calculate_H_skipbias = function(reads, Iso_mat, exon_len, readLen, readm, readsd, cutoff){
  nExon = ncol(Iso_mat)
  impute_len = impute_fragment_len(reads, Iso_mat, exon_len, nExon, readLen)
  meanlog = (4*log(readm) - log(readsd^2 + readm^2))/2
  sdlog = sqrt( log(readsd^2/(readm^2) + 1))
  Probf = dtrunc(impute_len, spec = "lnorm",
                 meanlog = meanlog, sdlog = sdlog,
                 a = cutoff[1], b = cutoff[2])
  
  Probf = matrix(Probf, ncol = ncol(impute_len), byrow = FALSE)
  Probf[impute_len == 0] = 0
  
  txs_len = as.numeric(Iso_mat %*% matrix(exon_len, ncol=1))
  eff_txs_len = txs_len - cutoff[1]
  Hmat = sweep(Probf, MARGIN = 2, 1/eff_txs_len, FUN = "*")
  return(Hmat)
}


# 1 if compatible
# 0 if not compatible
# for speeding up only, criteria not stringent
get_compat_mat = function(reads, Iso_mat){
  bin_mat = reads[, c("b1", "e1", "b2", "e2")]
  compat_mat = matrix( nrow=nrow(bin_mat), ncol=nrow(Iso_mat))
  M = nrow(Iso_mat)
  
  rowIds.beSameExon = which(bin_mat[ ,"b1"] == bin_mat[ ,"e2"])
  if (length(rowIds.beSameExon) > 0){
    exon.vec = bin_mat[rowIds.beSameExon, "b1"]
    ifExonInTranscript = t(sapply(exon.vec, function(x){
      Iso_mat[,x]==1
    }))
    compat_mat[rowIds.beSameExon,] = ifExonInTranscript
  }
  
  rowIds.be2Exons = which(apply(bin_mat, 1, function(x) length(unique(x)) >= 2)) 
  
  if (length(rowIds.be2Exons) > 0){
    exonPair.typeII.mat = bin_mat[rowIds.be2Exons, , drop = FALSE] 
    ifExonPairInTranscript.typeII = 
      ifExonPairInTranscript(exonPair = exonPair.typeII.mat, Iso_mat = Iso_mat)
    compat_mat[rowIds.be2Exons, ] = ifExonPairInTranscript.typeII
  }
  return(compat_mat)
}
```
