## Supplemental Code for "AIDE: annotation-assisted isoform discovery with high precision": estimate_starts_distribution_simplify.R

```
# instead of using both .fa and .fasta
# use .fa of chromosome sequences only


get_onetx_starts_data = function(gene_models, num_thre = 10, strandmode = 0, flag = 1,
                                 bam_path, genome, ncores){

  if (flag == 1){
    sel = sample(1:length(gene_models), round(length(gene_models)/2))
    gene_models = gene_models[sel]
  }
  reads_data = mclapply(1:length(gene_models), function(ii){

    if (ii %% 500 == 0) print(ii)
    cgene = gene_models[[ii]]
    genelen = sum(cgene$exonLens)

    readExoncoords = get_reads(cgene, num_thre, bam_path, strandmode = strandmode)
    if (is.null(readExoncoords)) return(NULL)
    if (cgene$str == "+"){
      starts = readExoncoords$starts
    }else{
      starts = readExoncoords$ends
    }
    exon_rpos = c(0,cumsum(cgene$exonLens))
    # fraction of reads on each exon
    readsfrac = findInterval(starts,exon_rpos+1)
    readsfrac = sapply(1:cgene$exonNum, function(x) sum(readsfrac == x))/length(starts)

    # use exon centers to calculate relative position of exons
    rpos = 0.5 * (cumsum(cgene$exonLens) + exon_rpos[1:cgene$exonNum]) / genelen

    # GC content on each exon
    dna = genome[[cgene$chr]]
    gc_exon = sapply(1:cgene$exonNum, function(i){
      start = cgene$exonStarts[i]
      end = cgene$exonEnds[i]
      dnaseq = dna[start:end]
      return(letterFrequency(dnaseq, "GC", as.prob = TRUE))
    })

    return(data.frame(readsfrac = readsfrac, rpos = rpos, gc = gc_exon,
                      exonlen = cgene$exonLens/genelen, id = ii))
  }, mc.cores = ncores)
  reads_data = Reduce(rbind, reads_data)


  return(reads_data)
}


get_bandwidth = function(starts_data, ncores){
  # if(nrow(starts_data) <= 1000)
  # nsplit = round(nrow(starts_data)/nsize)
  nsplit = 4
  geneids = unique(starts_data$id)
  ngene = length(geneids)
  test_geneid = sample(1:ngene, size = round(ngene/nsplit))
  testid = which(starts_data$id %in% geneids[test_geneid])
  test = starts_data[testid, ]
  train = starts_data[-testid, ]
  split_data = split(train, sample(1:(nsplit-1), size = nrow(train), replace=TRUE))

  bwobj_split = mclapply(1:(nsplit-1), function(i){
    print(i)
    ctrain = split_data[[i]]
    bwobj = npregbw(formula = readsfrac ~ rpos + gc + exonlen,
                    regtype = "lc",  bwmethod = "cv.ls", ckertype = "epanechnikov",
                    data = ctrain)
    return(bwobj)
  }, mc.cores = ncores)


  #bw_split = rowMeans(sapply(bwobj_split, function(x) x$bw))

  R_split = sapply(1:(nsplit-1), function(i) {
    #model = npreg(bws = bwobj_split[[i]], newdata = test, y.eval = TRUE)
    model = npreg(bws = bwobj_split[[i]],
                  txdat = train[,2:4], tydat = train[,1],
                  exdat = test[,2:4], eydat = test[,1])
    return(model$R2)
  })

  bws = bwobj_split[[which.max(R_split)]]$bw
  return(bws)
}


# get exons' relative position on each isoform
# use exons' centers
# (i, j) gives the prob of a position on exon j if it comes from tx i
# 0 if not compatible
estimate_startprob = function(cgene, Iso_mat, exon_len, genome, bws, starts_data){
  # cumulative length in isoforms
  # isoform by exon
  cum_len = sapply(1:length(exon_len), function(j){
    Iso_mat[,1:j, drop = FALSE] %*% matrix(exon_len[1:j], ncol=1)
  })
  if (nrow(Iso_mat) == 1){
    cum_len = matrix(cum_len, nrow = 1)
  }
  txs_len = cum_len[, length(exon_len)]
  cum_len = sweep(cum_len, MARGIN = 2, exon_len/2, FUN = "-")
  cum_len[Iso_mat == 0] = 0
  dat = sweep(cum_len, MARGIN = 1, txs_len, FUN = "/")
  colnames(dat) = 1:cgene$exonNum

  dat = as.data.frame(dat)
  dat$txid = 1:nrow(Iso_mat)
  exonid = NULL
  rpos = NULL
  dat = dat %>% tidyr::gather(exonid, rpos, 1:cgene$exonNum)
  dat = dat %>% dplyr::filter(rpos > 0)
  dat$exonid = as.numeric(dat$exonid)

  dna = genome[[cgene$chr]]
  gc = sapply(1:length(exon_len), function(i){
    start = cgene$exonStarts[i]
    end = cgene$exonEnds[i]
    dnaseq = dna[start:end]
    return(letterFrequency(dnaseq, "GC", as.prob = TRUE))
  })
  dat$gc = gc[dat$exonid]

  rexonlen = sweep(Iso_mat, MARGIN = 2, exon_len, FUN = "*")
  rexonlen = sweep(rexonlen, MARGIN = 1, txs_len, FUN = "/")
  dat$exonlen = rexonlen[dat$txid + nrow(rexonlen)*(dat$exonid -1)]

  # kernel regression
  # t1 = proc.time()
  model_np = npreg(bws = bws,
                txdat = starts_data[,c("rpos", "gc", "exonlen")],
                tydat = starts_data[,"readsfrac"],
                exdat = dat[,c("rpos", "gc", "exonlen")])
  # proc.time()-t1
  probs = matrix(0, ncol = ncol(cum_len), nrow = nrow(cum_len))
  probs[dat$txid + nrow(cum_len)*(dat$exonid -1)] = fitted(model_np)
  probs = sweep( probs, MARGIN = 1, rowSums( probs), FUN = "/")
  probs = sweep( probs, MARGIN = 2, exon_len, FUN = "/")
  return( probs)
}
```
