## Supplemental Code for "AIDE: annotation-assisted isoform discovery with high precision": estimation.R

```
# step one: no prior info; no regularization
est_alp_one = function(H_mat, J, init, nreads, iter = 500){
  alp_crr = init
  ii = 0
  while(ii < iter){
    print(ii)
    denom = H_mat %*% matrix(alp_crr, ncol = 1)
    alp_new = sapply(1:J, function(k){
#      sum(H_mat[,k]*alp_crr[k] / denom, na.rm = TRUE)/ nreads
      sum(H_mat[,k]*alp_crr[k] / denom, na.rm = TRUE)/ sum(denom != 0)
#       tp = H_mat[,k]*alp_crr[k] / denom
#       tp[denom == 0] = 1/J
#       return(sum(tp/nreads))
    })
    if (sum((alp_new - alp_crr)^2) < 1e-8){
      break
    }else{
      alp_crr = alp_new
    }
#    print(ii)
    ii = ii+1
  }
#  return(round(alp_crr, digits = digits))
  return(alp_crr)
}


# calculate complete likelihood function
get_Loglik = function(H_mat, alp_hat){
  # if any read cannot be explained
  # set the corresponding component to -1e10
  tp = H_mat %*% matrix(alp_hat, ncol = 1)
  ltp = log(tp)
  ltp[ltp == -Inf] = -1e100
  return(sum(ltp))
}

# get_Loglik = function(H_mat, alp_hat){
#   # if any read cannot be explained, set likelihood to -1e10
#   tp = H_mat %*% matrix(alp_hat, ncol = 1)
#   if (any(tp == 0)) {return(-1e10)
#     }else{ return (sum(log(tp))) }
# }
```
