## Supplemental Code for "AIDE: annotation-assisted isoform discovery with high precision": filter_by_reads.R

```
filter_Isomat_by_reads = function(reads, nExon, exon_len, readLen, cutoff){
  bin_mat = reads[, c("b1", "e1", "b2", "e2")]
  links = matrix(nrow = nExon, ncol = nExon)

  one_exon_ind = which(bin_mat[ ,"b1"] == bin_mat[ ,"e2"])
  if (length(one_exon_ind) > 0){
    one_exon = unique(bin_mat[one_exon_ind, "b1"])
    diag(links)[one_exon] = 1
  }

  multi_exon_ind = setdiff(1:nrow(bin_mat), one_exon_ind)
  if (length(multi_exon_ind) > 0){
    cbin = unique(bin_mat[multi_exon_ind, , drop = FALSE], MARGIN = 1)
    pair = rbind(cbin[, c("b1", "e1")], cbin[, c("b2", "e2")])
    pair = unique(pair, MARGIN = 1)
    pair = pair[pair[,2] - pair[,1] > 0, , drop = FALSE]
    if (nrow(pair) > 0){
      for (k in 1:nrow(pair)){
        #print(k)
        if (pair[k, 1]+1 == pair[k, 2]){
          links[pair[k, 1], pair[k, 2]] = 1
        }
        if(pair[k, 1]+1  < pair[k, 2]){
          # lens of exons in between cannot be longer than read length
          middles = (pair[k, 1]+1): (pair[k, 2]-1)
          middles = middles[exon_len[middles] < readLen+1]
          for(cand in middles){
            links[pair[k, 1], cand] = 1
            links[cand, pair[k,2]] = 1
          }
          # cand = sort(union(pair[k, ], middles))
          # for (ii in 1:(length(cand)-1)){
          #   for (jj in (ii+1):length(cand)){
          #     #print(paste(ii, ",", jj))
          #     links[cand[ii], cand[jj]] = 1
          #   }
          # }
        }
      }
    }

    pair = cbin[, c("e1", "b2"), drop = FALSE]
    pair = unique(pair, MARGIN = 1)
    pair = pair[pair[,2] - pair[,1] > 0, , drop = FALSE]
    if (nrow(pair) > 0){
      for (k in 1:nrow(pair)){
        if (pair[k, 1]+1 == pair[k, 2]){
          links[pair[k, 1], pair[k, 2]] = 1
        }
        if(pair[k, 1]+1  < pair[k, 2]){
          # lens of exons in between cannot be longer than max fragmemt length
          middles = (pair[k, 1]+1): (pair[k, 2]-1)
          middles = middles[exon_len[middles] < cutoff[2]+1]
          cand = sort(union(pair[k, ], middles))
          for (ii in 1:(length(cand)-1)){
            for (jj in (ii+1):length(cand)){
              #print(paste(ii, ",", jj))
              links[cand[ii], cand[jj]] = 1
            }
          }
        }
      }
    }

  }

  # softer threshold if exon length is shorter than read length
  short_ind = which(exon_len <= readLen )
  if(length(short_ind)>0){
    for (x in short_ind){
      if (x==1){ links[x, x+1]=1
      }else if(x==length(exon_len)){
        links[x-1, x]=1
      }else{
        links[x-1,x] = 1
        links[x,x+1] = 1
        }
    }
  }
  links[is.na(links)] = 0

  exonG = graph_from_adjacency_matrix(links, mode = "directed")
  paths = lapply(1:nExon, function(x) {
    all_simple_paths(exonG, from = x)
  })
  paths = unlist(paths, recursive = FALSE)
  paths = lapply(paths, function(x) as.numeric(x))

  index = stack(setNames(paths, seq_along(paths)))
  IsoM = matrix(0, ncol = nExon, nrow = length(paths))
  IsoM[(index[,1]-1)*nrow(IsoM) + as.numeric(index[,2])] = 1

  return(IsoM)
}


# when nExon is large, don't consider exons between e2 and b1
filter_Isomat_by_reads_nlarge = function(reads, nExon, exon_len, readLen, cutoff, nthre){

  bin_mat = reads[, c("b1", "e1", "b2", "e2")]
  links = matrix(nrow = nExon, ncol = nExon)

  one_exon_ind = which(bin_mat[ ,"b1"] == bin_mat[ ,"e2"])
  if (length(one_exon_ind) > 0){
    one_exon = unique(bin_mat[one_exon_ind, "b1"])
    diag(links)[one_exon] = 1
  }

  multi_exon_ind = setdiff(1:nrow(bin_mat), one_exon_ind)
  if (length(multi_exon_ind) > 0){
    cbin = unique(bin_mat[multi_exon_ind, , drop = FALSE], MARGIN = 1)
    pair = rbind(cbin[, c("b1", "e1")], cbin[, c("b2", "e2")])
    pair = unique(pair, MARGIN = 1)
    pair = pair[pair[,2] - pair[,1] > 0, , drop = FALSE]
    if (nrow(pair) > 0){
      for (k in 1:nrow(pair)){
        #print(k)
        if (pair[k, 1]+1 == pair[k, 2]){
          links[pair[k, 1], pair[k, 2]] = 1
        }
        # if(pair[k, 1]+1  < pair[k, 2]){
        #   for (ii in pair[k, 1]:(pair[k, 2]-1)){
        #       links[ii, ii+1] = 1
        #   }
        # }
        if(pair[k, 1]+1  < pair[k, 2]){
          # lens of exons in between cannot be longer than read length
          middles = (pair[k, 1]+1): (pair[k, 2]-1)
          middles = middles[exon_len[middles] < readLen+1]
          for(cand in middles){
            links[pair[k, 1], cand] = 1
            links[cand, pair[k,2]] = 1
          }
        }
      }
    }

    pair = cbin[, c("e1", "b2"), drop = FALSE]
    pair = unique(pair, MARGIN = 1)
    pair = pair[pair[,2] - pair[,1] > 0, , drop = FALSE]
    if (nrow(pair) > 0){
      for (k in 1:nrow(pair)){
        links[pair[k, 1], pair[k, 2]] = 1
      }
    }
  }

  links[is.na(links)] = 0

  exonG = graph_from_adjacency_matrix(links, mode = "directed")

  est = colSums(links)
  est = est[est != 0]
  est = sum(cumprod(est))
  if(est > 7000){
    paths = lapply(1:nExon, function(x) {
      if (nExon - x +1 < nthre) return(NULL)
      tp = all_simple_paths(exonG, from = x, to = (x+nthre-1):nExon)
      tx_nexon = sapply(tp, length)
      tp = tp[ tx_nexon >= nthre]
      return(tp)
    })
  }else{
    paths = lapply(1:nExon, function(x) {
      tp = all_simple_paths(exonG, from = x)
      return(tp)
    })
  }

  paths = unlist(paths, recursive = FALSE)
  paths = lapply(paths, function(x) as.numeric(x))

  index = stack(setNames(paths, seq_along(paths)))
  IsoM = matrix(0, ncol = nExon, nrow = length(paths))
  IsoM[(index[,1]-1)*nrow(IsoM) + as.numeric(index[,2])] = 1

  return(IsoM)
}


# when nExon is large, don't consider exons between e2 and b1
filter_Isomat_by_reads_strict = function(reads, nExon, exon_len, readLen, cutoff, nthre){

  bin_mat = reads[, c("b1", "e1", "b2", "e2")]
  links = matrix(nrow = nExon, ncol = nExon)

  one_exon_ind = which(bin_mat[ ,"b1"] == bin_mat[ ,"e2"])
  if (length(one_exon_ind) > 0){
    one_exon = unique(bin_mat[one_exon_ind, "b1"])
    diag(links)[one_exon] = 1
  }

  multi_exon_ind = setdiff(1:nrow(bin_mat), one_exon_ind)
  if (length(multi_exon_ind) > 0){
    cbin = unique(bin_mat[multi_exon_ind, , drop = FALSE], MARGIN = 1)
    pair = rbind(cbin[, c("b1", "e1")], cbin[, c("b2", "e2")])
    pair = unique(pair, MARGIN = 1)
    pair = pair[pair[,2] - pair[,1] > 0, , drop = FALSE]
    if (nrow(pair) > 0){
      for (k in 1:nrow(pair)){
        links[pair[k, 1], pair[k, 2]] = 1
      }
    }

    pair = cbin[, c("e1", "b2"), drop = FALSE]
    pair = unique(pair, MARGIN = 1)
    pair = pair[pair[,2] - pair[,1] > 0, , drop = FALSE]
    if (nrow(pair) > 0){
      for (k in 1:nrow(pair)){
        links[pair[k, 1], pair[k, 2]] = 1
      }
    }
  }

  links[is.na(links)] = 0


  exonG = graph_from_adjacency_matrix(links, mode = "directed")

  est = colSums(links)
  est = est[est != 0]
  est = sum(cumprod(est))
  if(est > 5000){
    paths = lapply(1:nExon, function(x) {
      if (nExon - x +1 <= nthre){
        tp = all_simple_paths(exonG, from = x)
      }else{tp = all_simple_paths(exonG, from = x, to = (x+nthre-1):nExon)}
      tx_nexon = sapply(tp, length)
      tp = tp[tx_nexon >= nthre]
      return(tp)
    })
  }else{
    paths = lapply(1:nExon, function(x) {
      tp = all_simple_paths(exonG, from = x)
      return(tp)
    })
  }

  paths = unlist(paths, recursive = FALSE)
  paths = lapply(paths, function(x) as.numeric(x))

  index = stack(setNames(paths, seq_along(paths)))
  IsoM = matrix(0, ncol = nExon, nrow = length(paths))
  IsoM[(index[,1]-1)*nrow(IsoM) + as.numeric(index[,2])] = 1

  return(IsoM)
}
```
