## Supplemental Code for "AIDE: annotation-assisted isoform discovery with high precision": get_fragment_length_dist.R

```
# only consider lens below 99% percentile
get_fragment_length_dist = function(gene_models, rowID, num_thre = 100,
                             bam_path, strandmode = 0, quant, ncores){
  print("start estimating mean and sd of fragment length ...")
  flag = 1
  sel = sample(1:length(rowID), round(length(rowID)/2))
  rowID1 = rowID[sel]
  rowID2 = rowID[-sel]
  fglensList = mclapply(rowID1, function(rowid){
    # print(rowid)
    cgene = gene_models[[rowid]]
    readTxcoords = get_reads(cgene, num_thre = num_thre, bam_path, strandmode = strandmode)
    if (is.null(readTxcoords)){ return(NULL)
    }else{
      return(readTxcoords$ends - readTxcoords$starts + 1)
    }
  }, mc.cores = ncores)
  gc()
  fglensList = fglensList[!sapply(fglensList, is.null)]
  fglens = unlist(fglensList)
  if(length(fglens < 10000)){
    flag = 2
    fglensList = mclapply(rowID2, function(rowid){
      # print(rowid)
      cgene = gene_models[[rowid]]
      readTxcoords = get_reads(cgene, num_thre = num_thre, bam_path, strandmode = strandmode)
      if (is.null(readTxcoords)){ return(NULL)
      }else{
        return(readTxcoords$ends - readTxcoords$starts + 1)
      }
    }, mc.cores = ncores)
    fglensList = fglensList[!sapply(fglensList, is.null)]
    fglens = c(fglens, unlist(fglensList))
  }

  print(paste("number of genes used:", length(fglensList)))
  fglens = unlist(fglensList)
  if(length(fglens) < 2000){
    print("do not have enough reads for estimation ...")
    print("using default values ...")
    return(list(mean = 250, sd = 80, cutoff = NA))
  }

  fglens = fglens[fglens < quantile(fglens, quant)]
  print(paste("number of fragments:", length(fglens)))
  print(paste("longest fragments:", max(fglens)))
  print(paste("shortest fragments:", min(fglens)))

  mean = mean(fglens)
  sd = sd(fglens)
  cutoff = range(fglens, na.rm = TRUE)
  return(list(mean = mean, sd = sd, cutoff = cutoff, flag = flag))
}
```
