## Supplemental Code for "AIDE: annotation-assisted isoform discovery with high precision": get_reads.R

```
# given a gene model
# return a data.frame readsTxcoords
# return NULL if number of reads does not reach num_thre
# filter out reads mapped to introns or with skip on the same exon

# strandmode = 0: unstranded
# strandmode = 1: secondstrand
# strandmode = 2, firststrand

#library(GenomicAlignments)

get_reads = function(cgene, num_thre, bam_path, strandmode = 0){
  genestart = cgene$exonStarts[1]
  geneend = cgene$exonEnds[cgene$exonNum]

  # get reads' coordinates from RNA-seq data
  region = GRanges(cgene$chr, IRanges(genestart, geneend), strand = cgene$str)
  param <- ScanBamParam(which = region)
  readpairs = readGAlignmentPairs(bam_path, param = param, strandMode = strandmode)
  if (strandmode !=0 ){
    readpairs = readpairs[strand(readpairs) == cgene$str]}

  reads4dim = data.frame(start(GenomicAlignments::first(readpairs)), end(GenomicAlignments::first(readpairs)),
                         start(GenomicAlignments::second(readpairs)), end(GenomicAlignments::second(readpairs)))

  # filter out reads mapped to introns
  indNoIntron = sapply(1:4, function(k){
    indexStarts <- findInterval(reads4dim[,k], cgene$exonStarts)
    indexEnds <- findInterval(reads4dim[,k], cgene$exonEnds + 1)

    ind = (indexStarts - 1 == indexEnds)
  })
  if (class(indNoIntron) == "matrix"){
    indNoIntron = rowSums(indNoIntron) == 4
  }else {return(NULL)}

  if (sum(indNoIntron) <= num_thre ) {return(NULL)}
  reads4dim = reads4dim[indNoIntron, , drop = FALSE]


  # filter out reads with skip on the same exon
  # reads4bin = sapply(1:4, function(k){
  #   findInterval(reads4dim[, k], cgene$exonStarts)
  # })

  exon_Len = cgene$exonLens
  readsTxcoords = sapply(1:4, function(k){
    indexs = findInterval(reads4dim[, k], cgene$exonStarts)
    position = sapply(indexs, function(x) sum(exon_Len[1:x])) +
      reads4dim[, k] - (cgene$exonEnds)[indexs]
    return(position)
  })

  maxReadLen = max(c(qwidth(first(readpairs)), qwidth(second(readpairs))))
  minReadLen = min(c(qwidth(first(readpairs)), qwidth(second(readpairs))))
  indNoskip = (readsTxcoords[,2] - readsTxcoords[,1] + 1 <= maxReadLen) &
    (readsTxcoords[,4] - readsTxcoords[,3] + 1 <= maxReadLen) &
    (readsTxcoords[,2] - readsTxcoords[,1] + 1 >= minReadLen) &
    (readsTxcoords[,4] - readsTxcoords[,3] + 1 >= minReadLen)

  if (sum(indNoskip) <= num_thre ) {return(NULL)}
  readsTxcoords = readsTxcoords[indNoskip, , drop = FALSE]
  #readcoords = data.frame(starts = apply(reads4dim, 1, min),
   #                       ends = apply(reads4dim, 1, max))


  # get indexes of exons overlapping with reads' starts and ends
  #indexStartsLt <- findInterval(readcoords$starts, cgene$exonStarts)
  # indexStartsRt <- findInterval(readcoords$starts,
  #                               cgene$exonEnds + 1)
  #indexEndsLt <- findInterval(readcoords$ends, cgene$exonStarts)
  # indexEndsRt <- findInterval(readcoords$ends,
  #                             cgene$exonEnds + 1)


  #starts = sapply(indexStartsLt, function(x) sum(exon_Len[1:x])) +
  #  readcoords$starts - (cgene$exonEnds)[indexStartsLt]
  #ends = sapply(indexEndsLt, function(x) sum(exon_Len[1:x])) +
  #  readcoords$ends - (cgene$exonEnds)[indexEndsLt]

  readTxcoords = data.frame(starts = apply(readsTxcoords, 1, min),
                            ends = apply(readsTxcoords, 1, max))

  return(readTxcoords)
}
```
