## Supplemental Code for "AIDE: annotation-assisted isoform discovery with high precision": get_reads_from_bam.R

```
# given a gene model
# return a matrix reads
# return NULL if number of reads does not reach num_thre
# filter out reads mapped to introns

# strandmode = 0: unstranded
# strandmode = 1: secondstrand
# strandmode = 2, firststrand

# library(GenomicAlignments)

get_reads_from_bam = function(cgene, num_thre, bam_path, strandmode = 0){
  genestart = cgene$exonStarts[1]
  geneend = cgene$exonEnds[cgene$exonNum]

  # get reads' coordinates from RNA-seq data
  region = GRanges(cgene$chr, IRanges(genestart, geneend), strand = cgene$str)
  param <- ScanBamParam(which = region)
  readpairs = readGAlignmentPairs(bam_path, param = param, strandMode = strandmode)
  if (strandmode !=0 ){
    readpairs = readpairs[strand(readpairs) == cgene$str]}

  reads4dim = data.frame(start(GenomicAlignments::first(readpairs)), end(GenomicAlignments::first(readpairs)),
                         start(GenomicAlignments::second(readpairs)), end(GenomicAlignments::second(readpairs)))
  switchInd = reads4dim[,3] <= reads4dim[,1]
  tp = reads4dim[switchInd, 3:4]
  reads4dim[switchInd, 3:4] = reads4dim[switchInd, 1:2]
  reads4dim[switchInd, 1:2] = tp

  # filter out reads mapped to introns
  indNoIntron = sapply(1:4, function(k){
    indexStarts <- findInterval(reads4dim[,k], cgene$exonStarts)
    indexEnds <- findInterval(reads4dim[,k], cgene$exonEnds + 1)
    ind = (indexStarts - 1 == indexEnds)
  })
  if (class(indNoIntron) == "matrix"){
    indNoIntron = rowSums(indNoIntron) == 4
  }else {return(NULL)}

  if (sum(indNoIntron) <= num_thre ) {return(NULL)}
  reads4dim = reads4dim[indNoIntron, , drop = FALSE]

  reads4bin = sapply(1:4, function(k){
    findInterval(reads4dim[, k], cgene$exonStarts)
  })


  b_bpOnExon = reads4dim[,1] - cgene$exonStarts[reads4bin[,1]] + 1
  e_bpOnExon = reads4dim[,4] - cgene$exonStarts[reads4bin[,4]] + 1

  reads = matrix(0, nrow = nrow(reads4dim), ncol = 6)
  for(j in 1:4){reads[,j] = reads4bin[,j]}
  reads[,5] = b_bpOnExon
  reads[,6] = e_bpOnExon
  # reads = cbind(reads4bin,  b_bpOnExon,  e_bpOnExon)
  colnames(reads) = c("b1", "e1", "b2", "e2", "b_bpOnExon", "e_bpOnExon" )

  return(reads)
}
```
