## Supplemental Code for "AIDE: annotation-assisted isoform discovery with high precision": gtf_to_gene_models.R

```
gtf_to_gene_models = function(gtf_path, save_path, ncores ){

  txdb = makeTxDbFromGFF(gtf_path, format="gtf")
  dropSeqlevels(txdb, "chrM")
  
  genes_list = genes(txdb)
  gene_names = names(genes_list)
  chrs = as.character(seqnames(genes_list))
  strands = as.character(strand(genes_list))
  exons_list_by_gene = exonsBy(txdb, by="gene")
  exons_list_by_tx = exonsBy(txdb, by="tx", use.names = TRUE)
  txs_list_by_gene = transcriptsBy(txdb, "gene")
  
  gene_models = mclapply(1:length(gene_names), function(geneid){
    #gene_models = lapply(1:2, function(geneid){
    exons = disjoin(exons_list_by_gene[[geneid]])
    exon_starts = start(exons)
    exon_ends = end(exons)
    exon_lens = width(exons)
    
    txs = txs_list_by_gene[[geneid]]
    txs_list = lapply(1:length(txs), function(ii){
      tx_name = txs$tx_name[ii]
      exons_in_tx = exons_list_by_tx[[tx_name]]
      index = sort(subjectHits(findOverlaps(exons_in_tx, exons)))
      unique(index)
    })
    names(txs_list) = txs$tx_name
    if (geneid %% 100 == 0) {print(geneid); gc()}
    
    return(list(chr = chrs[geneid], exonStarts = exon_starts,
                exonEnds = exon_ends, exonLens = exon_lens,
                txs = txs_list, exonNum = length(exon_starts), 
                txNum = length(txs_list),
                str = strands[geneid]))
    
  }, mc.cores = ncores)
  
  names(gene_models) = gene_names
  
  save(gene_models, file = save_path)
  return(0)
}
```
