## Supplemental Code for "AIDE: annotation-assisted isoform discovery with high precision": RcppExports.R

```
# Generated by using Rcpp::compileAttributes() -> do not edit by hand
# Generator token: 10BE3573-1514-4C36-9D1C-5A225CD40393

ifExonPairInTranscript <- function(exonPair, Iso_mat) {
    .Call('_AIDE_ifExonPairInTranscript', PACKAGE = 'AIDE', exonPair, Iso_mat)
}

est_alp_forward <- function(H_mat, J, init, nreads, iter) {
    .Call('_AIDE_est_alp_forward', PACKAGE = 'AIDE', H_mat, J, init, nreads, iter)
}

get_Loglike <- function(H_mat, alp_hat) {
    .Call('_AIDE_get_Loglike', PACKAGE = 'AIDE', H_mat, alp_hat)
}

forward_LRT <- function(maxSteps, Ind_check, Hmat, nreads, sig_fward, loglike_init) {
    .Call('_AIDE_forward_LRT', PACKAGE = 'AIDE', maxSteps, Ind_check, Hmat, nreads, sig_fward, loglike_init)
}

forward_LRT_v2 <- function(maxSteps, Ind_check, Hmat, nreads, sig_fward, loglike_init) {
    .Call('_AIDE_forward_LRT_v2', PACKAGE = 'AIDE', maxSteps, Ind_check, Hmat, nreads, sig_fward, loglike_init)
}

stepwise_LRT <- function(maxSteps, Ind_check, Hmat, nreads, sig_fward, sig_bward, loglike_init) {
    .Call('_AIDE_stepwise_LRT', PACKAGE = 'AIDE', maxSteps, Ind_check, Hmat, nreads, sig_fward, sig_bward, loglike_init)
}
```
