## Supplemental Code for "AIDE: annotation-assisted isoform discovery with high precision": rdata_to_gtf.R

```
rdata_to_gtf = function(tp_dir, gene_models, gtf_name, ncores){
  LRT_res = mclapply(1:length(gene_models), function(id){
    result = NULL
    #geneName = geneNames[id]
    #cgene = gene_models[[geneName]]
    cgene = gene_models[[id]]
    geneName = names(gene_models)[id]

    loadtry = try(load(paste0(tp_dir, "data/", id, ".RData")), silent = TRUE)
    if(class(loadtry) == "try-error") return(NULL)
    if(class(result) != "list") return(NULL)

    txs = result$txs
    txnames = names(cgene$txs)[match(result$Ind_check, result$Ind_annt)]
    txnames[is.na(txnames)] = paste("AIDE", id, 1:sum(is.na(txnames)), sep = ".")
    txnum = length(txs)
    tx_tb = lapply(1:txnum, function(j){
      tx = txs[[j]]
      type = c("transcript", rep("exon", length(tx)))
      start = cgene$exonStarts[tx]
      start = c(min(start), start)
      end = cgene$exonEnds[tx]
      end = c(max(end), end)

      # transcript_id = rep(txnames[j], length(tx))
      fpkm = round(result$rpkm[j], digits = 5)
      frac = round(result$alpha_check[j], digits = 5)
      tkey = paste0('gene_id "', geneName, '"; ',
                   'transcript_id "', txnames[j], '"; ',
                   'FPKM "', fpkm, '"; ', 'frac "', frac, '"; ')
      ekey = paste0('gene_id "', geneName, '"; ',
                   'transcript_id "', rep(txnames[j], length(tx)), '"; ',
                   'exon_number "', 1:length(tx), '"; ')
      key = c(tkey, ekey)

      summ = data.frame(chr = cgene$chr, source = "AIDE", type = type,
                        start = start, end = end, score = ".", strand = cgene$str,
                        frame = ".", key = key,
                        stringsAsFactors = FALSE)
      return(summ)
    })
    tx_tb = Reduce(rbind, tx_tb)
    gkey = paste0('gene_id "', geneName, '"; ',
                  'FPKM "', result$rpkm_gene, '"; ')
    geneline = data.frame(chr = cgene$chr, source = "AIDE", type = "gene",
                          start = cgene$exonStarts[1], end = cgene$exonEnds[cgene$exonNum],
                          score = ".", strand = cgene$str, frame = ".", key = gkey,
                          stringsAsFactors = FALSE)
    dat = rbind(geneline, tx_tb)
    return(dat)
  })
  LRT_res = Reduce(rbind, LRT_res)
  write.table(LRT_res, file = gtf_name, sep = "\t",
              row.names = FALSE, col.names = FALSE, quote = FALSE)
  return(0)
}
```
