## Supplemental Code for "AIDE: annotation-assisted isoform discovery with high precision": stepwise_LRT_wrap_detailed_notruth.R

```
# wrap up stepwise LRT and evaluation in one function
# output: list(precision, recall, mse, total variation)
stepwise_LRT_wrap = function(J_annt, Ind_annt, Hmat, nExon, nreads, 
                             sig_fward = 0.05, sig_bward = 0.01){
  # tick = proc.time()
  if (J_annt == 1){
    Ind_check = Ind_annt
    alp = 1
    loglike_null = get_Loglike(Hmat[, Ind_annt, drop = FALSE], alp_hat = alp)
  }else{
    alp = 1
    loglike_init = sapply(Ind_annt, function(k){
      return( get_Loglike(Hmat[, k, drop = FALSE], alp_hat = alp) )
    })
    loglike_null = max(loglike_init)
    Ind_check_annt = which.max(loglike_init)
    Ind_check_annt = stepwise_LRT(maxSteps = 3*nExon, Ind_check_annt, Hmat[, Ind_annt, drop = FALSE], 
                                 nreads, sig_fward, sig_bward, loglike_init = loglike_null)
    Ind_check = Ind_annt[Ind_check_annt]
    
    init = rep(1/length(Ind_check), length(Ind_check))
    alpha_check = est_alp_forward(H_mat = Hmat[, Ind_check, drop = FALSE], J = length(Ind_check), init, nreads, iter = 500)
    loglike_null = get_Loglike(Hmat[, Ind_check, drop = FALSE], alp_hat = alpha_check)
  }
  
  if (ncol(Hmat) > J_annt){
    Ind_check = stepwise_LRT(maxSteps = 3*nExon, Ind_check, Hmat, nreads, sig_fward, sig_bward,
                             loglike_init = loglike_null)
    init = rep(1/length(Ind_check), length(Ind_check))
    alpha_check = est_alp_forward(H_mat = Hmat[, Ind_check, drop = FALSE], J = length(Ind_check), init, nreads, iter = 500)
  }

  return(list(Ind_check = Ind_check, alpha_check = alpha_check))
}


stepwise_LRT_wrap_single = function(Ind_annt){
  Ind_check = Ind_annt
  alpha_check = 1
  return(list(Ind_check = Ind_check, alpha_check = alpha_check))
}
```
