## Supplemental Code for "AIDE: annotation-assisted isoform discovery with high precision": README.md

AIDE: annotation-assisted isoform discovery and quantification from RNA-seq data
================
Wei Vivian Li, Jingyi Jessica Li
2018-11-12

<!-- README.md is generated from README.Rmd. Please edit that file -->
Latest News
-----------

> 2018/11/12:

-   Version 0.0.2 is released!

> 2018/04/08:

-   Version 0.0.1 is released!

Introduction
------------

AIDE is a statistical method which identifies full-length mRNA isoforms from a novel perspective: using the likelihood ratio test to find novel isoforms in a stepwise manner given annotated isoforms, by prioritizing and selectively borrowing information from the annotated isoforms. Please refer to our preprint at [Biorxiv](https://www.biorxiv.org/content/early/2018/10/07/437350) for detailed description of our method.

Any suggestions on the package are welcome! For technical problems, please report to [Issues](https://github.com/Vivianstats/AID/issues). For suggestions and comments on the method, please contact Wei (<>) or Dr. Jessica Li (<>).

Installation
------------

You can install `AIDE` from github with:

``` r
# install.packages("devtools")
devtools::install_github("Vivianstats/AIDE")
```

Quick start
-----------

`aid` requires three input files:

-   The GTF file of the genome annotation;
-   The BAM file of the RNA-seq sample. The BAM file should be sorted and the index BAI file should be present in the same directory as the BAM file;
-   The FASTA file of the genome sequences.

The final output of `aide` is a GTF file named "transcripts.gtf", which contains the reconstructed transcripts and their corresponding abudance levels. The package has been tested using the [GENCODE annotation](https://www.gencodegenes.org/releases/24.html). This is a basic example which shows how to use the `aide` function.

``` r
aide(gtf_path = "./hg19.gtf",     #full path of the GTF file
    bam_path = "./example.bam",  #full path of the BAM file
    fasta_path = "./hg19.fa",    #full path of the FASTA file
    out_dir = "./",              #output directory of temporary and filnal results
    readLen = 100,               #read length used to calculate ioform effective length
    strandmode = 0,              #library type of the RNA-seq sample
    ncores = 20                  #number of cores used for parallel computation 
    )
```

Please refer to the package [manual](https://github.com/Vivianstats/AIDE/blob/master/inst/docs/) for a full list of arguments and detailed usage.
