## Supplemental Code for "AIDE: annotation-assisted isoform discovery with high precision": calculate_h_supp.cpp

```
// [[Rcpp::depends(RcppArmadillo)]]
#include <RcppArmadillo.h>
using namespace Rcpp;
using namespace arma;

// This code is to speed up the bottleneck part in
// R function impute_fragment_length()
// It calculates ifExonPairInTranscript.typeII

// [[Rcpp::export]]
arma::imat ifExonPairInTranscript(arma::imat exonPair, arma::imat Iso_mat){
  int npair = exonPair.n_rows;
//  Rcout << npair << endl;
  int ntx = Iso_mat.n_rows;
//  Rcout << ntx << endl;
  arma::imat result(npair, ntx);
  result.ones();
  for (int i=0; i<npair; i++){
//    Rcout << "i:" <<i << endl;
    for (int j=0; j<ntx; j++){
//      Rcout << "j:" <<j << endl;
      for (int k=0; k<4; k++){
        int crr = exonPair(i,k);
        if (Iso_mat(j, crr-1) == 0){
          result(i, j) = 0;
//          print(wrap(result));
          continue;
          }
      }
    }
  }
  return(result);
}
```
