## Supplemental Code for "AIDE: annotation-assisted isoform discovery with high precision": estimation.cpp

```
// [[Rcpp::depends(RcppArmadillo)]]
#include <RcppArmadillo.h>
using namespace Rcpp;
using namespace arma;

// These functions are used to speed up the forward likelihood ratio test
// V1: only include functions for forward LRT
// current: add backward steps


// [[Rcpp::export]]
arma::vec est_alp_forward(arma::mat H_mat, int J, arma::vec init, double nreads, int iter) {
  // estimate alpha given H matrix
  arma::vec alp_crr = init;
  int ii = 0;
  while(ii < iter){
    arma::vec denom = H_mat * alp_crr;
//    denom.elem( find(denom == 0.0) ) = 1e300;
    arma::vec null = denom.elem(find(denom == 0));
//    int nnull = null.n_elem;
    denom.replace(0, 1e300);  // replace each NaN with 0
    arma::vec alp_new;
    alp_new.set_size(J);
    for (int j=0; j<J; j++){
      arma::vec numer = H_mat.col(j) * alp_crr(j);
      arma::vec tp = numer / denom;
      alp_new(j) = sum(tp)/ (nreads - null.n_elem);
    }

    double error = sum(square(alp_new - alp_crr));
    if (error < 1e-8){
      break;
    }else{
      alp_crr = alp_new;
    }
    ii ++;
  }
  return (alp_crr);
}


// [[Rcpp::export]]
double get_Loglike(arma::mat H_mat, arma::vec alp_hat){
  arma::vec ltp = log(H_mat * alp_hat);
  //print(wrap(ltp));
  ltp.replace(-datum::inf, -1e10);
  //ltp.elem( find(tp == -Inf)) += exp(-1e10);
  //print(wrap(ltp));
  return(sum(ltp));
}


// [[Rcpp::export]]
arma::uvec forward_LRT(int maxSteps, arma::uvec Ind_check, arma::mat Hmat, int nreads,
                 float sig_fward, double loglike_init){
  int steps = 0;
  //int n = nExon;
  // int JJ = pow(2, n) - 1;
  int JJ = Hmat.n_cols;
  int ncrr = Ind_check.n_elem;
  double loglike_null = loglike_init;
  while(steps < maxSteps){

    arma::vec loglike_forward(JJ);
    // look for the next best candidate
    for (int k=0; k<JJ; k++){
      // if the new candidate is already checked, skip it
      if (any(Ind_check == k+1)){
        loglike_forward(k) = -1e50;
        continue;
      }
      // current candidates
      arma::uvec Ind_crr(ncrr + 1);
      for (int ii=0; ii<ncrr; ii++){
        Ind_crr(ii) = Ind_check(ii);
      }
      Ind_crr(ncrr) = k+1;
      arma::vec init(ncrr + 1);
      init.ones();
      init = init/(ncrr + 1);
      arma::vec alp_curr = est_alp_forward(Hmat.cols(Ind_crr-1), ncrr+1,
                                     init, (double)nreads, 500);
      //print(wrap(alp_curr));
      double like_curr = get_Loglike(Hmat.cols(Ind_crr-1), alp_curr);
      loglike_forward(k) = like_curr;
    }

    int Ind_newiso = loglike_forward.index_max() + 1;
    double loglike_altive = max(loglike_forward);
    if (loglike_altive <= loglike_null){break;}
    // uvec check = find(loglike_forward == loglike_altive);
    // if (check.n_elem > 1){
    //   Rcout << "max values:" << check.n_elem << endl;}
    double Q = -2 * (loglike_null - loglike_altive);
    //Rcout << Q << endl;
    arma::vec QQ(1);
    QQ(0) = Q;
    NumericVector p_val = 1-pchisq(as<NumericVector>(wrap(QQ)), 1.0);
    //Rcout << p_val(0) << endl;
    // add newly discovered isoform to answers if pass LRT
    if (p_val(0) <= sig_fward){
      //Rcout << "step:" << steps << endl;
      ncrr ++;
      //print(wrap(Ind_check));
      Ind_check.resize(ncrr);
      //print(wrap(Ind_check));
      Ind_check(ncrr-1) = Ind_newiso;

      arma::vec init(ncrr);
      init.ones();
      init = init/ncrr;
      arma::vec alp = est_alp_forward(Hmat.cols(Ind_check-1), ncrr,
                                init, nreads, 500);
      //print(wrap(alp));
      loglike_null = get_Loglike(Hmat.cols(Ind_check-1), alp);
      //print(wrap(loglike_null));
    }else{
      break; }
    steps ++;

  }
  return(Ind_check);
}


// [[Rcpp::export]]
arma::uvec forward_LRT_v2(int maxSteps, arma::uvec Ind_check, arma::mat Hmat, int nreads,
                 float sig_fward, double loglike_init){
  int steps = 0;
  //int n = nExon;
  //int JJ = pow(2, n) - 1;
  int JJ = Hmat.n_cols;
  int ncrr = Ind_check.n_elem;
  double loglike_null = loglike_init;
  while(steps < maxSteps){

    vec loglike_forward(JJ);
    // look for the next best candidate
    for (int k=0; k<JJ; k++){
      // if the new candidate is already checked, skip it
      if (any(Ind_check == k+1)){
        loglike_forward(k) = -1e50;
        continue;
      }
      // current candidates
      arma::uvec Ind_crr(ncrr + 1);
      for (int ii=0; ii<ncrr; ii++){
        Ind_crr(ii) = Ind_check(ii);
      }
      Ind_crr(ncrr) = k+1;
      arma::vec init(ncrr + 1);
      init.ones();
      init = init/(ncrr + 1);
      arma::vec alp_curr = est_alp_forward(Hmat.cols(Ind_crr-1), ncrr+1,
                                     init, (double)nreads, 500);
      //print(wrap(alp_curr));
      double like_curr = get_Loglike(Hmat.cols(Ind_crr-1), alp_curr);
      loglike_forward(k) = like_curr;
    }

    double loglike_altive = max(loglike_forward);
    arma::uvec Ind_newiso = find(loglike_forward == loglike_altive) + 1;

    double Q = -2 * (loglike_null - loglike_altive);
    //Rcout << Q << endl;
    arma::vec QQ(1);
    QQ(0) = Q;
    NumericVector p_val = 1-pchisq(as<NumericVector>(wrap(QQ)), Ind_newiso.n_elem);
    //Rcout << p_val(0) << endl;
    // add newly discovered isoform to answers if pass LRT
    if (p_val(0) <= sig_fward){
      //Rcout << "step:" << steps << endl;
      ncrr = ncrr + Ind_newiso.n_elem;
      Ind_check.insert_rows(Ind_check.n_elem, Ind_newiso);

      arma::vec init(ncrr);
      init.ones();
      init = init/ncrr;
      arma::vec alp = est_alp_forward(Hmat.cols(Ind_check-1), ncrr,
                                init, nreads, 500);
      //print(wrap(alp));
      loglike_null = get_Loglike(Hmat.cols(Ind_check-1), alp);
      //print(wrap(loglike_null));
    }else{
      break; }
    steps ++;

  }
  return(Ind_check);
}


// [[Rcpp::export]]
arma::uvec stepwise_LRT(int maxSteps, arma::uvec Ind_check, arma::mat Hmat, int nreads,
                float sig_fward, float sig_bward, double loglike_init){
  int steps = 0;
  //int n = nExon;
  //int JJ = pow(2, n) - 1;
  int JJ = Hmat.n_cols;
  int ncrr = Ind_check.n_elem;
  double loglike_null = loglike_init;
  while(steps < maxSteps){

    arma::vec loglike_forward(JJ);
    // look for the next best candidate
    for (int k=0; k<JJ; k++){
      //Rcout << k << endl;
      // if the new candidate is already checked, skip it
      if (any(Ind_check == k+1)){
        loglike_forward(k) = -1e200;
        continue;
      }
      // current candidates
      arma::uvec Ind_crr(ncrr + 1);
      for (int ii=0; ii<ncrr; ii++){
        Ind_crr(ii) = Ind_check(ii);
      }
      Ind_crr(ncrr) = k+1;
      arma::vec init(ncrr + 1);
      init.ones();
      init = init/(ncrr + 1);
      arma::vec alp_curr = est_alp_forward(Hmat.cols(conv_to< uvec >::from(Ind_crr-1)), ncrr+1,
                                    init, (double)nreads, 500);
      //print(wrap(alp_curr));
      double like_curr = get_Loglike(Hmat.cols(conv_to< uvec >::from(Ind_crr-1)), alp_curr);
      loglike_forward(k) = like_curr;
    }
    //Rcout << "check1" << endl;
    int Ind_newiso = loglike_forward.index_max() + 1;
    //print(wrap(Ind_newiso));
    double loglike_altive = max(loglike_forward);
    if (loglike_altive <= loglike_null){break;}

    double Q = -2 * (loglike_null - loglike_altive);
    arma::vec QQ(1);
    QQ(0) = Q;
    NumericVector p_val_fward = 1-pchisq(as<NumericVector>(wrap(QQ)), 1.0);

    // add newly discovered isoform to answers if pass LRT
    if (p_val_fward(0) <= sig_fward){
       //Rcout << "forward" << endl;
       ncrr ++;
       Ind_check.resize(ncrr);
       Ind_check(ncrr-1) = Ind_newiso;
       //print(wrap(Ind_check));
       arma::vec init(ncrr);
       init.ones();
       init = init/ncrr;
       arma::vec alp = est_alp_forward(Hmat.cols(conv_to< uvec >::from(Ind_check-1)), ncrr,
                            init, nreads, 500);

       loglike_null = get_Loglike(Hmat.cols(conv_to< uvec >::from(Ind_check-1)), alp);
       //Rcout << loglike_null << endl;

       // only consider move backward, after move one step forward
       arma::vec loglike_bward(ncrr);
       for (int kk = 0; kk < ncrr; kk++){
         arma::uvec Ind_crr = Ind_check;
         Ind_crr.shed_row(kk);

         arma::vec init(ncrr-1);
         init.ones();
         init = init/(ncrr - 1);
         arma::vec alp_curr = est_alp_forward(Hmat.cols(Ind_crr-1), ncrr-1,
                                        init, (double)nreads, 500);
         double like_curr = get_Loglike(Hmat.cols(Ind_crr-1), alp_curr);
         loglike_bward(kk) = like_curr;
       }
       Ind_newiso = loglike_bward.index_max() + 1;
       loglike_altive = max(loglike_bward);
       Q = -2 * (loglike_altive - loglike_null);
       QQ(0) = Q;
       NumericVector p_val_bward = 1-pchisq(as<NumericVector>(wrap(QQ)), 1.0);
       if (p_val_bward(0) > sig_bward){
         //Rcout << "backward" << endl;
         Ind_check.shed_row(Ind_newiso - 1);
         ncrr --;
         //print(wrap(Ind_check));
         init.resize(ncrr);
         init.ones();
         init = init/(ncrr);

         alp = est_alp_forward(Hmat.cols(Ind_check-1), ncrr,
                              init, nreads, 500);
         loglike_null = get_Loglike(Hmat.cols(Ind_check-1), alp);
         //Rcout << loglike_null << endl;
       }
    }else{
      break; }
    steps ++;

  }
  return(Ind_check);
}
```
