## Supplemental Code for "AIDE: annotation-assisted isoform discovery with high precision": RcppExports.cpp

```
// Generated by using Rcpp::compileAttributes() -> do not edit by hand
// Generator token: 10BE3573-1514-4C36-9D1C-5A225CD40393

#include <RcppArmadillo.h>
#include <Rcpp.h>

using namespace Rcpp;

// ifExonPairInTranscript
arma::imat ifExonPairInTranscript(arma::imat exonPair, arma::imat Iso_mat);
RcppExport SEXP _AIDE_ifExonPairInTranscript(SEXP exonPairSEXP, SEXP Iso_matSEXP) {
BEGIN_RCPP
    Rcpp::RObject rcpp_result_gen;
    Rcpp::RNGScope rcpp_rngScope_gen;
    Rcpp::traits::input_parameter< arma::imat >::type exonPair(exonPairSEXP);
    Rcpp::traits::input_parameter< arma::imat >::type Iso_mat(Iso_matSEXP);
    rcpp_result_gen = Rcpp::wrap(ifExonPairInTranscript(exonPair, Iso_mat));
    return rcpp_result_gen;
END_RCPP
}
// est_alp_forward
arma::vec est_alp_forward(arma::mat H_mat, int J, arma::vec init, double nreads, int iter);
RcppExport SEXP _AIDE_est_alp_forward(SEXP H_matSEXP, SEXP JSEXP, SEXP initSEXP, SEXP nreadsSEXP, SEXP iterSEXP) {
BEGIN_RCPP
    Rcpp::RObject rcpp_result_gen;
    Rcpp::RNGScope rcpp_rngScope_gen;
    Rcpp::traits::input_parameter< arma::mat >::type H_mat(H_matSEXP);
    Rcpp::traits::input_parameter< int >::type J(JSEXP);
    Rcpp::traits::input_parameter< arma::vec >::type init(initSEXP);
    Rcpp::traits::input_parameter< double >::type nreads(nreadsSEXP);
    Rcpp::traits::input_parameter< int >::type iter(iterSEXP);
    rcpp_result_gen = Rcpp::wrap(est_alp_forward(H_mat, J, init, nreads, iter));
    return rcpp_result_gen;
END_RCPP
}
// get_Loglike
double get_Loglike(arma::mat H_mat, arma::vec alp_hat);
RcppExport SEXP _AIDE_get_Loglike(SEXP H_matSEXP, SEXP alp_hatSEXP) {
BEGIN_RCPP
    Rcpp::RObject rcpp_result_gen;
    Rcpp::RNGScope rcpp_rngScope_gen;
    Rcpp::traits::input_parameter< arma::mat >::type H_mat(H_matSEXP);
    Rcpp::traits::input_parameter< arma::vec >::type alp_hat(alp_hatSEXP);
    rcpp_result_gen = Rcpp::wrap(get_Loglike(H_mat, alp_hat));
    return rcpp_result_gen;
END_RCPP
}
// forward_LRT
arma::uvec forward_LRT(int maxSteps, arma::uvec Ind_check, arma::mat Hmat, int nreads, float sig_fward, double loglike_init);
RcppExport SEXP _AIDE_forward_LRT(SEXP maxStepsSEXP, SEXP Ind_checkSEXP, SEXP HmatSEXP, SEXP nreadsSEXP, SEXP sig_fwardSEXP, SEXP loglike_initSEXP) {
BEGIN_RCPP
    Rcpp::RObject rcpp_result_gen;
    Rcpp::RNGScope rcpp_rngScope_gen;
    Rcpp::traits::input_parameter< int >::type maxSteps(maxStepsSEXP);
    Rcpp::traits::input_parameter< arma::uvec >::type Ind_check(Ind_checkSEXP);
    Rcpp::traits::input_parameter< arma::mat >::type Hmat(HmatSEXP);
    Rcpp::traits::input_parameter< int >::type nreads(nreadsSEXP);
    Rcpp::traits::input_parameter< float >::type sig_fward(sig_fwardSEXP);
    Rcpp::traits::input_parameter< double >::type loglike_init(loglike_initSEXP);
    rcpp_result_gen = Rcpp::wrap(forward_LRT(maxSteps, Ind_check, Hmat, nreads, sig_fward, loglike_init));
    return rcpp_result_gen;
END_RCPP
}
// forward_LRT_v2
arma::uvec forward_LRT_v2(int maxSteps, arma::uvec Ind_check, arma::mat Hmat, int nreads, float sig_fward, double loglike_init);
RcppExport SEXP _AIDE_forward_LRT_v2(SEXP maxStepsSEXP, SEXP Ind_checkSEXP, SEXP HmatSEXP, SEXP nreadsSEXP, SEXP sig_fwardSEXP, SEXP loglike_initSEXP) {
BEGIN_RCPP
    Rcpp::RObject rcpp_result_gen;
    Rcpp::RNGScope rcpp_rngScope_gen;
    Rcpp::traits::input_parameter< int >::type maxSteps(maxStepsSEXP);
    Rcpp::traits::input_parameter< arma::uvec >::type Ind_check(Ind_checkSEXP);
    Rcpp::traits::input_parameter< arma::mat >::type Hmat(HmatSEXP);
    Rcpp::traits::input_parameter< int >::type nreads(nreadsSEXP);
    Rcpp::traits::input_parameter< float >::type sig_fward(sig_fwardSEXP);
    Rcpp::traits::input_parameter< double >::type loglike_init(loglike_initSEXP);
    rcpp_result_gen = Rcpp::wrap(forward_LRT_v2(maxSteps, Ind_check, Hmat, nreads, sig_fward, loglike_init));
    return rcpp_result_gen;
END_RCPP
}
// stepwise_LRT
arma::uvec stepwise_LRT(int maxSteps, arma::uvec Ind_check, arma::mat Hmat, int nreads, float sig_fward, float sig_bward, double loglike_init);
RcppExport SEXP _AIDE_stepwise_LRT(SEXP maxStepsSEXP, SEXP Ind_checkSEXP, SEXP HmatSEXP, SEXP nreadsSEXP, SEXP sig_fwardSEXP, SEXP sig_bwardSEXP, SEXP loglike_initSEXP) {
BEGIN_RCPP
    Rcpp::RObject rcpp_result_gen;
    Rcpp::RNGScope rcpp_rngScope_gen;
    Rcpp::traits::input_parameter< int >::type maxSteps(maxStepsSEXP);
    Rcpp::traits::input_parameter< arma::uvec >::type Ind_check(Ind_checkSEXP);
    Rcpp::traits::input_parameter< arma::mat >::type Hmat(HmatSEXP);
    Rcpp::traits::input_parameter< int >::type nreads(nreadsSEXP);
    Rcpp::traits::input_parameter< float >::type sig_fward(sig_fwardSEXP);
    Rcpp::traits::input_parameter< float >::type sig_bward(sig_bwardSEXP);
    Rcpp::traits::input_parameter< double >::type loglike_init(loglike_initSEXP);
    rcpp_result_gen = Rcpp::wrap(stepwise_LRT(maxSteps, Ind_check, Hmat, nreads, sig_fward, sig_bward, loglike_init));
    return rcpp_result_gen;
END_RCPP
}

static const R_CallMethodDef CallEntries[] = {
    {"_AIDE_ifExonPairInTranscript", (DL_FUNC) &_AIDE_ifExonPairInTranscript, 2},
    {"_AIDE_est_alp_forward", (DL_FUNC) &_AIDE_est_alp_forward, 5},
    {"_AIDE_get_Loglike", (DL_FUNC) &_AIDE_get_Loglike, 2},
    {"_AIDE_forward_LRT", (DL_FUNC) &_AIDE_forward_LRT, 6},
    {"_AIDE_forward_LRT_v2", (DL_FUNC) &_AIDE_forward_LRT_v2, 6},
    {"_AIDE_stepwise_LRT", (DL_FUNC) &_AIDE_stepwise_LRT, 7},
    {NULL, NULL, 0}
};

RcppExport void R_init_AIDE(DllInfo *dll) {
    R_registerRoutines(dll, NULL, CallEntries, NULL, NULL);
    R_useDynamicSymbols(dll, FALSE);
}
```
