## Supplemental Figures for "AIDE: annotation-assisted isoform discovery with high precision"

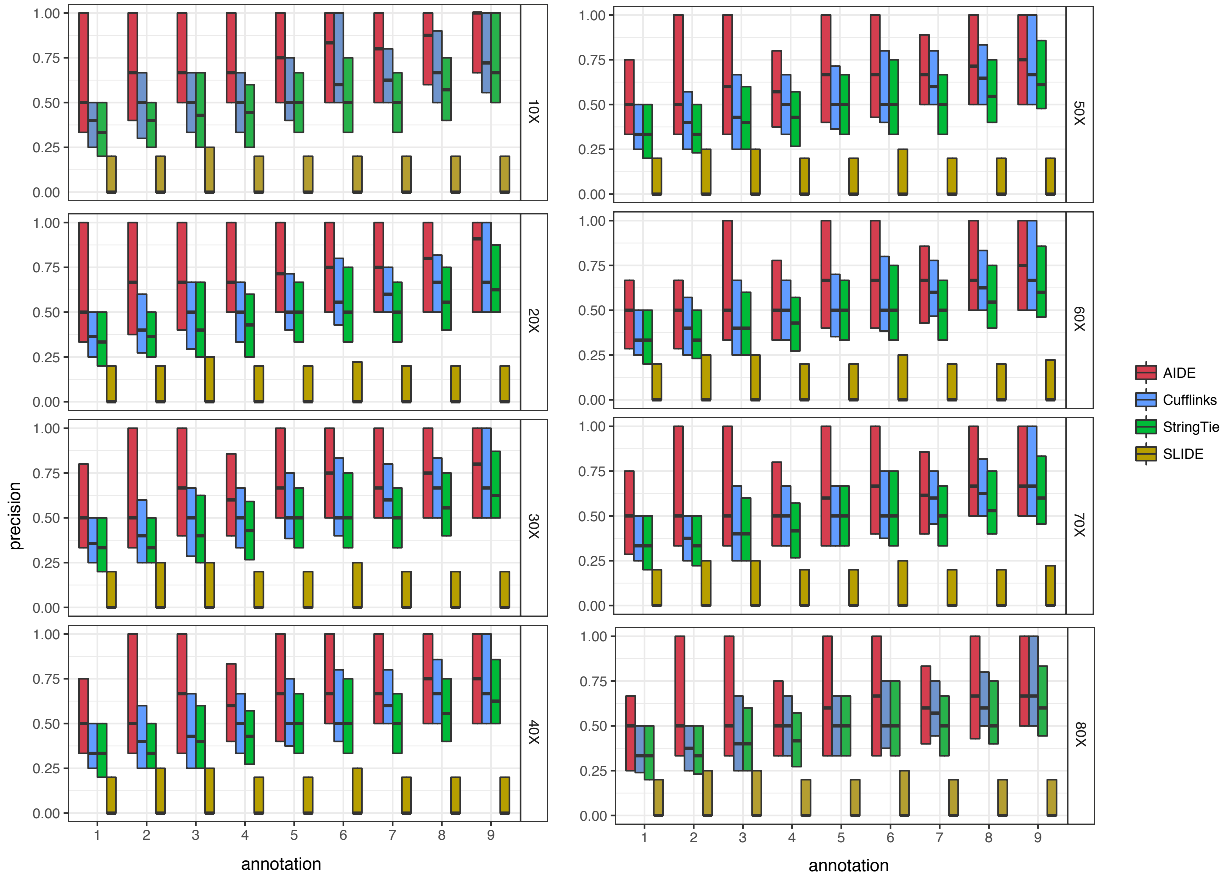


**Figure S1:** Gene-level precision rates of AIDE and the other three isoform discovery methods in simulation. Each box gives the 1st quantile, median, and 3rd quantile of the gene-level precision rates given the corresponding synthetic annotation set and read coverage.

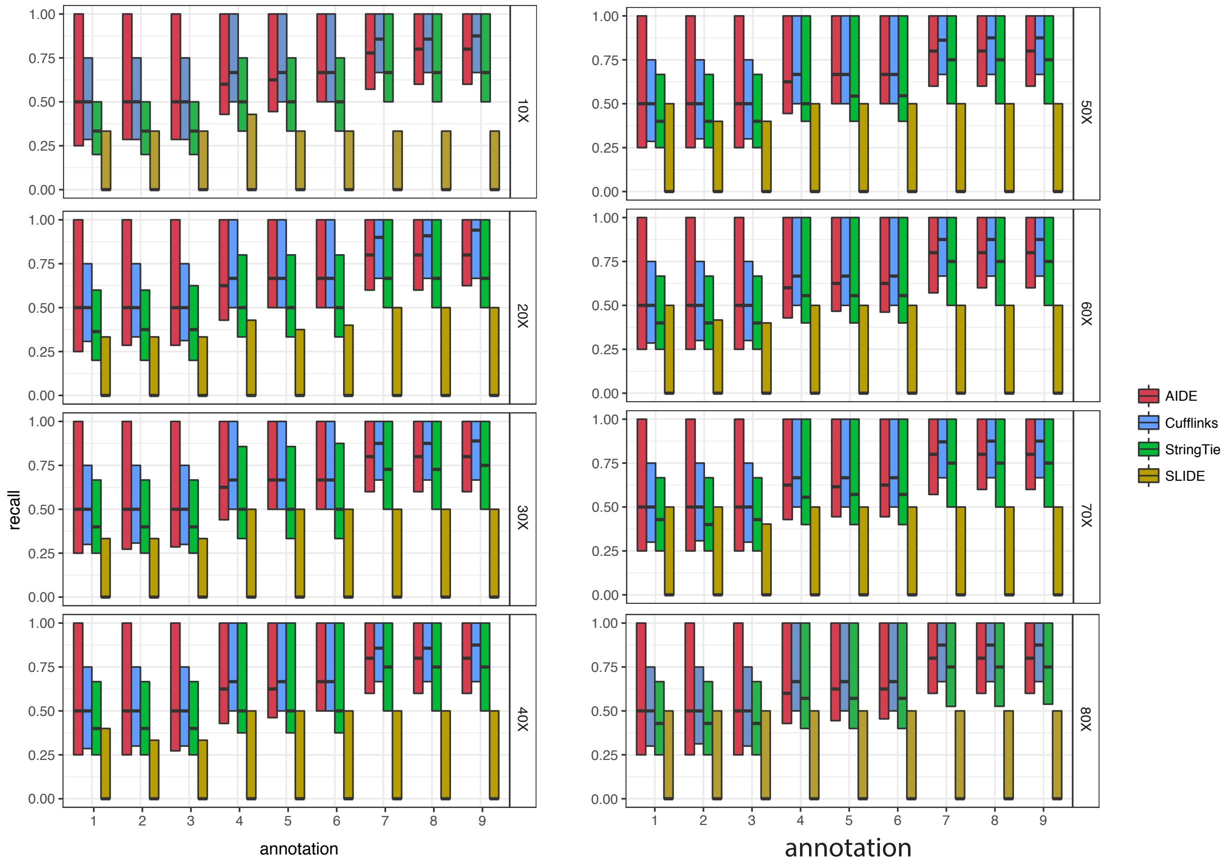


**Figure S2:** Gene-level recall rates of AIDE and the other three isoform discovery methods in simulation. Each box gives the 1st quantile, median, and 3rd quantile of the gene-level recall rates given the corresponding synthetic annotation set and read coverage.

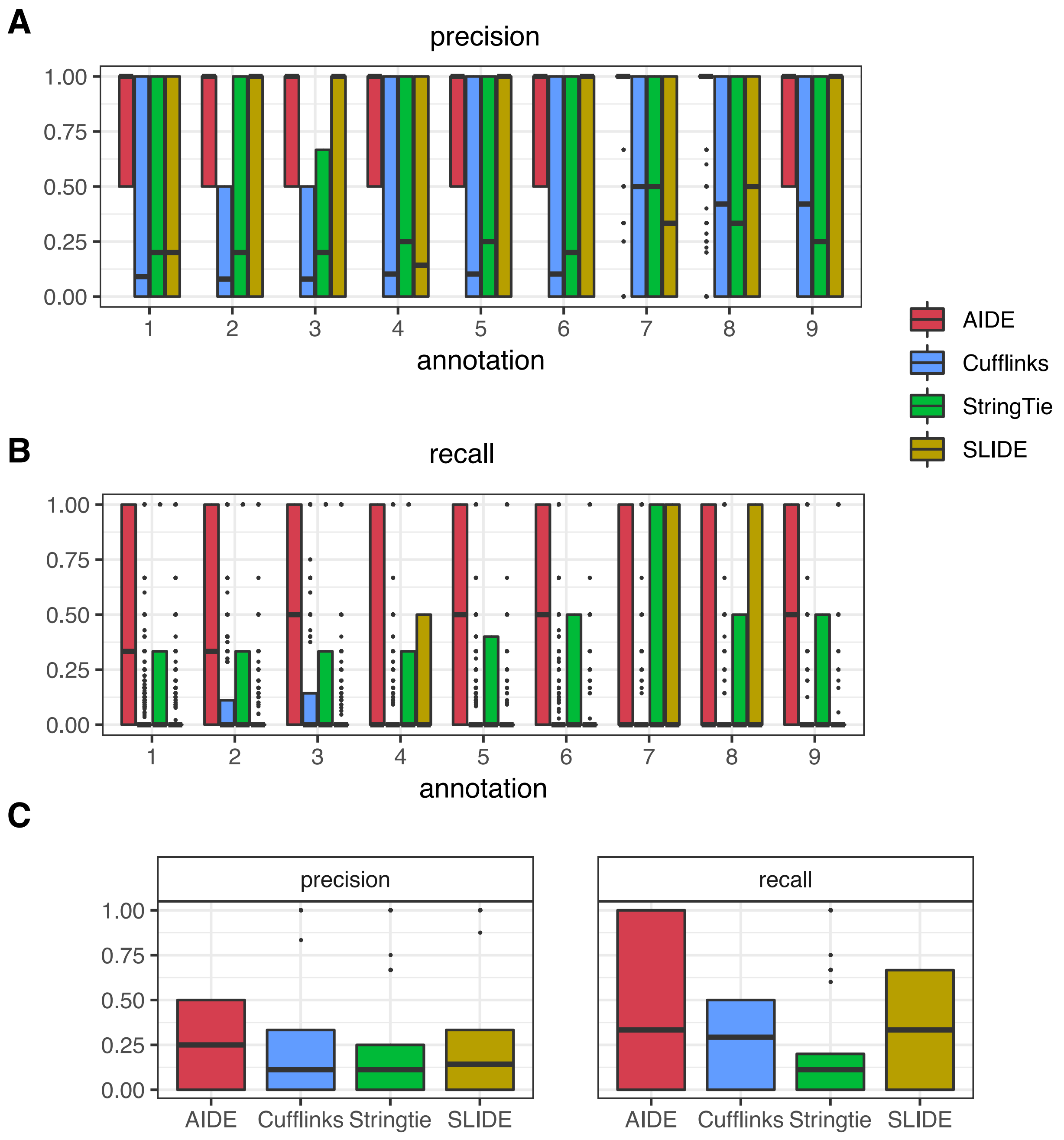


**Figure S3:** Gene-level accuracy of AIDE and the other three isoform discovery methods in simulation. **a-b**: Accuracy was calculated based on expressed but non-annotated isoforms. Each boxplot gives the 1st quantile, median, and 3rd quantile of the gene-level precision or recall rates given the corresponding synthetic annotation set and $10$x read coverage. Genes for which none of the methods could correctly identify any isoforms were excluded from the plots. **c**: Precision and recall rates were calculated based on expressed isoforms. No annotated isoforms were given as prior information, but exon boundaries were given to the four discovery methods. Genes for which none of the methods could correctly identify any isoforms were excluded from the plots.

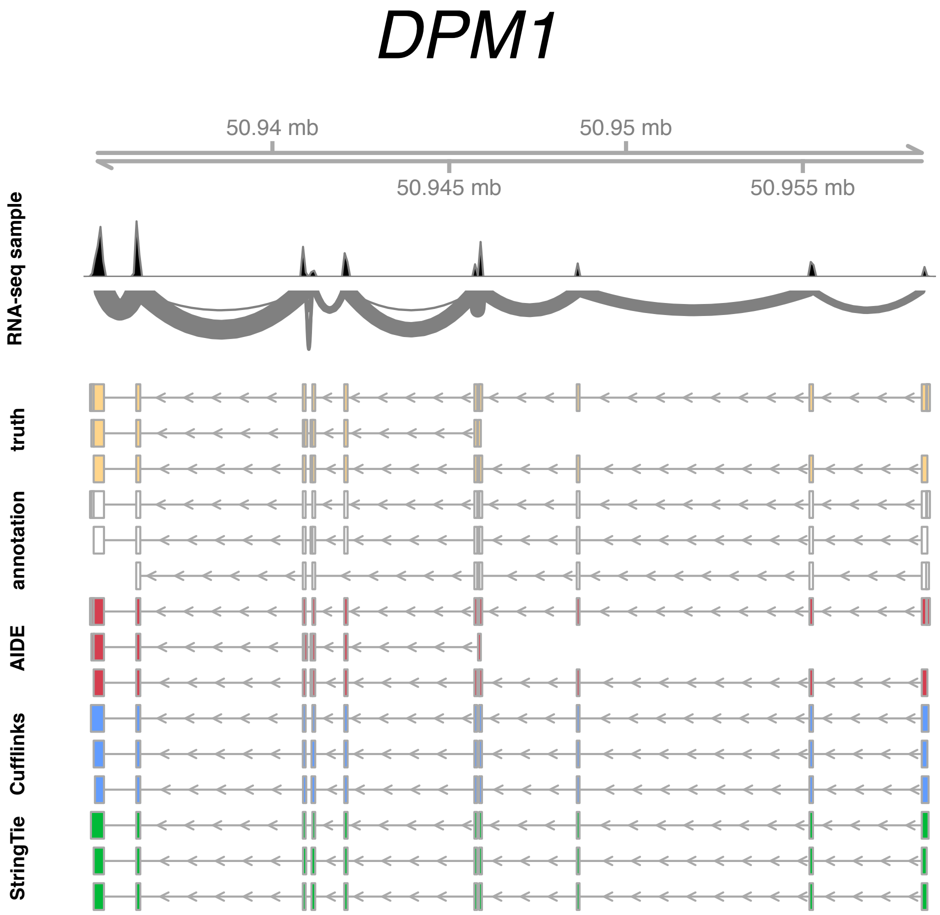


**Figure S4:** Isoform discovery for the human gene *DPM1* with the syethetic annotation set 1. The histogram and the sashimi plot denote the RNA-seq reads mapped to the gene *DPM1* in the RNA-seq data simulated by the `polyester` R package. The annotation (white) for this gene has a 67% purity and a 67% completeness, compared with the truly expressed isoforms (yellow). AIDE, Cufflinks, and StringTie each discovers three isoforms, but only AIDE is able to identify the shortest isoform missing in the annotation. SLIDE reports 17 isoforms, which are not displayed in the plot.

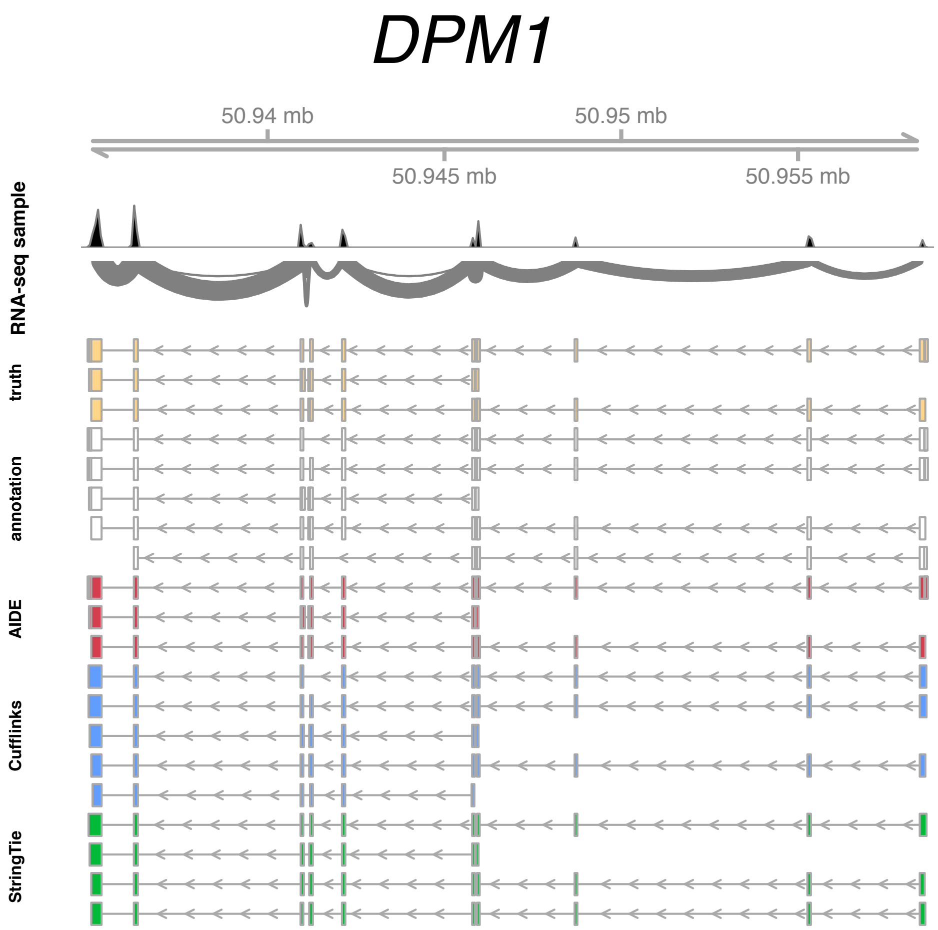


**Figure S5:** Isoform discovery for the human gene *DPM1* with the syethetic annotation set 9. The histogram and the sashimi plot denote the RNA-seq reads mapped to the gene *DPM1* in the RNA-seq data simulated by the `polyester` R package. The annotation (white) for this gene has a 60% purity and a 100% completeness, compared with the truly expressed isoforms (yellow). AIDE, Cufflinks, and StringTie respectively discovers three, five, and four isoforms, and only AIDE is able to identify the three true isoforms with 100% accuracy. SLIDE reports $20$ isoforms, which are not displayed in the plot.

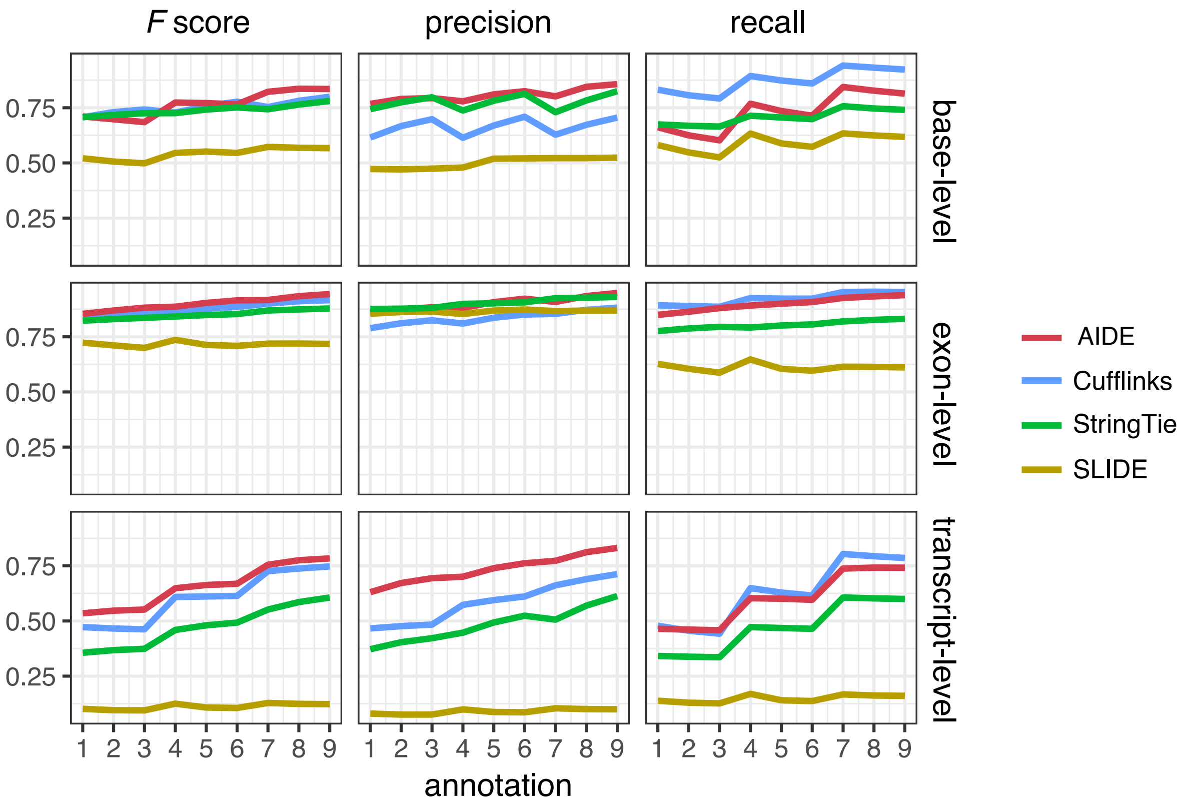


**Figure S6:** Comparison between AIDE and the other three isoform discovery methods in simulation. What is displayed is the genome-wide average performance of AIDE, Cufflinks, StringTie, and SLIDE given each of the nine synthetic annotation sets. The base-level, exon-level, and isoform-level precision rates, recall rates, and $F$ scores averaged across the human genes are calculated based on RNA-seq data with a $10$x coverage.

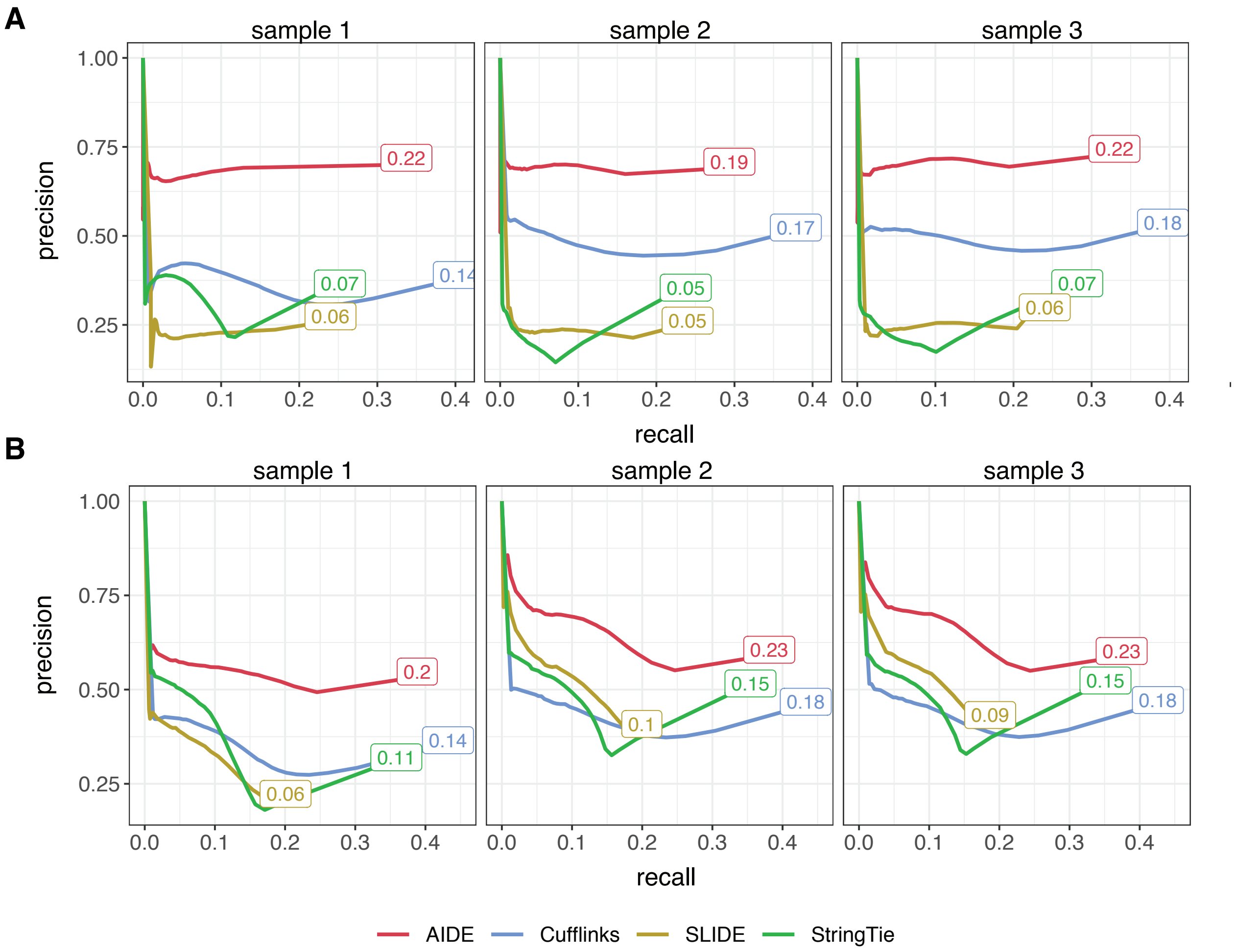


**Figure S7:** Comparison between AIDE and the other three isoform discovery methods in real data studies. We applied AIDE, Cufflinks, StringTie, and SLIDE to isoform discovery on three human ESC samples (**a**) and three mouse BMDM samples (**b**). The estimated expression levels of the predicted isoforms were then summarized in the FPKM unit. The precision-recall curves (at isoform-level) were obtained by thresholding the FPKM values of the predicted isoforms. The corresponding AUC of each method is also marked in the plot.

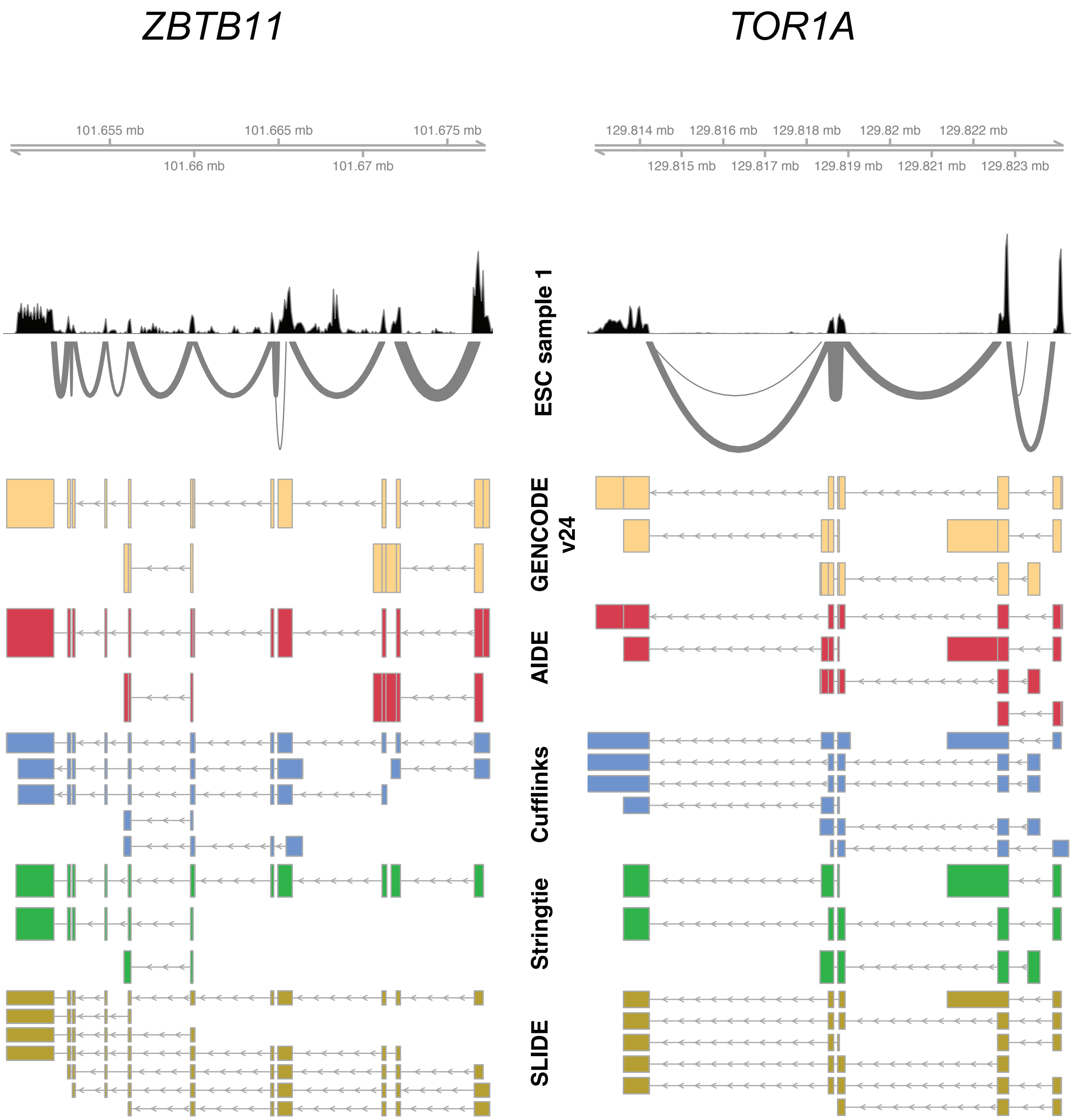


**Figure S8:** Isoform discovery for the human genes *ZBTB11* and *TOR1A* based on real data. The histogram and the sashimi plot denote the RNA-seq reads mapped to the two genes in the human ESC sample 1 (Table S1). Isoform discovery is based on the GENCODE human annotation version 24. AIDE achieves the best accuracy among the four methods.

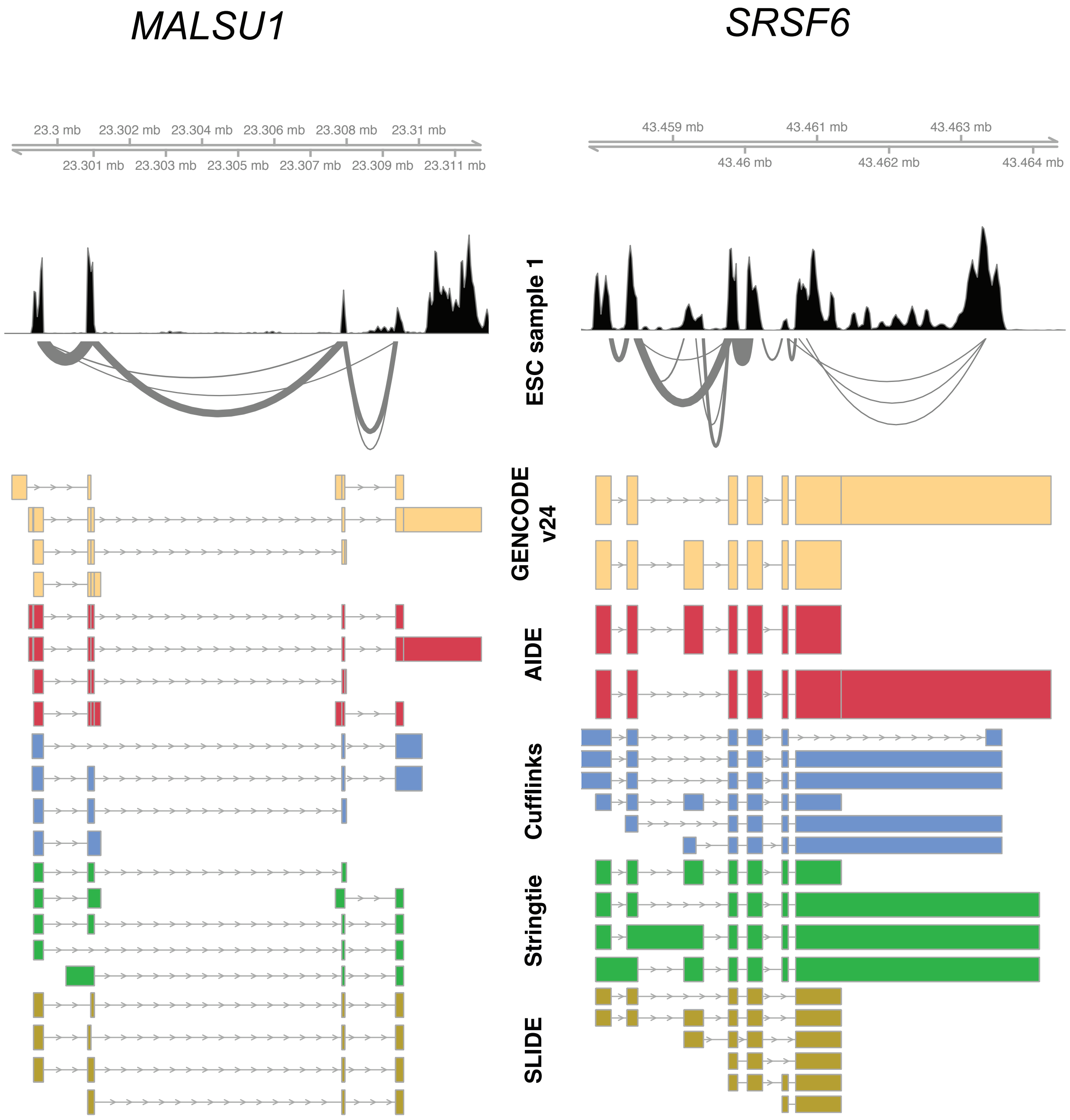


**Figure S9:** Isoform discovery for the human genes *MALSU1* and *SRSF6* based on real data. The histogram and the sashimi plot denote the RNA-seq reads mapped to the two genes in the human ESC sample 1 (Table S1). Isoform discovery is based on the GENCODE human annotation version 24. AIDE achieves the best accuracy among the four methods.

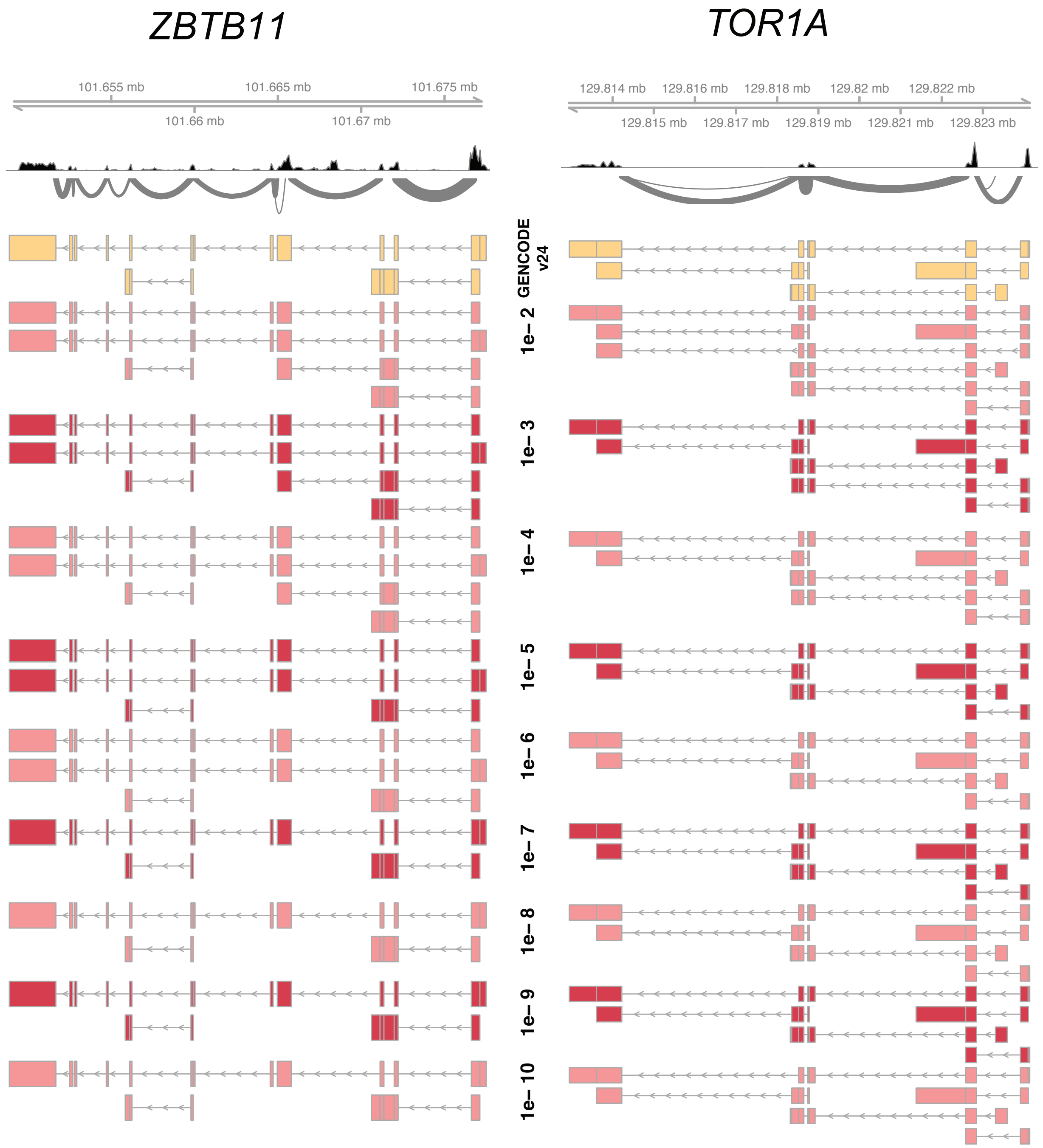


**Figure S10:** Isoform discovery for the human genes *ZBTB11* and *TOR1A* given different $p$-value thresholds. The histogram and the sashimi plot denote the RNA-seq reads mapped to the two genes in the human ESC sample 1 (Table S1). Isoform discovery is based on the GENCODE human annotation version 24. The threshold on the $p$-values resulted from the likelihood ratio tests decreases from $10^{-2}$ to $10^{-10}$.

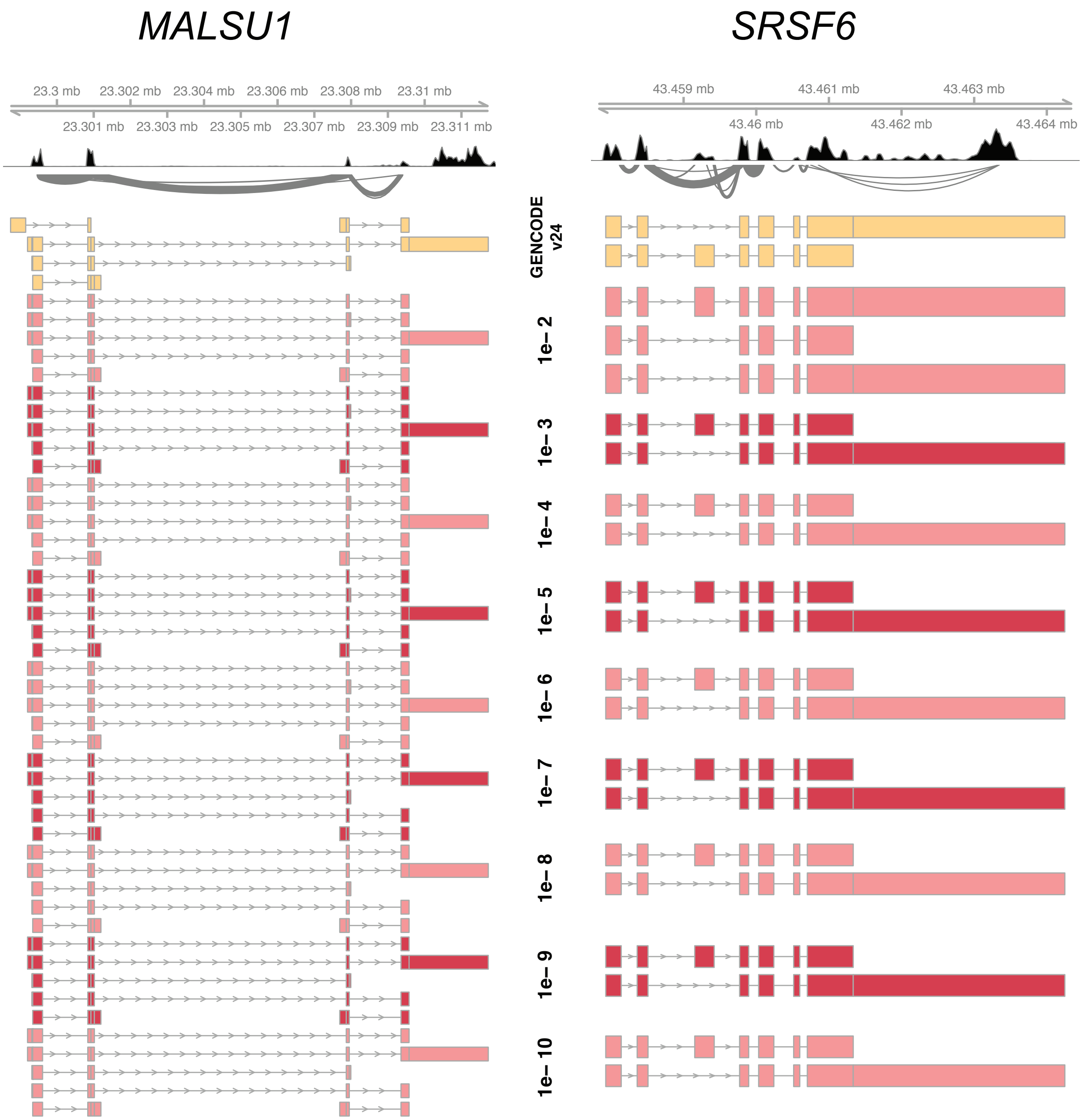


**Figure S11:** Isoform discovery for the human genes *MALSU1* and *SRSF6* given different $p$-value thresholds. The histogram and the sashimi plot denote the RNA-seq reads mapped to the two genes in the human ESC sample 1 (Table S1). Isoform discovery is based on the GENCODE human annotation version 24. The threshold on the $p$-values resulted from the likelihood ratio tests decreases from $10^{-2}$ to $10^{-10}$.

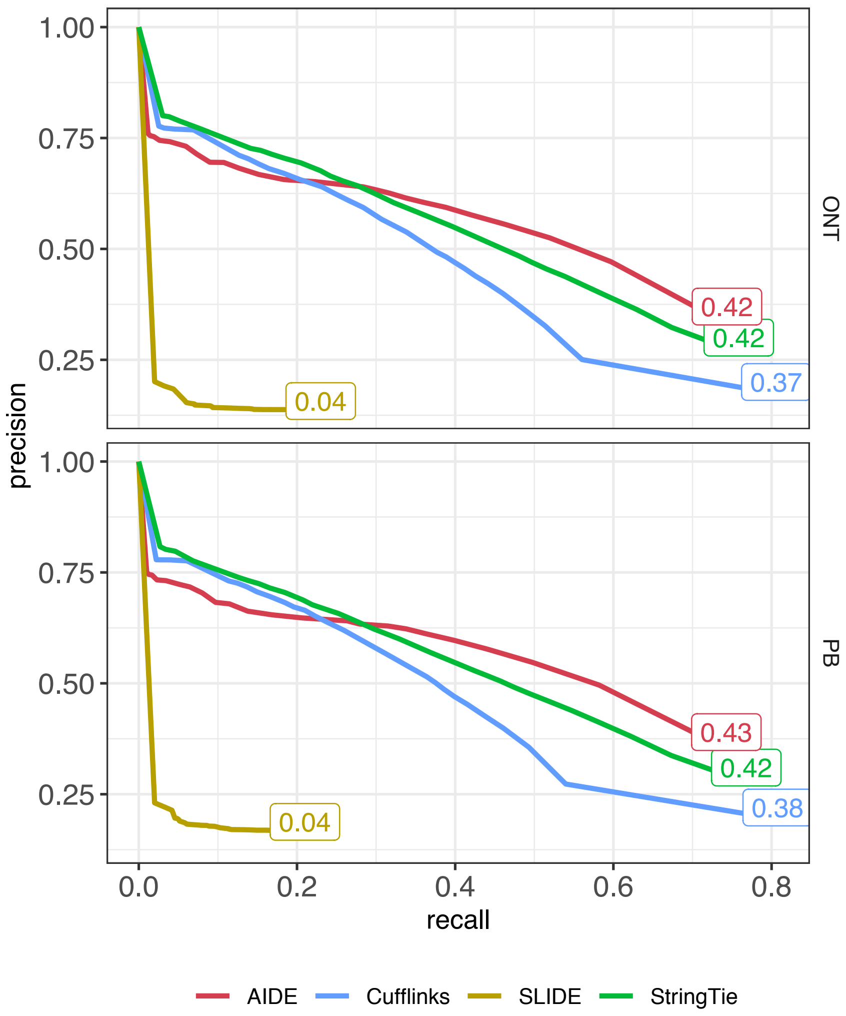


**Figure S12:** Comparison between AIDE and the other three isoform discovery methods based on long-read sequencing technologies. The estimated expression levels of the predicted isoforms were then summarized in the FPKM unit. The precision-recall curves (at transcript-level) were obtained by thresholding the FPKM values of the predicted isoforms. The corresponding AUC of each method is marked in the plot.

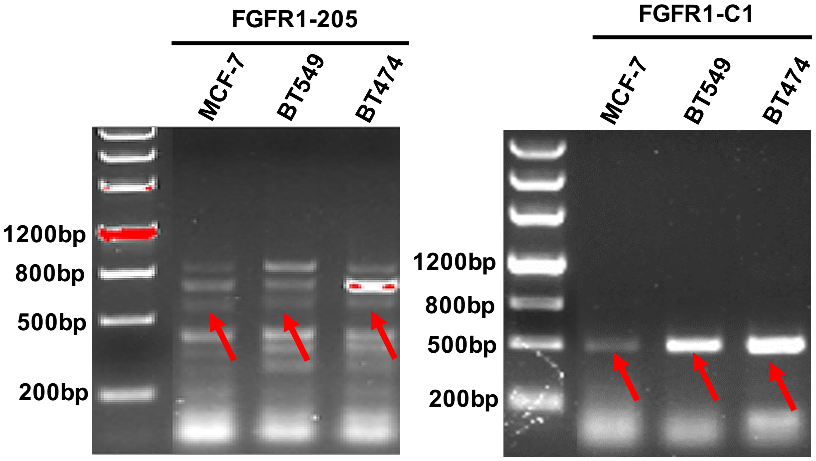


**Figure S13:** The expression of *FGFR1-205* and *FGFR1-C1* in breast cancer cell lines MCF7, BT474, and BT549 were validated by PCR.

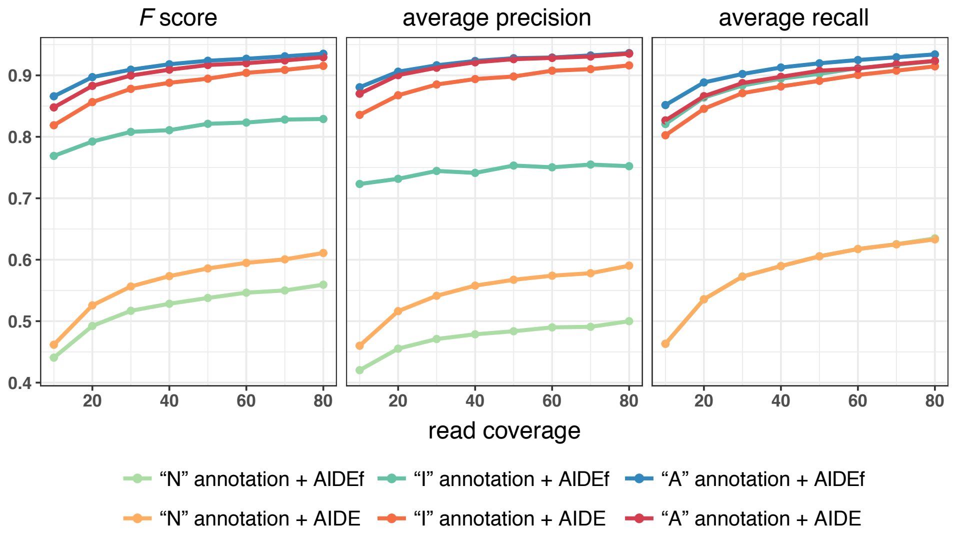


**Figure S14:** Comparison of AIDE (stepwise selection) and AIDEf (forward selection only) in terms of the average $F$ scores, precision rates, and recall rates (i.e., three measures) across the $2,262$ genes in simulation. For AIDE and AIDEf, each measure is calculated based on three types of annotations and RNA-seq samples with varying read coverages. The vertical axes of the three panels denote the values of the three measures, and the horizontal axis denotes the average per-base coverage of RNA-seq reads.

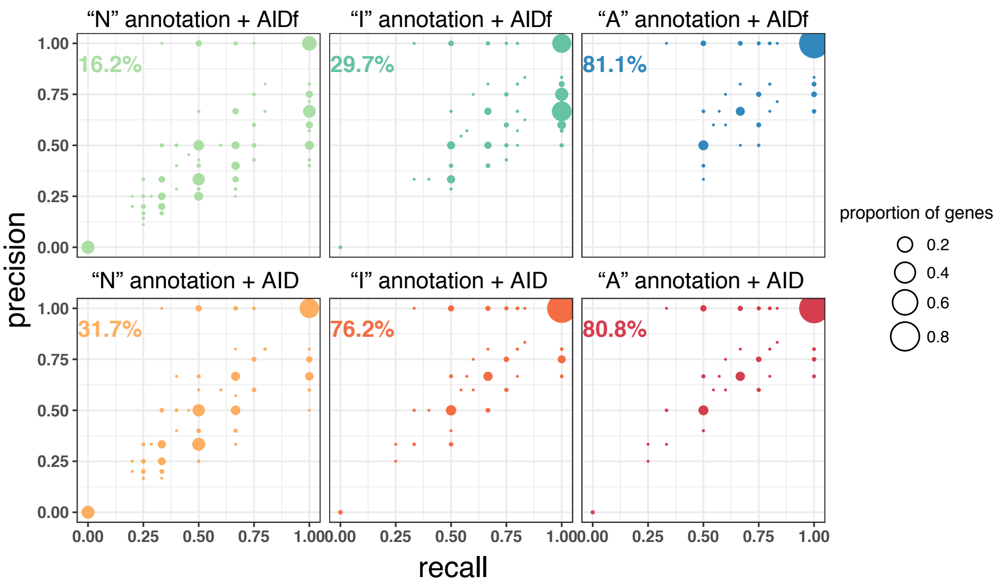


**Figure S15:** Comparison of AIDE (stepwise selection) and AIDEf (forward selection only) in terms of the per-gene precision and recall rates in simulation, with $80$x read coverage and three types of annotations. The circle sizes are proportional to the proportions of genes with the corresponding precision and recall rates, with the top-right circles indicating $100\%$ accuracy.

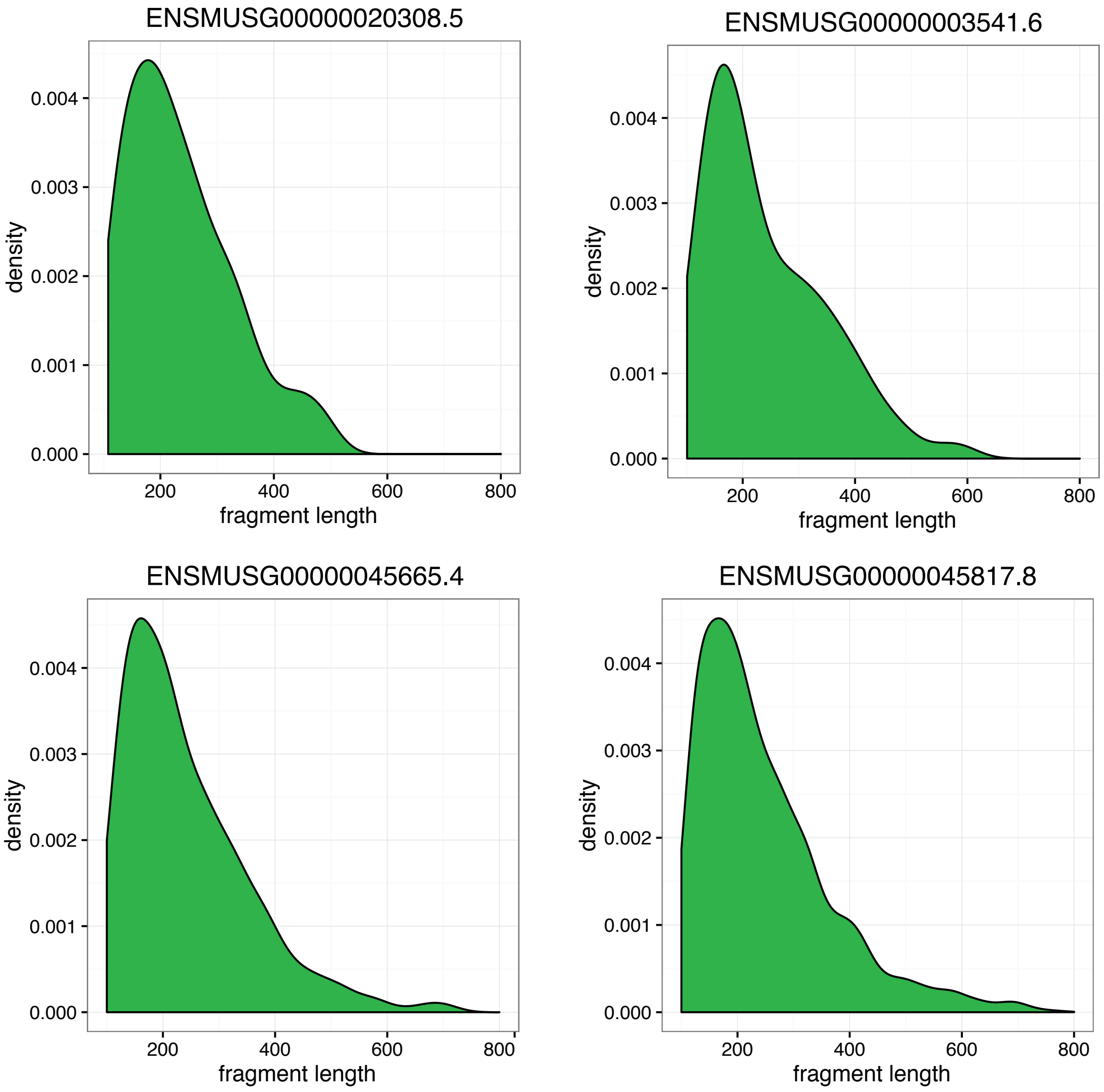


**Figure S16:** The distribution of fragment lengths in real RNA-seq data. The empirical fragment length distribution of four example mouse genes in the mouse bone marrow-derived macrophage dataset (Table S1).
