## Supplemental Figures for "AIDE: annotation-assisted isoform discovery with high precision"

### Supplemental Methods

#### Calculation of precision and recall rate of isoform reconstruction

We denote the true isoform set in an RNA-seq sample as $S$ and the discovered isoform set as $D$. We use $B(s)$ and $E(s)$ to denote the number of bases and the number of exons in isoform $s$, respectively. We calculate the isoform-level precision and recall rate of isoform discovery by comparing the two sets $S$ and $D$. When compare the isoform $s_1$ in $S$ and $s_2$ in $D$, we allow a small difference by requiring the number of mismatch bases to be smaller than $0.99B(s_1)$. The isoform-level precision and recall rates are calculated as

$$\text{precision}^{\text{isoform}} = \frac{|D \cap S|}{|D|}, \quad \text{recall}^{\text{isoform}} = \frac{|D \cap S|}{|S|}.$$

The exon-level precision and recall rates are calculated as

$$\text{precision}^{\text{exon}} = \frac{\sum_{s \in D \cap S} E(s)}{\sum_{s \in D} E(s)}, \quad \text{recall}^{\text{exon}} = \frac{\sum_{s \in D \cap S} E(s)}{\sum_{s \in S} E(s)}.$$

The base-level precision and recall rates are calculated as

$$\text{precision}^{\text{base}} = \frac{\sum_{s \in D \cap S} B(s)}{\sum_{s \in D} B(s)}, \quad \text{recall}^{\text{base}} = \frac{\sum_{s \in D \cap S} B(s)}{\sum_{s \in S} B(s)}.$$

#### Proof-of-concept simulation strategy

To evaluate the robustness of AIDE to the accuracy of annotation, we considered three types of annotation sets: (1) “N” (no) annotations: no annotated isoforms were used, and both AIDE and AIDEf were initialized with the candidate isoform compatible with the most reads and directly went into stage 2; (2) “I” (inaccurate) annotations: the “annotated isoforms” used in stage 1 consisted of half of the randomly selected true isoforms and the same number of false isoforms (not in the GENCODE annotation but randomly assembled from exons); (3) “A” (accurate) annotations: the “annotated isoforms” used in stage 1 (Figure 1) consisted of half of the randomly selected true isoforms. We generated RNA-seq data with different read coverages to compare AIDE and AIDEf with each of the three types of annotations, and we evaluated the performance of AIDE and AIDEf in terms of the precision rate, the recall rate, and the $F$ score.

We consider $2,262$ protein-coding genes from the human GENCODE annotation (version 24) [8]. The exon numbers of these genes range from $2$ to $10$. We treat the annotated isoforms in this GENCODE annotation as the true isoforms, randomly assign isoform abundance, and simulate paired-end RNA-seq reads from those isoforms with a $100$ bp read length. As we know the underlying true isoforms and their abundance in this simulation, we can use them to evaluate the accuracy of AIDE.

Suppose that gene $i$ has $J_i$ annotated isoforms in the GENCODE annotation. We assume these $J_i$ isoforms are the true isoforms, and we randomly select from them $\lceil \frac{J_i}{2} \rceil$ isoforms to be used in the "A" annotations or the "I" annotations. The isoform proportions of the true isoforms are simulated from a symmetric Dirichlet distribution with parameters $(1/\lceil \frac{J_i}{2} \rceil, \ldots, 1/\lceil \frac{J_i}{2} \rceil)'$. The total number of reads generated from gene $i$ depends on the gene length and the read coverage. Within each transcript, the starting positions of reads are uniformly distributed, and the fragment length follows a log-normal distribution with mean $180$ bp and standard deviation $65$ bp. Once we simulate the reads, we find the chromosome coordinates of the two ends of each read, and those coordinates are the only observed data input into AIDE and AIDEf.

#### Definition of subexons

As different isoforms of the same gene may consist of overlapping but not identical exons, we divide exons into subexons (see the figure below for an illustration), which are defined as transcribed regions between every two adjacent splicing sites in annotations, same as in SLIDE [18]. By this definition, every gene is composed of non-overlapping subexons and introns. For the ease of terminology, we refer to subexons as exons in our main text.

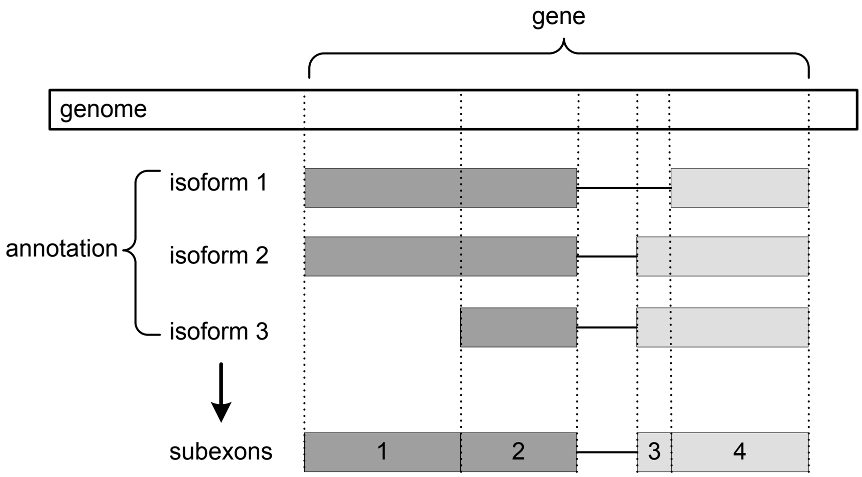


**Figure ST1:** Illustration of subexons. The example gene has two exons, represented by light and dark gray boxes, and three mRNA isoforms. The solid lines between exons represent introns in the gene that have been spliced out in isoforms. Adjacent splicing sites in these isoforms define four non-overlapping subexons: the first exon is divided into subexon 1 and 2, and the second exon is divided into subexon 3 and 4.

#### Transcriptome profiling by next-generation sequencing

Breast cancer biopsies were collected from Clinical Research Center for Breast of West China Hospital under the supervision of China Association for Ethical Studies. The whole-transcriptome RNA sequencing was performed with the next-generation sequencing standard protocol using the Illumina sequencer. Long RNA sequences were first converted into a library of cDNA fragments, and adaptors were subsequently added to each cDNA fragment. Illumina relies on the attachment of small DNA fragments to a platform, optically transparent surface, and solid-phase amplification to create an ultrahigh-density sequencing flow cell with $> 10$ million clusters, each containing $\sim 1,000$ copies of template/cm$^2$. These templates were sequenced by a robust four-color DNA sequencing-by-synthesis technology that employs reversible terminators with removable fluorescence. The high-sensitivity fluorescence was then detected by laser excitation. The typical output from a single reaction was approximately 2GB containing $20$-$40$ million reads. The Illumina-sequenced reads were assembled using the CLC bio software. Before the assembly, the reads were preprocessed by masking the polyA tails and removing the adapters.

#### Validation of transcripts by PCR and Sanger sequencing

Biopsy was collected freshly and the total RNA was extracted with the RiboPure Kit (Ambion). The reverse transcription reactions were performed using the RevertAid First-Strand cDNA Synthesis System kit (Thermo Fisher Scientific) . With the cDNAs as templates and primers from TSINGKE, the PCR procedure ($95$°C $5$min, $95$°C $30$s, $55$°C $30$s, $72$°C $1$min to $2$min, 40 to 50 cycles, $72$°C $5$min for extension) was conducted using a Thermo Fisher Scientific PCR system. PCR products were purified using Gel Extraction Kit (OMEGA) followed by Sanger sequencing with their special forward primers. Finally, sequencing result was analyzed using the Sequence Scanner Software.

#### Validation of isoform functions by colongenic assay

Breast cancer cell lines BT549, MB231, SUM149, BT474, SK-BR-3, and MCF-7 were from ATCC and cultured in the State Key Laboratory of Biotherapy. The cells' total RNA was extracted with the RiboPure Kit (Ambion), and the reverse transcription reactions were performed using the RevertAid First-Strand cDNA Synthesis System kit (Thermo Fisher Scientific). With the cDNAs as templates and primers from TSINGKE, the PCR procedure ($95$°C $5$min, $95$°C $30$s, $55$°C $30$s, $72$°C $1$min, 40 cycles, $72$°C $5$min for extension) was conducted using a Thermo Fisher Scientific PCR system. PCR products were purified using the Gel Extraction Kit (OMEGA) followed by Sanger sequencing with special forward primers. The five siRNAs specifically targeting *FGFR1-238* were synthesized at Shanghai GenePharma. Lipofactamine3000 from Invitrogen were used in siRNA transferring in breast cancer cells on the first day and the sixth day. The cells' colonegenic assays lasted for 10-12 days.
