## Supplemental Tables for "AIDE: annotation-assisted isoform discovery with high precision"

**Table S1:** Description of real RNA-seq data sets.

| sample | cell type | read length | accession number |
|---|---|---|---|
| 1 | HepG2 | 50×2 | ENCFF084JYA |
| 2 | HepG2 | 50×2 | ENCFF790CFB |
| 3 | HepG2 | 100×2 | ENCFF916YZY, ENCFF800YJR |
| 4 | HepG2 | 100×2 | ENCFF179TFY, ENCFF782TAX |
| 5 | HepG2 | 76×2 | ENCFF168NGI |
| 6 | HepG2 | 76×2 | ENCFF711DJN |
| sample | cell type | read length | accession number |
| 1 | human ESC | 76×2 | GSE90225 |
| 2 | human ESC | 76×2 | GSE33480 |
| 3 | human ESC | 101×2 | GSE47626 |
| sample | cell type | read length | accession number |
| 1 | mouse BMDM | 100×2 | ENCSR614DLJ |
| 2 | mouse BMDM | 100×2 | ENCSR822FMG |
| 3 | mouse BMDM | 100×2 | ENCSR614KOV |

**Table S2:** Experimental validation of identified transcripts.

| | Transcript | GENCODE | AIDE | cufflinks | Isoform was validated by Sanger |
|---|---|---|---|---|---|
| Category 1 | MTHFD2-203 | + | - | + | No |
| | NPC2-205 | + | - | + | No |
| | RBM7-208 | + | - | + | No |
| | CD164-210 | + | - | + | No |
| | XBP1-205 | + | - | + | Yes |
| | SYNGR2-new | - | - | + | Yes |
| Category 2 | FGFR1-238 | + | + | - | Yes |
| | ZFAND5-208 | + | + | - | Yes |
| | BRCA2-207 | + | + | - | No |
| | PDCD5-201 | + | + | - | No |

+, Isoform was predicted; -, isoform was NOT predicted

**Table S3:** Primers designed for PCR validation of identified transcripts.

| Primer name | sequence | Predicted size(bp) | PCR product size(bp) |
|---|---|---|---|
| MTHFD2-203-F | TTCTGGAAGGAAACTGGCCC | 366 | 750 |
| MTHFD2-203-R | ACCAACTTGGGTTTGGCAGT | | |
| NPC2-205-F | AGTGAATGTGAGCCCATGCC | 318 | 491 |
| NPC2-205-R | TCTGCTACAGAGCACCTCCT | | |
| RBM7-208-F | GAAGCGGATCGCACTCTCTT | 234 | 479 |
| RBM7-208-R | TGATCCAGAGGTGAAGAACCA | | |
| CD164-210-F | ATCTCCAACGTAACCTCGGC | 340 | 1032 |
| CD164-210-R | TGTAGTTCCTTGTGTGGCATCT | | |
| XBP1-205-F | CGACGGGACCCCTAAAGTTC | 389 | 389 |
| XBP1-205-R | AGGGGCTGAAACAACTTGGG | | |
| SYNGR2-new-F | CTTCCCACCCGAAGTTGAGG | 653 | 653 |
| SYNGR2-new-R | GAAGGCTCCAGAGGTGCTG | | |
| FGFR1-238-F | ACTGCAGAACTGGGATGTGG | 468 | 468 |
| FGFR1-238-R | CTACGGGCATACGGTTTGGT | | |
| ZFAND5-208-F | TCGGGGAAGGGTCGGATTAT | 486 | 486 |
| ZFAND5-208-R | CGGGGTCTGGTTAGTCTCCT | | |
| BRCA2-207-F | ACAAAGGCAACGCGTCTTTC | 427 | 419 |
| BRCA2-207-R | AGGCACATTCCATAGCTGCC | | |
| PDCD5-201-F | ACGAGGAGCTTGAGGCGCTGA | 408 | 413 |
| PDCD5-201-R | TAGACTTGTTCCGTTAAGTTC | | |
| FGFR1-238-E18-F | CAGAAACTGAAACCCAGACATGTG | 533 | 533 |
| FGFR1-238-E18-R | AATAGTCGCCAACACTGCAGCT | | |
| FGFR1-C1-F | AAGAGAGAGAAGGGGTTAGG | 528 | 528 |
| FGFR1-C1-R | TTCTTAAGTGAAGCACCTCC | | |
| FGFR1-205-F | TCTAACTGCAGAACTGGGATGTGG | 510 | 510 |
| FGFR1-205-R | AGTGGGCAGCAGTTTCTGAAGC | | |

**Table S4:** Specific siRNAs designed for isoforms *FGFR1-238*, *FGFR1-C1*, and *FGFR1-205*.

| FGFR1-238 siRNA | sense | antisense |
|---|---|---|
| 1 | GCGCAGGUUCCUUGUAACCUCUUCU | AGAAGAGGUUACAAGGAACCUGCGC |
| 2 | CCAUGGAUGGUUUCCUCCAAGGAAA | UUUCCUUGGAGGAAACCAUCCAUGG |
| 3 | CAAAUGAUGAAGGUCUGCAGAAACU | AGUUUCUGCAGACCUUCAUCAUUUG |
| 4 | UCUGCAGAAACUGAAACCCAGACAU | AUGUCUGGGUUUCAGUUUCUGCAGA |
| 5 | GCAAUGUUGUGUGAAGGGAUGAAGA | UCUUCAUCCCUUCACACAACAUUGC |
| **FGFR1-C1 siRNA** | GCUUCAGAAACUGCUGCCCACUAAC | GUUAGUGGGCAGCAGUUUCUGAAGC |
| **FGFR1-205 siRNA** | GGUCUCAUGUCCUGUGCUU | AAGCACAGGACAUGAGACC |
